# Molecular characterization revealed the role of thaumatin-like proteins in stress response in bread wheat

**DOI:** 10.1101/2020.09.24.311928

**Authors:** Alok Sharma, Himanshu Sharma, Ruchika Rajput, Ashutosh Pandey, Santosh Kumar Upadhyay

**Affiliations:** Department of Botany, Panjab University, Chandigarh, India-160014; I.K. Gujral Punjab Technical University, Kapurthala, Jalandhar, India-144603; National Institute of Plant Genome Research, Aruna Asaf Ali Marg, New Delhi, India-110067

**Keywords:** Abiotic stress, Biotic stress, Cereals, Thaumatin-like proteins, *TaTLP2-B*, *Triticum aestivum*

## Abstract

Thaumatin-like proteins (TLPs) are related to the pathogenesis-related-5 (PR-5) family and involved in stress response. Herein, a total of 93 *TLP* genes were identified in the genome of *Triticum aestivum.* Further, we identified 26, 27, 39 and 37 *TLP* genes in the *Brachypodium distachyon*, *Oryza sativa*, *Sorghum bicolor* and *Zea mays* genomes for comparative characterization, respectively. They could be grouped into small and long TLPs with conserved thaumatin signature motif. Tightly clustered genes exhibited conserved gene and protein structure. The physicochemical analyses suggested significant differences between small and long TLPs. Evolutionary analyses suggested the role of duplication events and purifying selection in the expansion of the TLP gene family. Expression analyses revealed the possible roles of *TLPs* in plant development and abiotic and fungal stress response. Recombinant expression of *TaTLP2-B* in *Saccharomyces cerevisiae* provided significant tolerance against cold, heat, drought and salt stresses. The results depicted the importance of TLPs in cereal crops that would be highly useful in future crop improvement programs.

## 1. Introduction

The growth and development of plants have always been affected by various abiotic and biotic stress conditions. In response to these stress conditions, plants produce numerous molecules including pathogenesis-related (PR) proteins [1, 2]. PR proteins belong to a superfamily of defence-related proteins consisting of PR1-PR17 protein families, that have been classified based on amino acid sequences, serological reaction, enzymatic activity etc. [3]

Thaumatin-like proteins (TLPs) are related to the PR5 family of the PR superfamily [3]. These are called TLPs due to their significant similarity with a ∼23 kDa sweet-tasting protein known as thaumatin [4]. Thaumatin protein was first isolated from an African shrub known as *Thaumatococcus danielli* [5]. Over the period, TLPs have been reported in a diverse group of organisms including insects, nematodes, fungi and plants [6, 7]. In plants, TLPs are reported from algae to angiosperms [6–9].

Based on the molecular weight (MW), TLP proteins are grouped as long (L-type) and small (S-type) TLPs, having the MW ∼21-26 kDa and ∼16-17 kDa, respectively. Long TLPs have been identified from each group of plants, while small TLPs are confined to only gymnospermous and monocot plants [6, 9]. A total of 16 and 10 conserved cysteine residues, forming eight and five disulfide linkages, are known to be present in long and small TLPs, respectively. These disulfide bonds provide resistance against extreme pH, heat and protease degradation [6, 10]. Each TLP consists of a conserved REDDD motif and a thaumatin signature motif G-X-[GF]-X-C-X-T-[GA]-D-C-X-(1,2)-G-X-(2,3)-C. The REDDD motif is involved in receptor binding for their antifungal action [6]. Numerous studies reported that the overexpression of TLPs provides significant resistance against various fungi in both dicot and monocot plants [11–13]. The mechanism of action of TLPs in fungal resistance is ambiguous; however, it has been presumed that they work by degradation and permeabilization of the fungal cell walls [6, 14]. Besides, TLPs are also known to be involved in antifreeze action and abiotic stress resistance in plants [15]. Further, they are also reported to be involved in flowering, fruit ripening and seed germination [16–18].

Despite utmost importance, a detailed characterization of TLPs is still limited in various cereal crops. Therefore, the current study aims at an inclusive characterization of TLPs in bread wheat, an important cereal crop. Besides, we have also performed a comparative analysis of various features of TLPs in other cereals including *B. distachyon*, *O. sativa*, *S. bicolor* and *Z. mays*. Chromosomal distribution, phylogeny, physicochemical properties, gene duplication events, *cis*-regulatory elements, and gene and protein structural analyses were performed. To decipher their roles in plant development and under abiotic and biotic stresses, their expression analysis was performed using high throughput RNA-seq data. The expression of eight selected *TaTLP* genes was validated using qRT-PCR. An abiotic stress-responsive gene, *TaTLP2-B*, of *T. aestivum* was cloned and used for functional characterization in *Saccharomyces cerevisiae* (yeast) cells. Recombinant expression of TaTLP2-B provided significant tolerance to the yeast in spot assay against various abiotic stresses. The study will increase the knowledge about numerous TLP proteins and provide the base for the functional characterization of identified genes in future studies.

## 2. Material and methods

### 2.1. Plant materials, growth conditions and stress treatments

Surface sterilization of bread wheat seeds (cv. Chinese Spring) was done with 1.2% sodium hypochlorite in 10% ethanol followed by the washing with double autoclaved water. The sterilized seeds were kept overnight on moist Whatman filter papers at 4^D^C for stratification and were further kept at room temperature for germination. The seedlings were then transferred to fresh phytaboxes and were allowed to grow in plant growth chambers under 60% relative humidity, 22^D^C temperature and a 16 and 8 h light and dark period, respectively [19]. Seven-days old seedlings were kept under heat (40^D^C), drought (20% PEG) and combined heat and drought stress treatments for 6, 12, 24 and 48 hours (h). For salinity stress, plants were subjected to 150 mM NaCl for 6, 12, 24 and 48 h. The plant samples including roots and shoots were collected and stored at −80^D^C till further experiments.

### 2.2. Identification and nomenclature of the TLP genes

Extensive BLAST searches were used to identify the TLP proteins in five different cereals including *B. distachyon*, *O. sativa*, *S. bicolor*, *T. aestivum* and *Z. mays*. Arabidopsis TLP sequences were used as a query against the protein model sequences of each crop downloaded from the Ensembl Plants [9, 20]. The presence of the thaumatin (PF00314) domain was analyzed using the Hidden Markov Model (HMM) and Pfam BLAST searches at e-value 10^-10^. Identified TLP sequences were further subjected to the NCBI Conserved Domain Database (CDD) BLAST to further confirm the presence of the thaumatin domain. The *TLP* genes, having a complete thaumatin family signature, were selected for further analysis. Identified *TLPs* were named as per their sequence of occurrence at various chromosomes in each crop, except *T. aestivum*. The international rules for gene symbolization of *T. aestivum* (http://wheat.pw.usda.gov/ggpages/wgc/98/Intro.htm) were followed for the nomenclature of *TaTLPs*.

### 2.3. Chromosomal localization and duplication events

Plant Ensemble (http://plants.ensembl.org/) was used for gathering the chromosomal and sub-genomic location of *TLPs* of all five crops. The homeologous *TLP* genes in *T. aestivum* were ≥90% sequence similarity and their occurrence at the related chromosomes. The MapInspect software was used for the graphical representation of *TLP* genes on their respective chromosomes (http://www.plantbreeding.wur.nl/uk/software_mapinspect.html.2012). Duplication events (DEs) were identified using a bidirectional blast hit approach with sequence ≥80%, while tandem and segmental DEs were segregated based on their distance and occurrence at respective chromosomes as per the previous studies [21].

### 2.4. Multiple sequence alignments, phylogeny and structural analysis

The Muscle and Multalin tools were used for the multiple sequence alignments of all the TLP*s* with a known thaumatin protein (P02883.2|THM1_THADA) to find the conserved residues [22, 23]. The phylogenetic tree was constructed by the maximum likelihood method with 1000 bootstraps using the MEGA X software [24].

### 2.5. Ka/Ks and Tajima’s relativity test

The alignment of the protein and nucleotide sequences of the paralogous gene pairs was done using the ClustalOmega server (https://www.ebi.ac.uk/Tools/msa/clustalo). The synonymous substitution per synonymous site (Ks), the non-synonymous substitution per non-synonymous site (Ka), and their ratio (Ka/Ks) were calculated using the PAL2NAL programme [25, 26]. The calculation of the divergence time of each pair of duplicated genes was done using the formula T=Ks/2r, where T represents the divergence time and r represents the divergence rate. The divergence rate was assumed to be 6.5 × 10^-9^ for cereals [27]. The Tajima’s relativity test was performed to find out the evolutionary rate between paralogous genes [28].

### 2.6. Gene structure organization

The genomic and coding DNA sequence (CDS) of identified *TLP* genes were used for the analysis of gene structure in terms of exon-intron organization and intron phases using the GSDS 2.0 server as done in earlier studies [29, 30].

### 2.7. Physicochemical properties of the TLPs

Various physicochemical characteristics such as peptide length, molecular weight (MW) and isoelectric point (pI) were analyzed using the ExPasy tool, which were further confirmed from the Ensemble plants and Sequence Manipulation Suite [31–33]. Tools including CELLO v.2.5, ngLOC, ProtComp9 and WoLF PSORT were used for the prediction of subcellular localization of TLP proteins [34–37]. Tools such as Phobius and DAS-Tmfilter were used for the prediction of transmembrane regions [38, 39]. The signal peptides were detected using the tools Phobius and SignalP [38, 40]. SMART server was used for the domain analysis, whereas, motifs were analyzed using MEME v.4.11.4 [41, 42].

### 2.8. Cis-regulatory element analysis

For *cis-*regulatory elements analysis, 1.5 kb upstream genomic sequences from the initiation codon were retrieved for each *TLP* gene. These promoter sequences were analyzed using the PLACE software [43]. Identified *cis*-regulatory elements were categorized based on their functions.

### 2.9. Expression profiling using RNA-seq data

Expression analysis in various tissues of each crop was done using the high throughput RNA-seq data retrieved from the URGI database (wheaturgi.versailles.inra.fr/files/RNASeqWheat/) and Expression ATLAS [44–46]. In *T. aestivum*, the expression data generated in replicates for various tissue developmental stages, under biotic (fungal pathogen) and abiotic (heat, drought and salt) conditions were used [47–49]. Data generated for three developmental stages of root, stem, leaf, spike and grain was used to analyze the tissue-specific expression in wheat. Further, the RNA-seq data (PRJNA243835) generated in triplicates after the 24, 48 and 72 h of inoculation of *Blumeria graminis* f. sp. *tritici* (Bgt) and *Puccinia striiformis* f. sp. *tritici* (Pst) in 7-d-old seedlings were used for the *TaTLP* expression analysis under biotic stress [47]. Under abiotic stress conditions, duplicate RNA-seq data (SRP045409) for 1 and 6 h of treatments of heat (40° C), drought (20% PEG 6000) and a combination of both heat and drought stresses were used to study the *TaTLP* expression [48]. Root RNA-seq data (SRP062745) available in triplicates for the treatment of 150 mM NaCl at 6, 12, 24 and 48 h treatment were used to analyze the expression of *TaTLPs* under salt stress [49]. Trinity package was used to calculate the expression value in terms of FPKM value [50]. Hierarchical Clustering Explorer 3.5 was used to generate heat maps of differentially expressed genes, which were clustered using the Euclidean distance method [51].

### 2.11. RNA isolation, cDNA synthesis and qRT-PCR

The total RNA of each sample was isolated using the Spectrum™ Plant Total RNA kit (Sigma, USA). Seedlings (shoot and root) grown under normal conditions were used as a control. TURBO DNA-free™ Kit (Invitrogen, USA) was used to remove the genomic DNA contamination. The qualitative and quantitative analysis of RNA was done using agarose gel electrophoresis and nanodrop quantification. The Superscript III First-Strand Synthesis Supermix (Invitrogen, USA) was used to synthesize the cDNA from one microgram of total RNA. A real-time qRT-PCR was performed with the gene-specific primers of selected genes using SYBR Green at the 7900 HT Fast Real-Time PCR System (Applied Biosystems) following the method established in our laboratory [21]. The fold expression change was calculated using the delta-delta CT method (2^−ΔΔ^) using the expression of ADP-ribosylation factor (*TaARF*) as an internal control [52]. All the experiments were carried out in triplicates and expressed as mean ± SD. A significant difference between the control and treatments was examined by using the two-tailed student’s t-test.

### 2.12. Cloning and functional characterization

The full-length open reading frame (ORF) of the *TaTLP2-B* gene was amplified from cDNA using the *TaTLP2-B* forward (5’ GTAATGGCTCTTCTTCCTCCTCTGCTTCTG 3’) and *TaTLP2-B* reverse (5’ AATCTGGGCCACACGATCGCCCC 3’) primers. The amplified DNA was cloned into the pJET1.2 cloning vector and sequenced. *TaTLP2-B* gene was re-amplified from the confirmed clone using the same primers and ligated into the pYES2.1/V5-His-Topo vector (Invitrogen, USA). The recombinant plasmid (pYES2.1-*TaTLP2-B*) was transformed into *S. cerevisiae* (W303) (yeast) cells for further characterization. For control, *lacz* gene containing pYES2.1 vector (pYES2.1/V5-His/lacZ) was used during the studies. Spot assays using recombinant yeast cells were performed under various abiotic stress conditions. Recombinant yeast cells were grown overnight in SD/−ura (with 2% dextrose) medium at 30^°^C and 200 rpm as primary culture and further inoculated for secondary culture (1:100 dilutions). As soon as OD_600_ reached 0.4, 2% galactose was added to induce the expression of recombinant protein in yeast and kept at 30^°^C and 200 rpm for 6 h. The OD_600_ was adjusted to 0.4 and an equal volume (500 μl) of each induced culture was further diluted in 10 ml medium. The diluted cultures were treated with heat (37°C), cold (4°C), drought (30% PEG), combined heat-drought and salt (1 M NaCl) stresses for 24 h, separately. After the treatments, serial dilutions (10^0^, 10^-2^, 10^-4^ and 10^-6^) were prepared, 5 μl of each dilution was spotted on SD/-ura agar plates, and incubated at 30^°^C for 2–3 d [21]. All the experiments were performed in triplicates and results were compared visually.

### 2.13. Subcellular localization of the TaTLP2-B

Subcellular localization of TaTLP2-B was analyzed using CaMV35S-driven C-terminal YFP fusion construct generated by Gateway LR-recombination into binary vector PEG101. Plasmids were then transformed into *Agrobacterium tumefaciens* GV3101 strain. The cells were resuspended in a freshly prepared infiltration medium (10 mM MgCl^2^, 10 mM MES/KOH, pH 5.7 and 150 μM acetosyringone). The resulting constructs were infiltrated onto the abaxial surface of *Nicotiana benthamiana* leaf and kept at 22°C for 48 h. The YFP fluorescence was observed using a Leica TCS SP8 (Leica Microsystems, Wetzlar, Germany) laser-scanning confocal microscope at 514-527 nm.

## 3. Results

### 3.1. Identification and chromosomal localization of the TLP genes

Numerous properties of TLPs ascribed to defence and development pathways and lack of such studies in numerous cereals lead us to perform a comprehensive analysis of the TLP family in five major cereals. An extensive BLAST search identified 26, 27, 39, 93 and 37 *TLP* genes in *B*. *distachyon, O*. *sativa, S*. *bicolor, T*. *aestivum* and *Z*. *mays*, respectively. The genes lacking or having incomplete thaumatin signature motifs were excluded from the study. Four *TLPs* were identified as small *TLPs* (sTLPs) in each *B. distachyon, O. sativa* and *Z. mays*, while 10 and 20 *sTLPs* were detected in *S. bicolor* and *T. aestivum,* respectively, (**supplementary file S1**). In the case of *T. aestivum*, identified *TLP* genes from the A, B and D sub-genomes formed 32 homeologous groups based on their sequence homology (≥90%). All of the identified proteins exhibited a complete thaumatin signature motif (**supplementary fig. S1)**.

The chromosomal localization suggested the scattered distribution of the *TLP* genes at the majority of chromosomes in each crop (Fig. 1). In *B. distachyon,* chromosome 4 consisted of a maximum of 10 *TLP* (both long and small) genes, while in the case of *O. sativa, S. bicolor, T. aestivum* and *Z. mays*, the majority of genes (6, 11, 18 and 11) were localized onchromosomes 12, 8, 5A and 1, respectively.

**Fig. 1.**
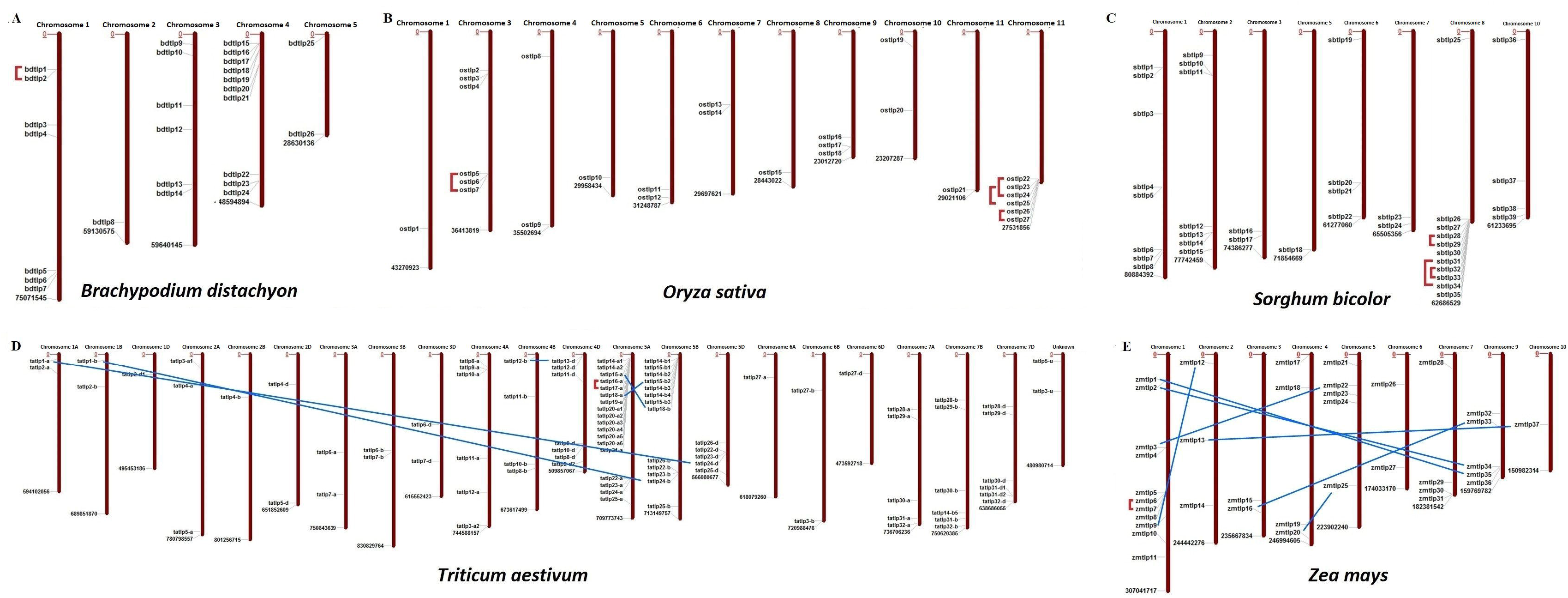
Chromosomal distribution and paralogous gene analysis of *TLPs* in five cereal crops. Image depicts the scattered localization of *TLP* genes of **(A)** *B. distachyon*, **(B)** *O. sativa*, **(C)** *S. bicolor*, **(D)** *T. aestivum* and **(E)** *Z. mays* at their respective chromosomes. Red lines and blue lines denote the tandem and segmental duplication events, respectively.

### 3.2. Multiple sequence alignments

TLP proteins consist of various characteristic and conserved residues, which are important for their functional activities. Multiple sequence alignments of TLP proteins with a known thaumatin protein (P02883.2|THM1_THADA) revealed the identification of 16 and 10 conserved cysteine residues in the majority of long and small TLP proteins of the studied cereal crops, respectively. Moreover, 7-15 cysteine residues had also been observed in a few TLPs of *O. sativa, S. bicolor, T. aestivum* and *Z. mays*. The REDDD motif was also found conserved in the majority of TLP proteins. However, in certain TLP proteins, a few amino acid residues (AAs) were replaced by other AAs having either similar or different properties. For instance, Arginine (R) was replaced by Lysine (K) or Asparagine (N). This could be responsible for the differential evolvement of various characteristic features of TLPs during evolution. The thaumatin signature motif, which is a characteristic feature of TLPs [6], was found to be highly conserved in all the identified TLP proteins. Some other regions like FF hydrophobic motif and bottom of acidic cleft forming amino acids were also found well conserved (Fig. 2, **supplementary fig. S1**).

**Fig. 2.**
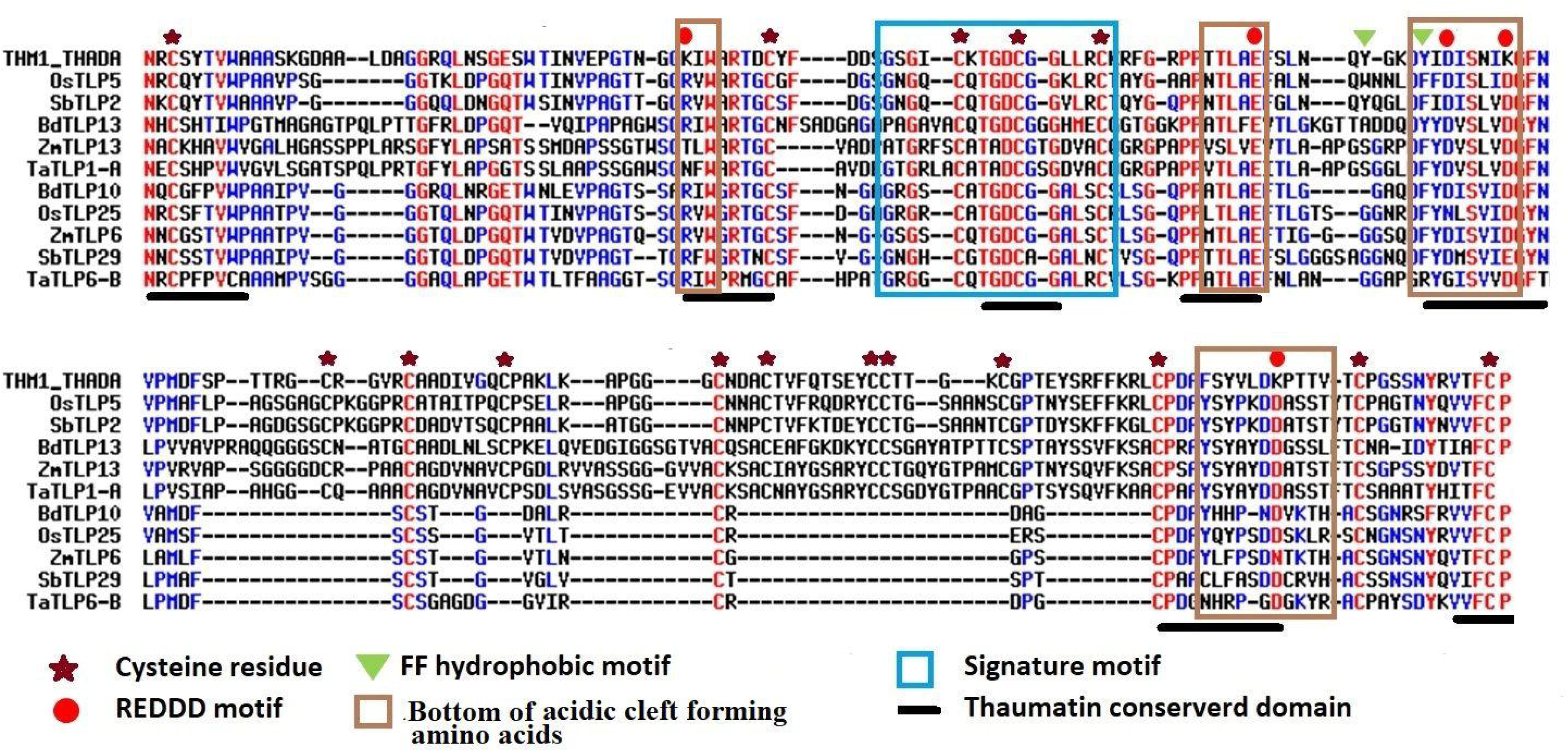
Multiple sequence alignment of selected long and small TLPs of *B. distachyon, O. sativa,* S. *bicolor, T. aestivum* and *Z. mays* with a known thaumatin protein sequence. Figure shows the alignment of long (OsTLP5, SbTLP2, BdTLP13, ZmTLP13 and TaTLP1-A) and small (BdTLP10, OsTLP25, ZmTLP6, SbTLP29 and TaTLP6-B) TLPs with a known thaumatin (P02883.2|THM1_THADA) protein sequence. The characteristic thaumatin signature motif is shown in the sky blue box. All the conserved cysteine residues were marked with an asterisk. The REDDD motif was marked with a red circle. The FF hydrophobic motif and bottom of acidic cleft forming regions are marked with a green triangle and brown box, respectively. The conserved regions of the thaumatin domain are marked with a black line.

### 3.3. Evolutionary analyses

#### 3.3.1. Phylogeny

To decipher the evolutionary relationship, a phylogenetic tree was built using the full-length TLP protein sequences of the five cereals and *A. thaliana*. These were clustered into 11 different clades based on their phylogenetic relatedness, named as groups I to XI (Fig. 3). The highest number of TLPs were found in group XI, followed by group II and group X, while group IV was the smallest with only five genes. All the sTLPs were tightly clustered into group XI, which could be due to their smaller size. Further, the majority of groups consisted of TLPs from all the five cereals, except groups III, V and VI that lacked members from one or more plant species. Besides, group IV comprised only three TaTLP and two AtTLP proteins. Besides, the homeologous TaTLPs of *T. aestivum* were tightly clustered in proximity.

**Fig. 3.**
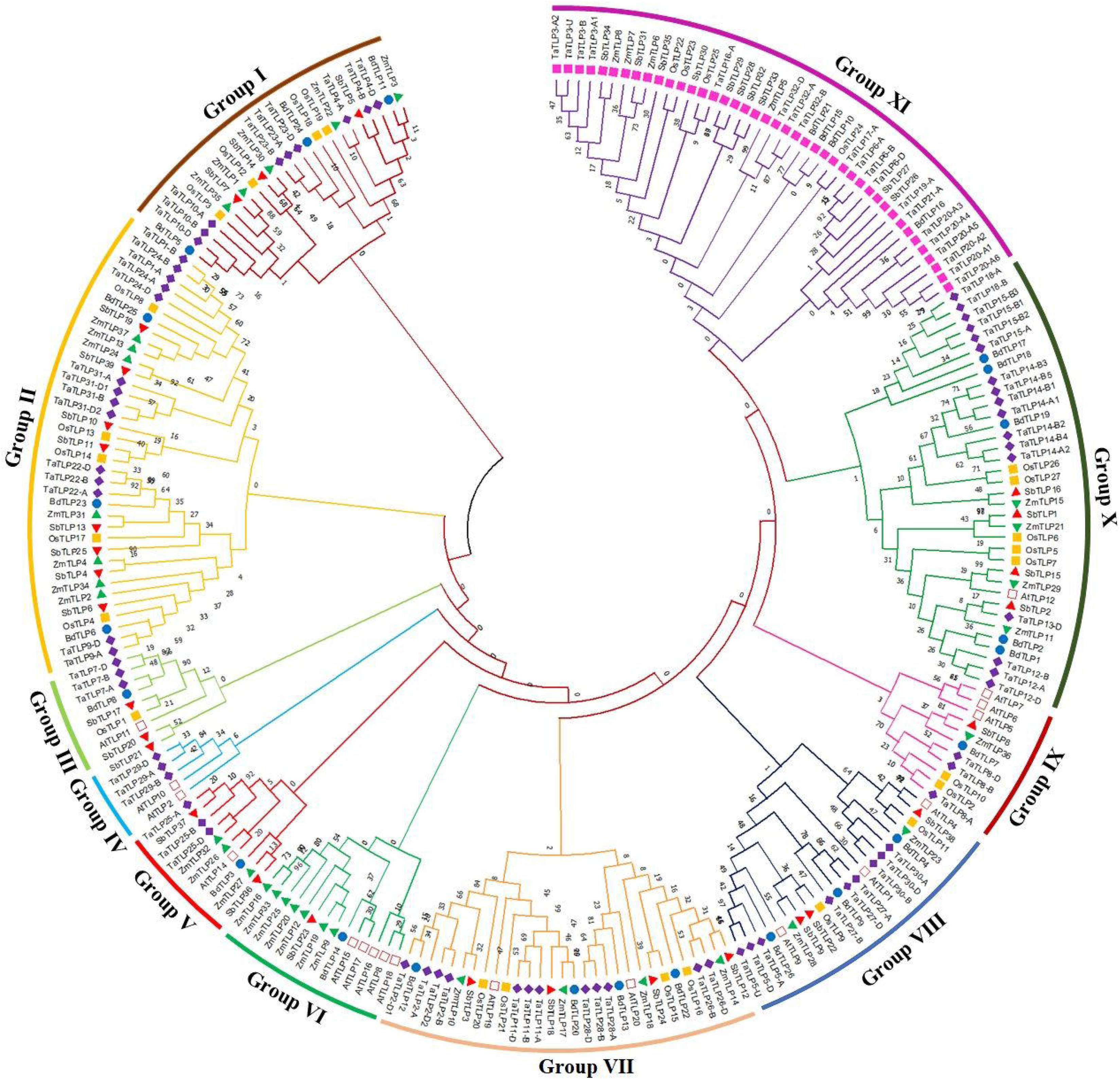
Phylogenetic tree analysis of *A. thaliana, B. distachyon, O. sativa. S, bicolor, T. aestivum and Z. mays* using full-length peptide of TLPs. The figure shows the phylogenetic tree of TLPs, built by the maximum likelihood approach with 1000 bootstrap values using MEGAX software. The phylogenetic tree displays the clustering of TLPs of six plant species in I-XI different groups, each group is coloured differently. The TLPs from each plant are marked with a different colour.

#### 3.3.2. Duplication events investigation

Duplication event (DE) analysis was carried out to understand their roles in the evolution and expansion of the TLP gene family in the studied crop species. A total of one, four, three, six and eight DEs were predicted in *B. distachyon, O. sativa, S. bicolor*, *T. aestivum* and *Z mays*, respectively (Fig. 1, **supplementary file S2**). Only tandem duplication events (TDEs) were found in the case of paralogous genes of *B. distachyon* (*BdTLP1-BdTLP2*)*, O. sativa* (*OsTLP5-OsTLP7, OsTLP22-OsTLP24, etc.*) and *S. bicolor* (*SbTLP28-SbTLP29, SbTLP31-SbTLP34, etc.*). However, in the case of *T. aestivum*, five DEs (*TaTLP1-A-TaTLP24-D, TaTLP1-B-TaTLP24-B, etc.*) were segmental and one was TDE (*TaTLP16-A-TaTLP17-A*). In *Z. mays*, seven segmental (*ZmTLP1-ZmTLP35, ZmTLP2-ZmTLP34* etc.) and one TDE (*ZmTLP6-ZmTLP7*) were found. All the duplicated gene pairs of each cereal crop were tightly clustered in proximity in the phylogenetic tree.

#### 3.3.3. Ka/Ks and Tajima’s relative rate test

Over the course of evolution, various evolutionary forces and natural pressures affected the duplicated genes [53]. To understand the evolutionary divergence between the paralogous gene pairs, the Ka/Ks analysis was carried out. The Ka/Ks ratio of more than one suggests the positive (non-purifying) and less than one indicates negative (purifying) selection pressure. All the paralogous genes showed the Ka/Ks value lesser than one, which suggested the negative or purifying selection on duplicated *TLP* genes (Table 1). However, Ka/Ks analysis for *ZmTLP6* and *ZmTLP7* was not performed due to their 100% similarity. Additionally, the divergence time of DEs was also calculated using the Ks value and previously described methods [54, 55]. The divergence time of duplicated genes was calculated as 83 million years ago (MYA) in *BdTLPs*, while the range varied from 52-108, 6-137, 63-99 and 1-63 MYA in *O. sativa, S. bicolor, T. aestivum* and *Z. mays*, respectively (Table 1). Tajima’s relative rate test found the insignificant χ^2^ value for all the duplicated gene pairs at P > 0.05 (Table 2). The P-value of more than 0.05 depicted their acceptance of the molecular evolutionary clock hypothesis [28].

**Table 1.**
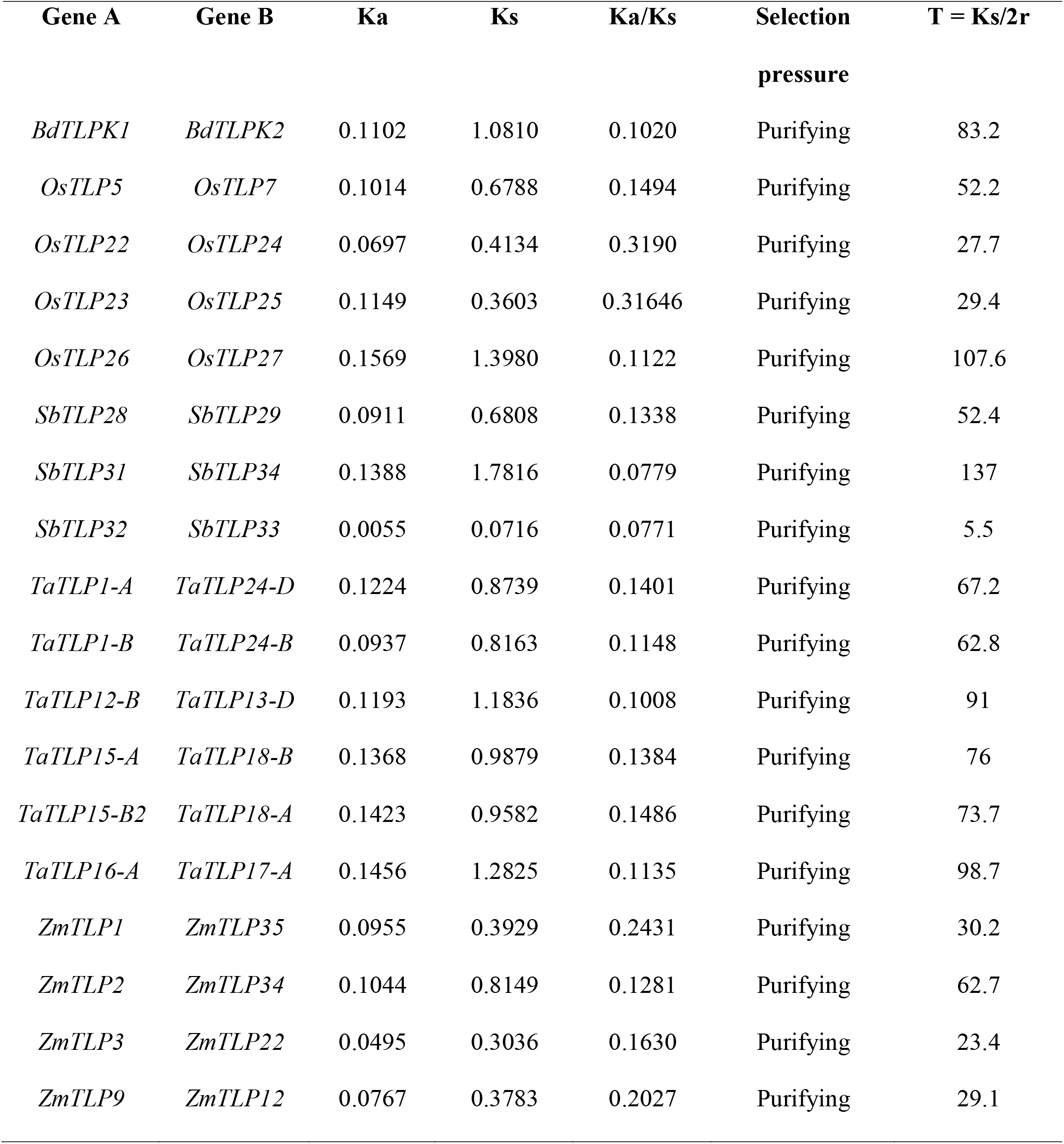
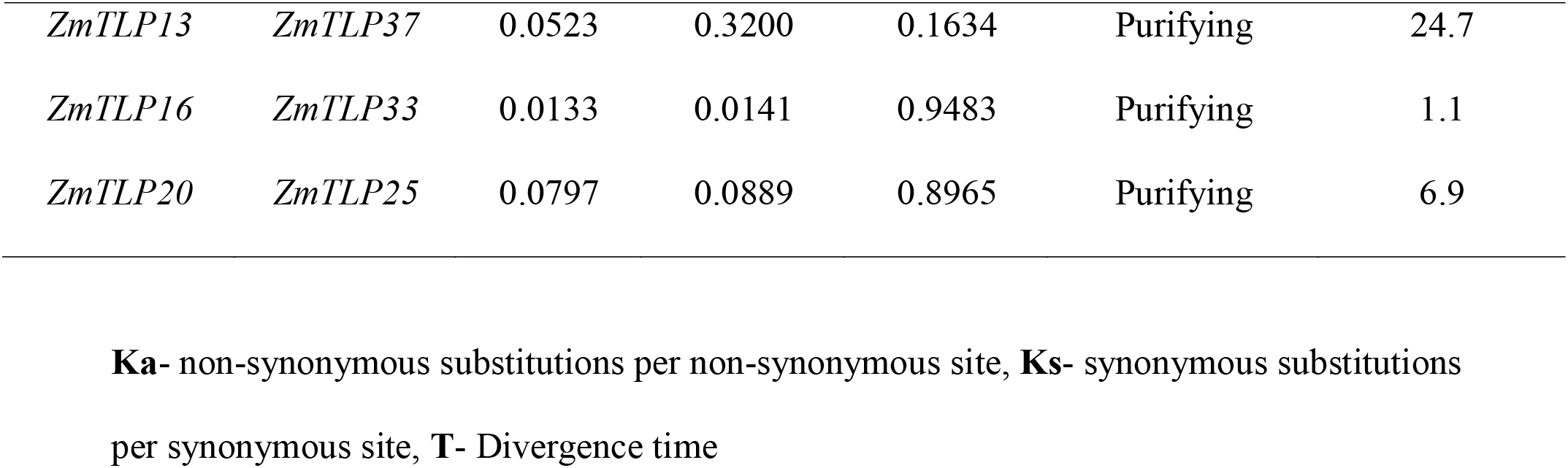
The Ka/Ks ratio and divergence time of duplicated *TLP* gene pairs.

| **Gene A** | **Gene B** | **Ka** | **Ks** | **Ka/Ks** | **Selection**<br>**pressure** | **T = Ks/2r** |
| --- | --- | --- | --- | --- | --- | --- |
| *BdTLPK1* | *BdTLPK2* | 0.1102 | 1.0810 | 0.1020 | Purifying | 83.2 |
| *OsTLP5* | *OsTLP7* | 0.1014 | 0.6788 | 0.1494 | Purifying | 52.2 |
| *OsTLP22* | *OsTLP24* | 0.0697 | 0.4134 | 0.3190 | Purifying | 27.7 |
| *OsTLP23* | *OsTLP25* | 0.1149 | 0.3603 | 0.31646 | Purifying | 29.4 |
| *OsTLP26* | *OsTLP27* | 0.1569 | 1.3980 | 0.1122 | Purifying | 107.6 |
| *SbTLP28* | *SbTLP29* | 0.0911 | 0.6808 | 0.1338 | Purifying | 52.4 |
| *SbTLP31* | *SbTLP34* | 0.1388 | 1.7816 | 0.0779 | Purifying | 137 |
| *SbTLP32* | *SbTLP33* | 0.0055 | 0.0716 | 0.0771 | Purifying | 5.5 |
| *TaTLP1-A* | *TaTLP24-D* | 0.1224 | 0.8739 | 0.1401 | Purifying | 67.2 |
| *TaTLP1-B* | *TaTLP24-B* | 0.0937 | 0.8163 | 0.1148 | Purifying | 62.8 |
| *TaTLP12-B* | *TaTLP13-D* | 0.1193 | 1.1836 | 0.1008 | Purifying | 91 |
| *TaTLP15-A* | *TaTLP18-B* | 0.1368 | 0.9879 | 0.1384 | Purifying | 76 |
| *TaTLP15-B2* | *TaTLP18-A* | 0.1423 | 0.9582 | 0.1486 | Purifying | 73.7 |
| *TaTLP16-A* | *TaTLP17-A* | 0.1456 | 1.2825 | 0.1135 | Purifying | 98.7 |
| *ZmTLP1* | *ZmTLP35* | 0.0955 | 0.3929 | 0.2431 | Purifying | 30.2 |
| *ZmTLP2* | *ZmTLP34* | 0.1044 | 0.8149 | 0.1281 | Purifying | 62.7 |
| *ZmTLP3* | *ZmTLP22* | 0.0495 | 0.3036 | 0.1630 | Purifying | 23.4 |
| *ZmTLP9* | *ZmTLP12* | 0.0767 | 0.3783 | 0.2027 | Purifying | 29.1 |
| *ZmTLP13* | *ZmTLP37* | 0.0523 | 0.3200 | 0.1634 | Purifying | 24.7 |
| *ZmTLP16* | *ZmTLP33* | 0.0133 | 0.0141 | 0.9483 | Purifying | 1.1 |
| *ZmTLP20* | *ZmTLP25* | 0.0797 | 0.0889 | 0.8965 | Purifying | 6.9 |

**Ka**- non-synonymous substitutions per non-synonymous site, **Ks**- synonymous substitutions
per synonymous site, **T**- Divergence time

**Table 2.** Tajima’s relative rate test of paralogous genes.

| **Group A** | **Group B** | **Outgroup** | **Nt** | **Na** | **Nb** | $\chi^2$ | **P** |
| --- | --- | --- | --- | --- | --- | --- | --- |
| *BdTLP1* | *BdTLP2* | *BdTLP11* | 403 | 47 | 42 | 0.28 | 0.59611 |
| *OsTLP5* | *OsTLP7* | *OsTLP18* | 401 | 28 | 29 | 0.02 | 0.89463 |
| *OsTLP22* | *OsTLP24* | *OsTLP18* | 333 | 15 | 17 | 0.13 | 0.72367 |
| *OsTLP23* | *OsTLP25* | *OsTLP18* | 313 | 20 | 27 | 1.04 | 0.30723 |
| *OsTLP26* | *OsTLP27* | *OsTLP18* | 411 | 34 | 34 | 0 | 1 |
| *SbTLP28* | *SbTLP29* | *SbTLP5* | 316 | 19 | 30 | 2.47 | 0.11608 |
| *SbTLP31* | *SbTLP34* | *SbTLP5* | 312 | 27 | 32 | 0.42 | 0.51508 |
| *SbTLP32* | *SbTLP33* | *SbTLP5* | 347 | 2 | 2 | 0 | 1 |
| *TaTLP1-A* | *TaTLP24-D* | *TaTLP4-D* | 483 | 37 | 30 | 0.73 | 0.39245 |
| *TaTLP1-B* | *TaTLP24-B* | *TaTLP4-D* | 486 | 36 | 31 | 0.37 | 0.5413 |
| *TaTLP12-B* | *TaTLP13-D* | *TaTLP4-D* | 321 | 24 | 33 | 1.42 | 0.23323 |
| *TaTLP15-A* | *TaTLP18-B* | *TaTLP4-D* | 391 | 36 | 38 | 0.05 | 0.81615 |
| *TaTLP15-B2* | *TaTLP18-A* | *TaTLP4-D* | 260 | 28 | 27 | 0.02 | 0.89274 |
| *TaTLP16-A* | *TaTLP17-A* | *TaTLP4-D* | 291 | 36 | 31 | 0.37 | 0.5413 |
| *ZmTLP1* | *ZmTLP35* | *ZmTLP28* | 551 | 36 | 33 | 0.13 | 0.71798 |
| *ZmTLP2* | *ZmTLP34* | *ZmTLP28* | 541 | 34 | 25 | 1.37 | 0.24132 |
| *ZmTLP3* | *ZmTLP22* | *ZmTLP28* | 560 | 11 | 14 | 0.36 | 0.54851 |
| *ZmTLP9* | *ZmTLP12* | *ZmTLP28* | 372 | 6 | 12 | 2 | 0.1573 |
| *ZmTLP13* | *ZmTLP37* | *ZmTLP28* | 464 | 19 | 22 | 0.22 | 0.63941 |
| *ZmTLP16* | *ZmTLP33* | *ZmTLP28* | 551 | 5 | 7 | 0.33 | 0.5637 |
| *ZmTLP20* | *ZmTLP25* | *ZmTLP28* | 525 | 17 | 23 | 0.9 | 0.34278 |

**Nt**- Identical sites in all three sequences, **Na**-Unique differences in Sequence A, **Nb**- Unique
differences in Sequence B

### 3.4. Gene architecture, domain and motif analyses

The number of exons varied from one to three in *O. sativa,* while one to four in *B. distachyon, S. bicolor* and *T. aestivum*. In the case of *Z. mays,* most of the *TLPs* consisted of one to three exons, while *ZmTLP16, ZmTLP20, ZmTLP25* and *ZmTLP33* exhibited eleven, seven, eight and ten exons, respectively. Moreover, a total of 44% (98/222) identified *TLPs* were intronless. Intriguingly, all the *sTLPs* were intronless, except *TaTLP6-A* and *TaTLP20-A1*. The intron phase analysis revealed that the occurrence of a maximum number of introns in phase 1, followed by phase 2, while the least number of introns were in phase 0 (**supplementary fig. S2**).

The functional nature of a protein depends upon the occurrence of domain composition. All of the identified cereals’ TLPs consisted of a thaumatin domain (PF00314), which confirmed that they are thaumatin-like proteins. The size of the thaumatin domain ranged from 202-217 AAs to134-154 AAs in the long and small TLPs, respectively. In addition to the thaumatin domain, ZmTLP16, ZmTLP20, ZmTLP25 and ZmTLP33 also consisted of a nuclear protein 96 (NUP96) domain of ∼211 AAs at the C-terminus of these proteins (**supplementary file S3**).

Motif investigation revealed the occurrence of 10 highly conserved motifs in the TLP proteins. Motifs 1-9 were parts of the thaumatin domain, while motif 10 was unknown. Motifs 5, 6 and 8 were the most conserved motifs in TLP proteins, in which the thaumatin signature sequence was present in motif 8 (**supplementary fig. S2**). The occurrence of conserved motifs across the cereal species suggested the conserved nature of TLP proteins in related plant species.

### 3.5. Physicochemical properties

To understand the various important features, several physicochemical properties of identified TLPs were studied. The TLPs were analyzed for molecular weight (MW), peptide length, isoelectric point (pI), subcellular localization, transmembrane (TM) helix and signal peptide (**supplementary file S4**). The average length of the long and small TLPs ranged from284-334 AAs to 175-185 AAs in the studied cereal crops, respectively. Similarly, the average MWs ranged from ∼29-35 kDa to ∼17-19 kDa, respectively. However, the average pI ranged from 5.89 to 6.95 and from 4.90 to 6.82 for the long and small TLPs, respectively (Table 3, **supplementary file S4**).

**Table 3.** Physicochemical properties of long and small TLP proteins.

| **Property** | **Type** | *Brachypodium*<br>*distachyon* | *Oryza*<br>*sativa* | *Sorghum*<br>*bicolor* | *Triticum*<br>*aestivum* | *Zea mays* |
| --- | --- | --- | --- | --- | --- | --- |
| **Average** | Long TLPs | 299 | 289 | 294 | 284 | 334 |
| **Protein length** | Small TLPs | 185 | 178 | 178 | 177 | 175 |
| **Average** | Long TLPs | 30.8 | 29.8 | 30.0 | 29.1 | 35.0 |
| **Molecular**<br>**weight (kDa)** | Small TLPs | 19.5 | 18.1 | 18.3 | 18.22 | 17.7 |
| **Average** | Long TLPs | 6.18 | 6.95 | 6.05 | 6.18 | 5.89 |
| **pI** | Small TLPs | 6.82 | 6.06 | 5.74 | 6.17 | 4.90 |

The majority of TLP proteins lacked TM helices, which suggested their soluble/cytoplasmic nature. However, five TLPs of *B. distachyon* and *O. sativa*, seven of *T. aestivum* and *Z. mays*, and 10 TLPs of *S. bicolor* consisted of a single TM region, and OsTLP18, TaTLP4-D and TaTLP5-D comprised of two TM helices, which suggested their membrane-bound nature (**supplementary file S4**).

A total of 23, 21, 36, 86 and 33 TLPs of *B. distachyon, O. sativa, S. bicolor*, *T. aestivum* and *Z. mays*, consisted of the N-terminal signal peptide, respectively. Further, the majority of TLP proteins were predicted to be localized in the extracellular region, while three TLPs from *O. sativa* (OsTLP10, OsTLP15 and OsTLP16) showed nuclear, and BdTLP16, TaTLP21-A, ZmTLP20, ZmTLP25 and ZmTLP33 showed plasma membrane localization (**supplementary file S4**). To validate the subcellular localization, a *TLP* gene (*TaTLP2-B*) was cloned and expressed with YFP as a translational fusion protein. Analysis of YFP fluorescence of fusion protein confirmed their extracellular localization (Fig. 4). Moreover, we could also see some fluorescence in the cytoplasmic region. The results suggested that the majority of TaTLP2-B was localized in the extracellular region, where they could be involved in various defence related functions (such as anti-fungal etc.). The cytoplasmic localization could be due to their translation in the cytoplasm, or it might also function inside the cytoplasm.

**Fig. 4.**
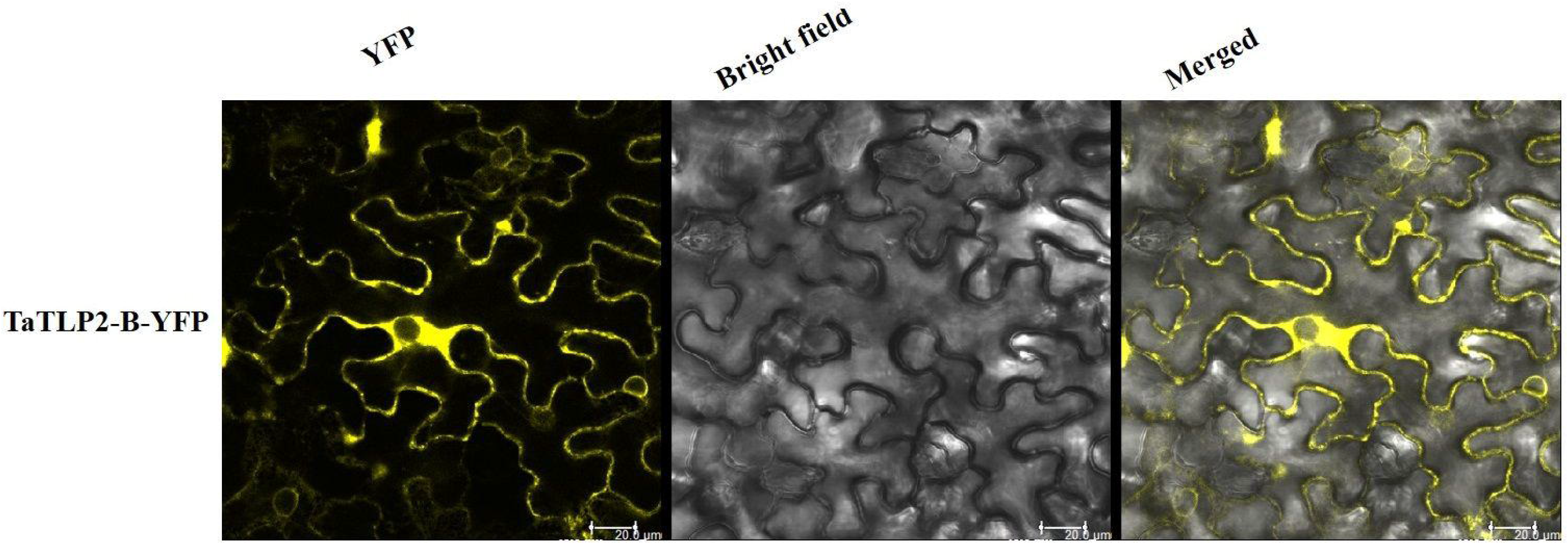
Subcellular localization analysis of TaTLP2-B by YFP fusion. The YFP fluorescence depicts the localization of TaTLP2-B protein majorly in the extracellular region.

### 3.6. Promoter region analyses

Promoter elements are necessary for the regulation of gene expression under different conditions. Therefore, we performed the *cis*-regulatory analysis of *TLPs* to foresee their regulatory mechanisms. Based on their putative functions, the identified *cis*-regulatory elements were segregated into four groups; growth and development, light, hormone and stress-responsive elements (**supplementary table S1**). The promoter regions of *TLP* genes comprised of several growth-related elements including TE2F2NTPCNA, EBOX, RYREPEAT4, RAV1AAT, POLLEN1LEAAT52 etc. In the case of light-responsive elements, GT1CONSENSUS, GATA box, GBOXLERBCS, BOXII, IBOXCORE were some common *cis*-regulatory elements. However, under the hormonal responsive category, DPBFCOREDCD3, ARFAT, ARR1, ERELEE4 and GARE10OSREP1 were abscisic acid, auxin, cytokinin, ethylene and gibberellic acid-responsive elements, respectively.

### 3.7. Expression profiling of the TLP genes under different tissues and developmental stages

Expression analysis of genes is a significant way to understand their involvement in various developmental and physiological processes. Previously, TLPs are reported to be involved in plant growth and developmental processes [56, 57]. Therefore, we performed the expression analysis of TLPs under various tissues and their developmental stages. A total of 23, 27, 37, 84 and 29 *TLP* genes showed expression in one or more tissue developmental stages in *B. distachyon, O. sativa, S. bicolor, T. aestivum* and *Z. mays*, respectively (Fig. 5A-E). Based on expression profile, these genes were clustered into 3-5 groups in various crop species. *BdTLPs* of group 1 were highly expressed in early and emerging inflorescence stages, while *BdTLP9* and *BdTLP12,* and *BdTLP20* and *BdTLP23* were upregulated in endosperm and embryo, respectively. Group 2 genes were highly expressed in the leaf, pistil, embryo and endosperm tissues. However, group 3 genes were highly expressed at 10 days after pollination (DAP) of seeds; moreover, *BdTLP8* and *BdTLP18* were upregulated in the pistil, as well (Fig. 5A). In *O. sativa*, the majority of group 1 genes were highly expressed in seed, group 2 in callus, group 3 genes in the shoot, group 4 in inflorescence and group 5 genes in emerging inflorescence and anther (Fig. 5B). In *S. bicolor,* group 1 and group 4 genes showed higher expression in root and leaf tissues, respectively. However, groups 2 and 3 *TLP* genes were highly expressed in various reproductive tissues like anther, pistil, endosperm, embryo and flower (Fig. 5C).

**Fig. 5.**
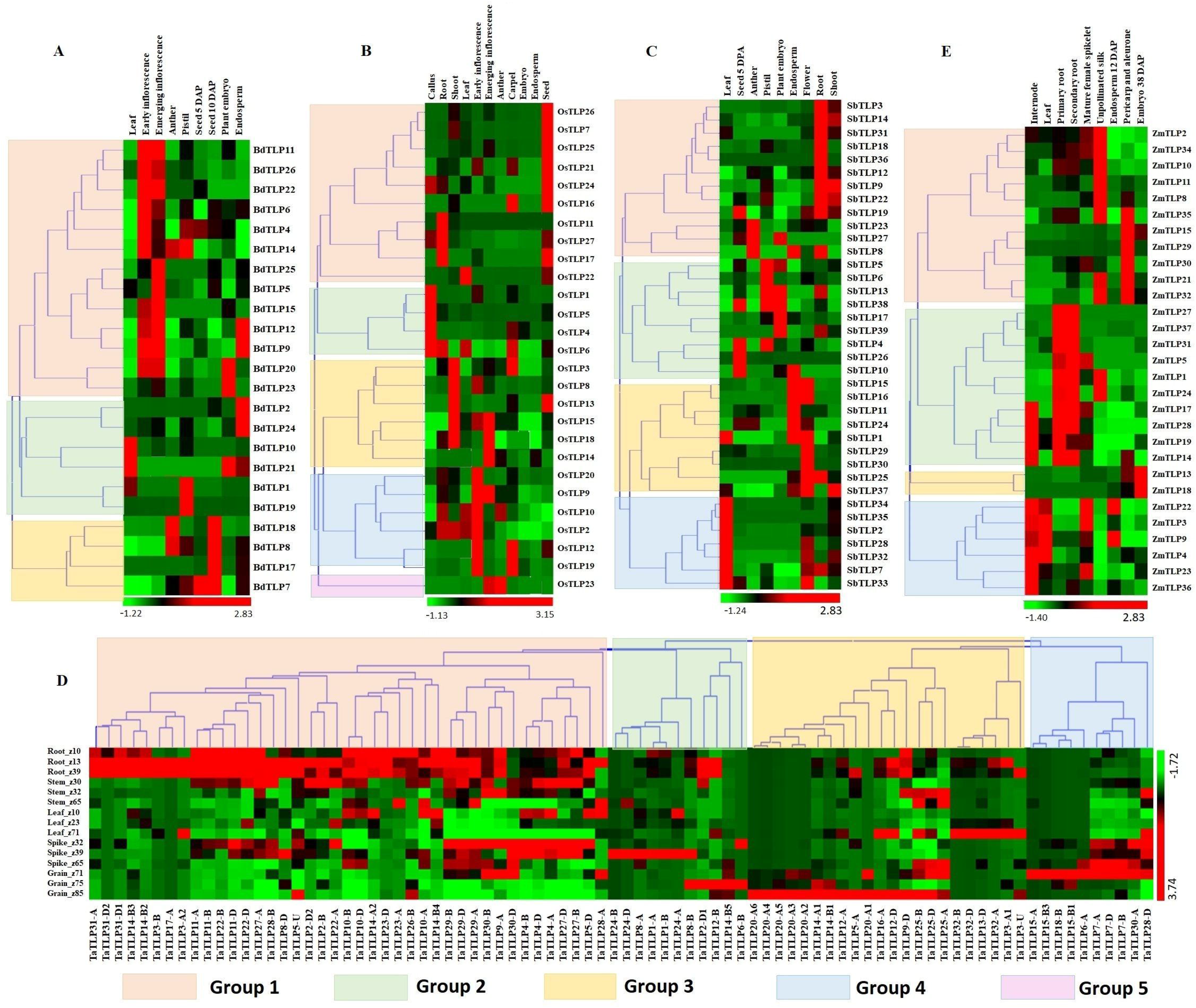
Expression analysis of *TLP* genes of *B. distachyon* (*BdTLP*), *O. sativa* (*OsTLP*). *S. bicolor* (*SbTLP*), *T. aestivum* (*TaTLP*) and *Z. mays* (*ZmTLP*) under various tissue and developmental stages. Heatmaps **(A-E)** shows the clustering and expression profiling of *TLP* genes in (**A**) *B. distachyon*, **(B)** *O. sativa*, **(C)** *S. bicolor*, **(D)** *T. aestivum* and **(E)** *Z. mays*. The different clusters are formed based on expression values of *TLP* genes in the respective crop which are represented with different colours. The colour bar depicts the upregulated and downregulated expression with red and green colours, respectively.

In *T. aestivum*, the majority of group 1 genes exhibited higher expression in root tissue followed by the shoot. However, most of the groups 2, 3 and 4 *TLP* genes were highly expressed in various developmental stages of spike and grain tissues (Fig. 5D).

In the case of *Z. mays,* group 1 genes showed significant expression in unpollinated silk, pericarp and aleurone layer. While, the majority of group 2 genes were specifically upregulated in the root tissues, and group 4 genes in the leaf and internode tissues. Group 3 genes were specifically upregulated in the embryo (Fig. 5E). These results suggested the role of *TLP* genes in both vegetative and reproductive tissues development in all the studied cereal crops. However, the higher expression of a few genes in a specific tissue or developmental stages suggested their precise role in particular tissue development.

### 3.8. Expression profiling of the TaTLP genes under biotic and abiotic stress conditions

Since TLPs belong to the defence-related protein family and are well known for their stress-responsive behaviour. The expression analysis of *TaTLP* genes was also performed under various biotic and abiotic stress conditions to reveal their roles in stress resistance [47–49]. For biotic stress, the available RNA-seq data generated at 24, 48 and 72 h of infestation of two important fungal pathogens, namely, *Blumeria graminis* f. sp. *tritici* (Bgt) and *Puccinia striiformis* f. sp. *tritici* (Pst) was used for expression analysis [47]. In *T. aestivum*, a total of 50 *TaTLP* genes exhibited differential expression (≥2 fold) in these stress conditions, which could be clustered into four groups (Fig. 6A). The majority of group 1 *TaTLP* genes were upregulated at 24 and 48 h of Bgt infestation. However, all the group 3 and 4 *TaTLP* genes were significantly upregulated at the late (72 h) and early (24 h) periods of Pst infestations, respectively. *TaTLP3-A2*, *TaTLP12-A,* and *TaTLP32-A* were the most upregulated genes after Bgt infection with 37, 36 and 35 folds, respectively. However, in the case of Pst infestation, *TaTLP10-A* (51 fold up) and *TaTLP27-D* (17 fold up) were highly upregulated.

**Fig. 6.**
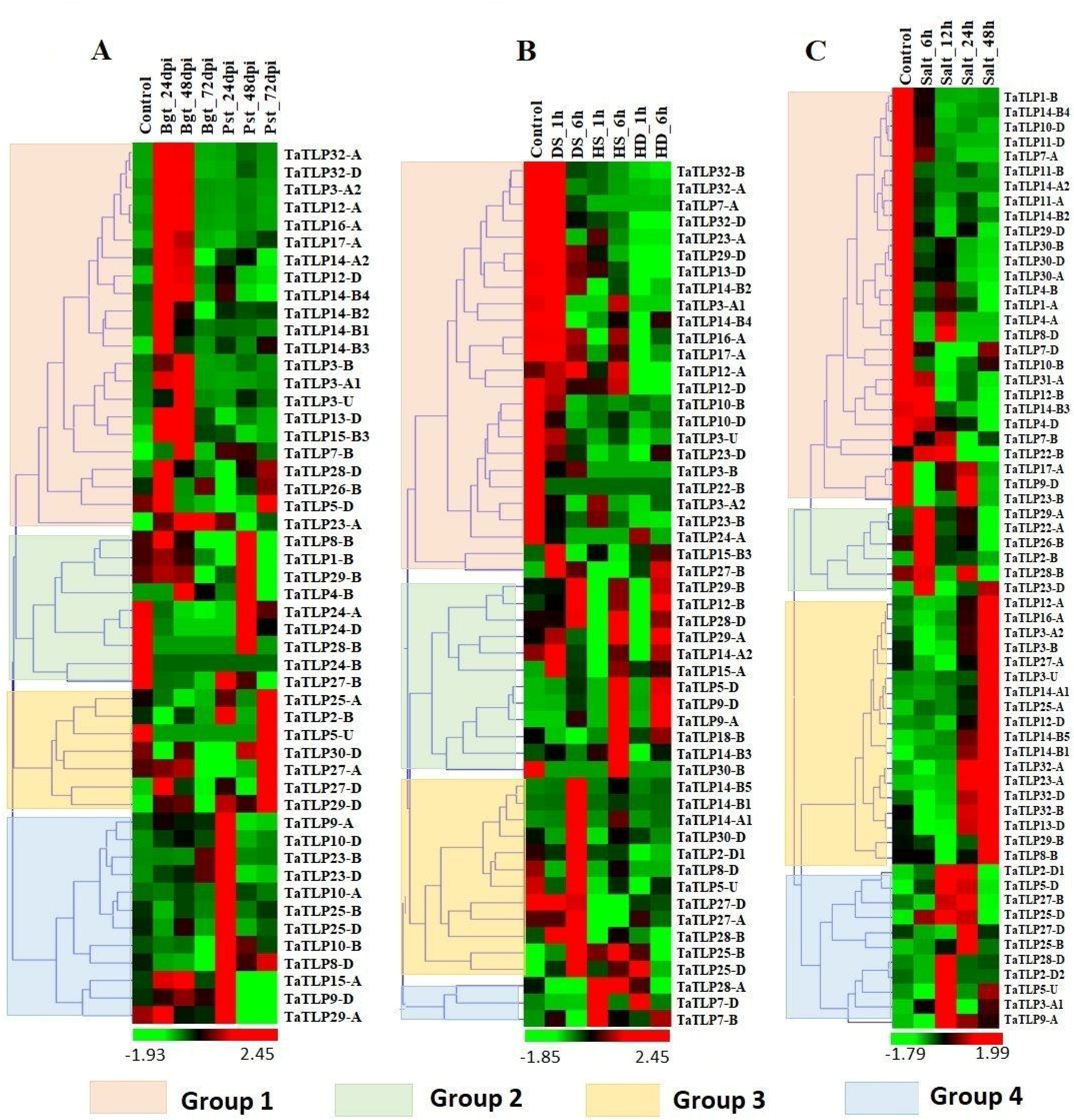
Expression profiling of *TLP* genes of *T. aestivum* under biotic and abiotic stress conditions. Heat maps **(A-C**) shows the clustering and expression profiling of *TaTLP* genes under **(A)** *Blumeria graminis* (Bgt) and *Puccinia striiformis* (Pst), **(B)** heat (HS), drought (DS) and combined heat-drought (HD) and **(C)** salt stress conditions. The clustering of *TaTLP* genes in each heat map is based on their expression pattern in respective stress conditions. The colour bar shows high and low expressions with red and green colours, respectively.

Abiotic stresses are another major threat to plants, which affect various physiological and biological processes. To understand the putative role of *TaTLPs* under abiotic stress conditions, expression analysis was carried out under drought (DS), heat (HS) and combined heat and drought (HD) stress using online available RNA-seq data [48]. A total of 52 *TaTLPs* exhibited significant differential expression in these stresses, which formed four clusters in the heatmap (Fig. 6B). Almost all the genes of group 1 were downregulated in the DS, HS and HD stresses, except *TaTLP12-A* that was slightly upregulated in DS treatment. In the case of group 2, the majority of genes were upregulated at 6 h of DS, HS and HD treatments, while either unaffected or downregulated at 1 h of treatment. These might be the late responsive *TaTLP* genes. Most of the group 3 and group 4 *TaTLP* genes were found to be DS and HS responsive, respectively.

Expression analysis of the *TaTLP* genes was also performed at 6, 12, 24 and 48 h of salt stress using available RNA-seq data [49]. Out of 93 genes, 63 *TaTLP* genes were found to be differentially expressed under salt stress, which were clustered into 4 groups, based on their expression patterns (Fig. 6C). All the group 1 genes were downregulated at all the stages of salt stress treatment. In contrast, all the group 3 genes were found significantly upregulated at 48 h of treatment. However, group 2 genes were upregulated at the early stage (6 h) of treatment, while they get normalized at later stages. The results suggested that the group 2 and group 3 genes are early and late responsive *TaTLP* genes, respectively.

To validate the RNA-seq expression data, qRT-PCR analysis of eight *TaTLP* genes was performed using gene-specific primers at various hours of heat, drought and salt treatments (Fig. 7A-P**, supplementary file S5**). Overall the result was in agreement with the expression observed using RNA-seq data, with a few exceptions. In the case of heat stress, *TaTLP2-B*, *TaTLP7-D*, *TaTLP14-B1* and *TaTLP25-B* were more upregulated at 1 h, while other genes were highly upregulated at 6 h of treatment. In the case of drought stress, *TaTLP-7D* was exclusively more upregulated at 1 h, whereas other genes were upregulated at 6 h. However, *TaTLP2-B* and *TaTLP10-D* were downregulated at DS 1 h. The majority of genes were highly upregulated at 1 h HD treatment. In the case of salt stress, seven genes were highly upregulated at 12 h of salt stress, except *TaTLP14-B1* that showed maximum expression at 48 h. The results further confirmed that a few *TaTLP* genes are early while others are late responsive.

**Fig. 7.**
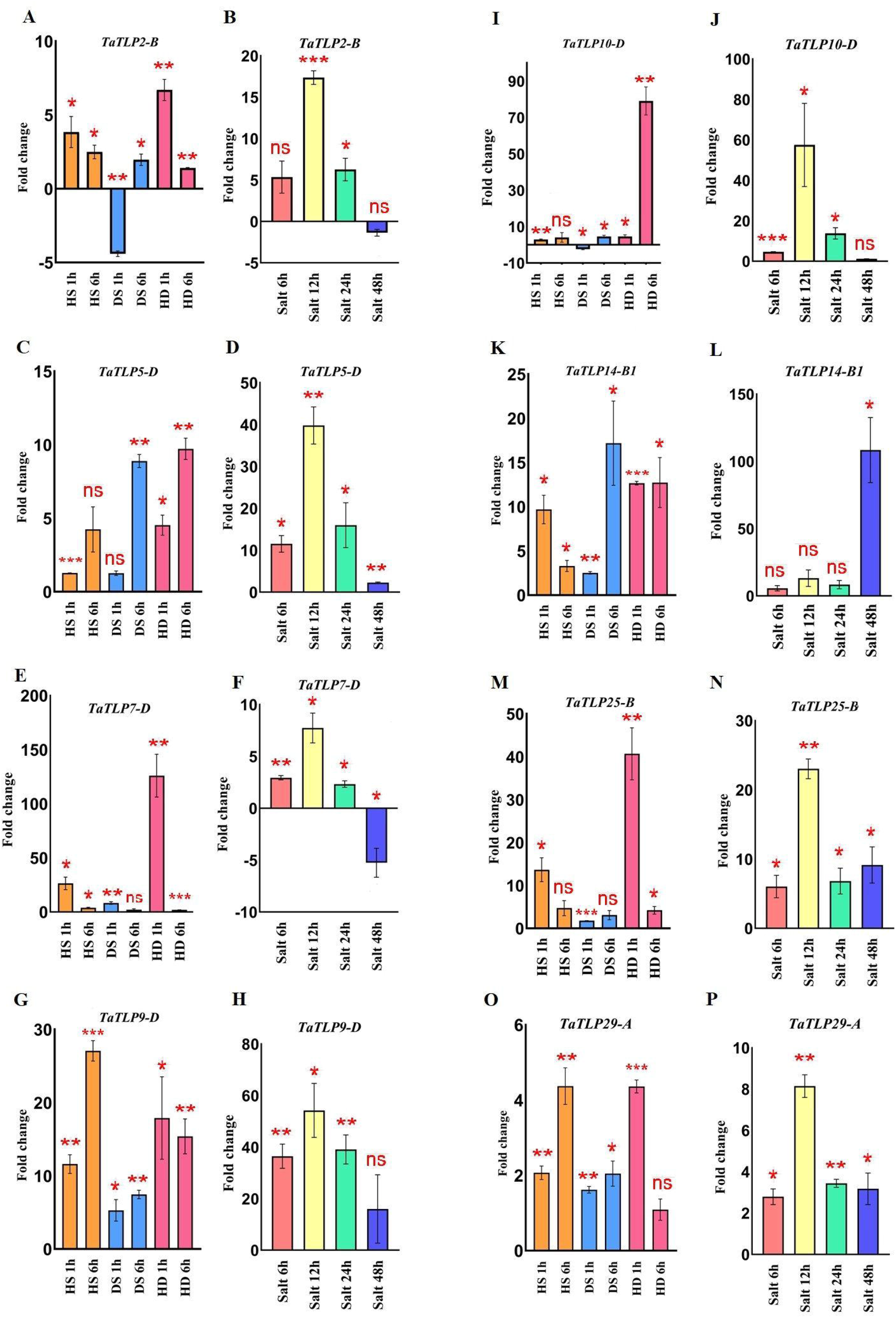
qRT-PCR analysis of eight *TaTLP* genes. Expression patterns in the form of fold changes have been shown in the bar graphs. Figures 7 A, C, E, G, I, K, M**, and** O show expression of *TaTLP2-B*, *TaTLP5-D, TaTLP7-D, TaTLP9-D, TaTLP10-D, TaTLP14-B1, TaTLP25-B and TaTLP29-A* genes, under HS, DS and HD stress conditions, while figure 7 B, D, F, H, J, L, N**, and** P shows the expression pattern of these genes under salt stress, respectively. Significance between the control and treated conditions is carried using a two-tailed student’s t-test. The ns, *, ** and *** markings represent the significance at p-value >0.05, <=0.05, <=0.01, <=0.001, respectively.

### 3.10. Comparative expression profiling of the TaTLP paralogous genes

To understand the function of duplicated genes, a comparative expression profiling of each paralogous pair was performed. Generally, based on expression pattern, duplicated genes could categorize into the retention of function, pseudo-functionalization and neo-functionalization. Out of six paralogous gene pairs, three pairs (*TaTLP1-A-TaTLP24-D, TaTLP1-B-TaTLP24-B, TaTLP15-A-TaTLP18-B*) showed a similar trend of expression, suggested retention of function in these duplicated genes (Fig. 8A-C). Two duplicated pairs (*TaTLP12-D-TaTLP13-D* and *TaTLP16-A-TaTLP17-A*) showed insignificant expression of one of the gene suggested pseudo-functionalization (Fig. 8D, E). Retention of function in the majority of duplicated genes and the absence of neo-functionalization suggested functional conservation in *TLP* genes during evolution.

**Fig. 8.**
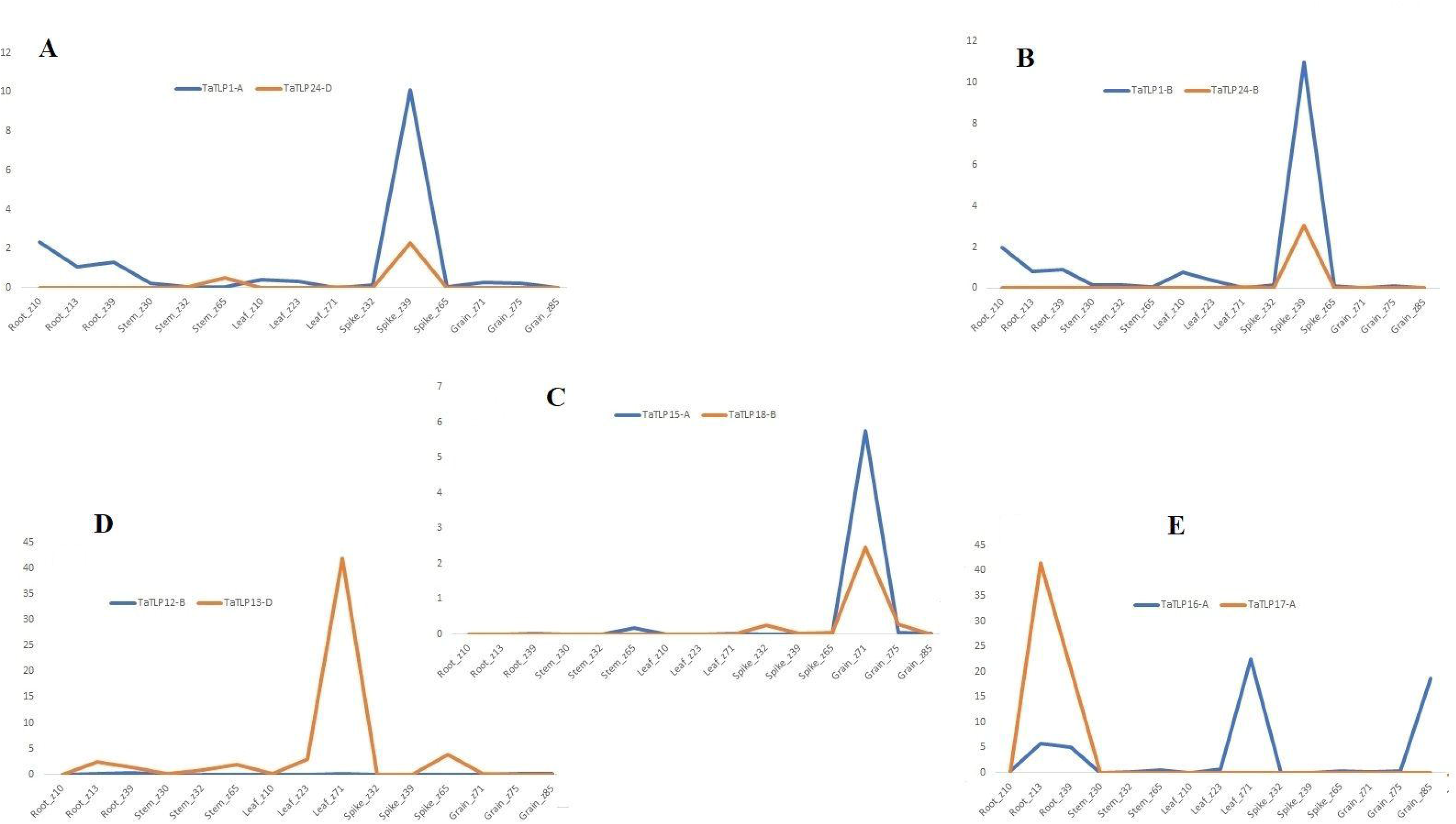
Comparative expression analysis of duplicated gene pairs in *T. aestivum*. Graphs **(A-E)** represents the comparative expression pattern of paralogous *TaTLPs* under various tissue and developmental stages. Based on their expression trend, *TaTLPs* are categorized into the retention of function **(A-C)** and pseudo functionalization types **(D, E)**.

### 3.11. Cloning and functional characterization of the TaTLP2-B in Saccharomyces cerevisiae

In our study, since the *TaTLP2-B* exhibited significant differential expression under abiotic stress conditions, it was selected for functional characterization. *TaTLP2-B* open reading frame was amplified using end primers and cloned into pYES2.1 yeast expression vector. The *LacZ* gene-containing vector was used as control (Fig. 9A-F). In the spot assay, we observed similar growth of both control (LacZ) and TaTLP2-B expressing yeast cells in the controlled conditions (Fig. 9A). However, in the case of cold, drought, heat, combined heat drought and salt stress conditions, TaTLP2-B recombinant cells exhibited higher growth than the control cells (Fig. 9B-F). The results revealed that the overexpression of the TaTLP2-B provided abiotic stress tolerance to the recombinant yeast cells. This gene can be used for the development of abiotic stress-resistant transgenic crops in future studies.

**Fig. 9.**
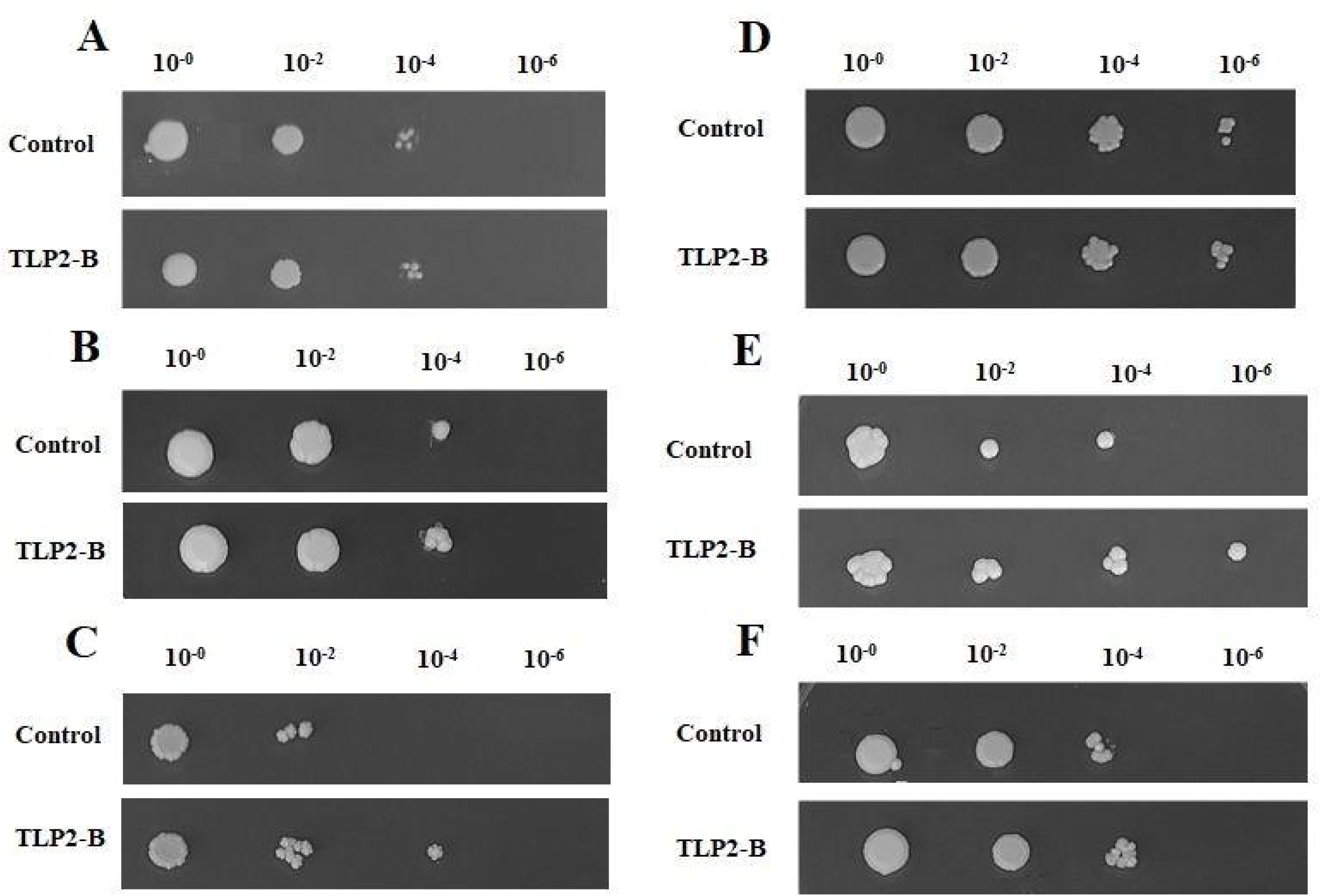
Spot assay of recombinant *Saccharomyces cerevisiae* (yeast) cells under various stress conditions. Figure **(A)** shows similar growth of both TaTLP2-B (pYES2.1-TaTLP2-B) and control vector (pYES2.1/V5-His/lacZ) containing yeast cells under control conditions. Figure **(B-F)** shows the spot assay analysis of *TaTLP2-B* and control vector containing yeast cells in (**B**) cold (4°C), (**C**) drought (30% PEG), (**D**) heat (37°C), (**E**) combined heat and drought (37°C and 30% PEG), and (**F**) salt (1M NaCl) stress conditions, respectively.

## 4. Discussion

TLPs are important proteins involved in the tissue development and stress resistance pathways of plants [6]. Therefore, we have studied the TLPs in five major cereals including *B*. *distachyon*, *O*. *sativa*, *S*. *bicolor*, *T*. *aestivum* and *Z*. *mays*. In the present study, we have identified a total of 26, 27, 39, 93 and 37 *TLPs* in the genome of *B*. *distachyon*, *O*. *sativa*, *S*. *bicolor*, *T*. *aestivum* and *Z*. *mays*, respectively. Variable numbers of the *TLP* genes in different plants have been reported in earlier studies. For instance, 24, 33 and 49 *TLP* genes have been reported in diploid *A*. *thaliana*, *Vitis vinifera* and *Populus trichocarpa*, respectively [7, 58]. In the case of *O*. *sativa* and *Z*. *mays*, 49 *TLP* genes in each are reported in the earlier study [7], because probably they had not removed the genes with incomplete thaumatin signature motifs, as done in the present study. The occurrence of the highest number of *TLP* genes in *T*. *aestivum* could be attributed to their complex allohexaploid genome configuration (AABBDD) [20]. Further, differential expansion of the *TLP* genes in various plant species might be attributed to the number of duplication events.

The duplication events are responsible for the expansion of gene families and these are the major forces to attain neo-functionality by causing genetic variability [59]. In DEs analysis, our results suggested that tandem DEs are a major factor in *B*. *distachyon*, *O*. *sativa* and *S*. *bicolor* and segmental DEs in *T*. *aestivum* and *Z*. *mays* behind the gene expansion of TLP family. The role of tandem and segmental duplications in the evolution of TLP gene family has been earlier reported in various plant species including *A*. *thaliana*, *O*. *sativa*, and *Z*. *mays* [7]. The higher number of paralogous genes in *T*. *aestivum* and *Z*. *mays* might be attributed to their larger genome size and the presence of more transposable elements [53, 60].

Furthermore, the divergence time of monocots from eudicots was estimated as ∼170-235 Mya, which were further diverged into grasses around ∼77 Mya [60, 61]. Therefore, the paralogous *TLP* genes in *B*. *distachyon* might have evolved before the divergence of grasses. The main duplication incidence in *S*. *bicolor* and *O*. *sativa* genome was estimated at around ∼70 Mya [62]. However, our results indicated that the majority of paralogous *TLP* genes of *O*. *sativa* and *S*. *bicolor* are evolved after the main DE occurrence. In *Z*. *mays*, paralogous *TLP* genes were probably formed close to the divergence time of maize and sorghum, i.e. ∼11-28 Mya [62]. However, in the case of *T*. *aestivum*, paralogous *TLP* genes were probably evolved earlier than the hybridization event of A, B and D subgenomes [63]. However, two duplicated pairs, having NUP 96 domain showed their recent incidence of DEs. Duplicated genes face various environmental forces and natural pressures during the process of evolution [53]. Our results suggested the purifying selection of *TLP* genes as a major force of selection, as reported in the various other plant species [7, 64]. Moreover, positive selection has also been reported for *TLP* genes in poplar, which suggested varied natural selection procedures during TLPs’ evolution [8]. Since the TLPs are paraphyletic in origin, they are supposed to be derived from multiple ancestral genes [65]. Clustering of identified TLPs in multiple clades is in agreement with the paraphyletic nature of origin. Further, a variable number of TLPs in each clade is due to the differential duplication of *TLP* genes. Similarly, the uneven distribution and paraphyletic nature of TLPs have also been reported by previous studies [7, 64]. The tight clustering of homeologous and paralogous genes could be due to the high sequence homology among the respective sequences.

The organization of introns and exons of *TLPs* have been reported to be in the range of one to ten exons [6,7,9,53]. On similar patterns, our results suggest comparable findings with one to four exons in most of the TLPs. However, most of the monocots’ TLPs have intron-less nature, which was also found in our analysis [7]. Additionally, our results with respect to the protein lengths, molecular weights and isoelectric points of long and small TLP proteins, found in accordance with previously studied TLPs [6,7,9]. In our analysis, the TLP2-B was found to be localized in extracellular region. Similarly, the extracellular localization of a wheat TLP (TaLr19TLP1) is also reported in an earlier study [66].

In addition, the presence of thaumatin-like domain in all the TLP proteins, make our studies similar to previously reported findings. However, four ZmTLPs (ZmTLP16, ZmTLP20, ZmTLP25, and ZmTLP33) have an additional NUP 96 domain. The NUP 96 domain is reported to be involved in plant development and stress resistance against pathogens [67]. Various abiotic and biotic stress-responsive elements were also found in most of the studied *TLPs*, for instance, DRE2COREZMRAB17, ASF1, BIHD1, CBFHV, CGG-box, GCCCORE, LTRECOREATCOR15, MYB, MYC, WRKY1, W-box, T/G-box etc. Various plant hormone-related and stress-related *cis*-regulatory elements have also been reported in four cotton species [56]. Moreover, in previous studies, the elements like ERE, W-box, WRKY, were found to be involved in plant growth and stress resistance [68–70]. The occurrence of numerous *cis*-regulatory elements suggested diverse functions of *TLP* genes in plants. Further, the expression profiling of *TLPs* under various tissues and developmental stages also advocated the putative involvement of these genes in development of vegetative and reproductive tissues. Similar expression trend in various plant tissues and their role in reproductive tissues like flowers and seeds has also been reported in earlier studies, for instance, *TLP* genes in *Gossypium hirsutum*, also showed their varied expression pattern in vegetative as well as in reproductive organs [18, 56, 71]. The role of *TLP* genes in fungal resistance has been reported in earlier studies against numerous pathogens including *Fusarium* sp. [72], *Microdochium nivale* [73], *Rhizoctonia solani* [74] and *Verticillium dahlia* [56] in wheat and other plant species. The differential expression of *TaTLP* genes in response to Pst and Bgt infestation further established their roles in fungal stress responses. Moreover, similar to our finding’s differential expression and upregulation of *TLP* genes at various hours of drought, salt and cold stress treatment has also been reported in other plants like *Brassica rapa* and *Gossypium* sp. [56, 75]. Further, the over-expression of *TLP* genes provided increased tolerance against various abiotic stresses in tobacco, cotton, Arabidopsis etc. [11,56,76]. In a recently conducted study, the important role of a cotton *TLP* (*GhTLP19*) gene was studied in combating drought stress [56]. Moreover, the GbTLP1 and ObTLP1 of *G*. *barbadense* and *Ocimum basilicum*, respectively, were found effective against drought and salt stresses [57, 76]. Collectively, current findings depict that the TLP proteins could be utilized to develop the stress resistance transgenic cereal crops, which will be a boost for the agriculture.

## 5. Conclusions

Thaumatin-like proteins (TLPs), a part of the PR5 protein family, are known to be involved in various biotic and abiotic stresses. The curiosity to unravel the various possibilities led us to a detailed characterization of a total of 222 *TLP* genes in five cereal crops. Phylogenetic analysis revealed the paraphyletic origin of TLPs in cereals, while the occurrence of duplication events suggested the role of paralogous genes in the expansion of the TLP gene family. Gene expression analysis using RNA-seq-data and qRT-PCR suggested the important roles of *TLP* genes in plant growth and development and abiotic and biotic stress responses. Significant tolerance in the *TaTLP2-B* expressing yeast cells further established their role in stress response. Overall the study revealed that the *TLP* genes can be used for stress-resistant transgenic crop development. Since the identified genes are directly derived from the food crops, they would be easily de-regulated from the regulatory authority of the transgenic crops. Further, the current study will facilitate the detailed functional characterization of *TLP* genes of cereal crops in future studies.

## Author’s contribution

SKU conceived the idea. AS and SKU designed the experiments. AS, HS and RR performed the experiments. AS, HS and AP analyzed the data. AS, HS and SKU wrote the manuscript. All the authors have read and finalized the manuscript.

## Acknowledgements

The authors are grateful to the Panjab University, Chandigarh, and National Institute of Plant Genome Research, New Delhi, India for research facilities, Ensembl Plants, URGI and NCBI for data availability. AS and HS is grateful to CSIR for the senior research fellowship. HS is also thankful to the IKGPTU, Jalandhar for the Ph.D registration. SKU is grateful to the Council of Scientific and Industrial Research (CSIR) and the Department of Science and Technology (DST), Government of India for a research grant (No. 38(1489)/19/EMR-II) and partial financial support under the Promotion of University Research and Scientific Excellence (PURSE) grant scheme, respectively.

## Supplementary files

**Figure S1.** Multiple sequence alignment of 222 TLPs of *B. distachyon, O. sativa, S. bicolor*, *T. aestivum* and *Z. mays*. Conserved cysteine residues are marked with an asterisk. The red dot denotes the REDDD motif. The FF hydrophobic motif is highlighted with a green triangle. The thaumatin signature motif, conserved domain, amino acids forming the bottom of the acidic cleft are marked with sky blue, red and brown lines, respectively.

**Figure S2.** Intron/exon organization and conserved motifs of TLPs of *B. distachyon*, *O. sativa*, *S. bicolor, T. aestivum* and *Z. mays*.

**File S1.** List of identified *TLP* genes with their proposed nomenclature in *B. distachyon, O. sativa S, bicolor, T. aestivum* and *Z. mays*. The Red colour represents the *sTLPs* of each crop.

**File S2.** List of paralogous *TLP* genes in *B. distachyon*, *O. sativa*, *S. bicolor*, *T. aestivum* and *Z. mays*.

**File S3**. Domain organization of TLP proteins of *B. distachyon*, *O. sativa*, *S. bicolor*, *T. aestivum* and *Z. mays*.

**File S4**. The characterization table of *B. distachyon, O. sativa*, *S. bicolor*, *T. aestivum* and *Z. mays* TLPs.

**File S5.** List of qRT-PCR primers.

**Table S1.** List of predicted *cis*-regulatory elements in the promoter region of *TLP* genes of *B. distachyon, O. sativa, S. bicolor, T. aestivum* and *Z, mays*.

