## Supplementary material for "Molecular characterization revealed the role of thaumatin-like proteins in stress response in bread wheat": Supplementary table 1.docx

Supplementary table S1. List of different predicted *cis*-regulatory elements in the promoter region of *TLP* genes of *B. distachyon, O. sativa, S. bicolor, T. aestivum* and *Z. mays*.

| *B. distachyon* | | | | |
| --- | --- | --- | --- | --- |
| Gene | Light Response | Growth and Development | Stress Response | Hormone Response |
| *BdTLP1* | GATA box, GT1CONSENSUS, I-box, PRE | CACT, DOFCORE, E-box, POLLEN1, RHEs | A-box, ACGT ELRE, ASF1, BP5, DRE, CRT/DRE, CuRE, MYB, MYC, T/G box, W-box, WRKY | ABRE, ARR1, CAREs, CGCG box, D1, DPBF |
| *BdTLP2* | BOXII, CIACADIAN, GATA box, GT1CONSENSUS, I-box, PRE, LRE, SORLIP, T-box | ANAERO, DOFCORE, E-box, E2F, POLLEN1 | ACGT, DRE, CuRE, MYB, MYC, LTRE, PREAT, SURE, TAAAG, W-box, WRKY, | ABRE, ARR1 |
| *BdTLP3* | BOXL, GT1, I-box | ANAERO, CACT, DOFCORE, E-box, LEAFY, L1-box, XYL | BIHD1, CuRE, DRE, ELRE MYB, MYC, RY- elements, SEBF, W-box, WRKY | ABRE, ARF, ACGT, ARR1, CPB, CAREs, GARE, |
| *BdTLP4* | GT1CONSENSUS,  GATA-box, BOXL, PRE, SORLIP | DOFCORE, E-box, POLLEN1 | BIHD1, CuRE, CRT/DRE, LTRE. MYB, MYC, RY-ELEMENTS, SURE, WRKY | ABRE, ARF, CAREs, GARE |
| *BdTLP5* | GT1CONSENSUS,  GATA-box, I-box, PRE, SORLIP, T-box | DOFCORE, E-box | ASF1, BIHD1, DRE, CuRE, LTRE, MYB, MYC, RY-ELEMENTS, T/G box, W-box, WRKY | ARF, ARR1, GARE |
| *BdTLP6* | GT1CONSENSUS,  GATA-box, I-box, SORLIP, T-box | ANAERO, CAN | ACGT, ASF1, CRT/DRE, LTRE, MYB, MYC, SURE, T/G box, W-box, WRKY | ABRE, ARF |
| *BdTLP7* | GT1CONSENSUS,  GATA-box, I-box, PRE, SORLIP | ANAERO, DOFCORE, POLLEN1, NODCON | ACGT, BIDIH1, CuRE, IRO, LTRE, MYB, MYC, W-box, WRKY | ABRE, ARR1, CGCG box, GARE |
| *BdTLP8* | GT1CONSENSUS,  GATA-box, I-box, SORLIP | ANAERO, DOFCORE, E-box | ACGT, BIDH1, DRE, CuRE, LTRE. MYB, MYC, W-box, WRKY, P1BS | ABRE, ARR1, DPBF, GARE |
| *BdTLP9* | BOXCP, GT1CONSENSUS,  GATA-box | ANAERO, CAN, E-box, POLLEN1, | ASF1, CuRE, MYC, W-box, WRKY | ABRE, NTBBF |
| *BdTLP10* | GT1CONSENSUS,  GATA-box, I-box, PRE, SORLIP | ANAERO, DOFCORE, E-box | ACGT, ASF1, CRT/DRE, CuRE, LTRE, MYB, MYC, W-box. WRKY | ARF, ARR1, DPBF |
| *BdTLP11* | GT1CONSENSUS,  GATA-box, PRE, RBCS, SORLIP | ANAERO, CAN, DOFCORE, E-box | ACGT, ASF1, BIHD1, CRT/DRE, CuRE, LTRE, MYB, MYC, W-box, WRKY, SURE | ABRE, ARF, ARR1, DPBF |
| *BdTLP12* | GT1CONSENSUS,  GATA-box, SORLIP | CAN, DOFCORE, E-box, POLLEN1 | ACGT, ASF1, CRT/DRE, CuRE, GCC-box, MYB, MYC, W-box, WRKY | ABRE, ARF, ARR1, DPFB, ERE |
| *BdTLP13* | BOXII, BOXL, GT1CONSENSUS,  GATA-box, I-box, SORLIP | CAN, DOFCORE, E- box, POLLEN1 | ASF1, CRT/DRE, CuRE, MYB, MYC, WRKY | ARF, ARR1, GARE, NTBBF |
| *BdTLP14* | GT1CONSENSUS,  GATA-box, PRE, SORLIP | AMMORE, ANAERO, CAN, DOFCORE, POLLEN1 | ACGT, ASF1, BIHD1, BOX-A, CRT/DRE, DRE, GCC-box, MYB, MYC, LTRE, W-box, WRKY | ABRE, ARF, ARR1, DPBF, ERE, NTBBF |
| *BdTLP15* | BOXII, GT1CONSENSUS,  GATA-box, I-box SORLIP | DOFCORE, E-box, POLLEN1, L1box | CRT/DRE, CuRE, DRE, LTRE, MYB, MYC, P1BS, T/G box, W-box, WRKY | ABRE, ARR1, AUX, GARE |
| *BdTLP16* | BOXII, GT1CONSENSUS,  GATA-box, I-box, PRE, SORLIP, Z-box | CAN, DOFCORE, E-box, POLLEN1 | ACGT, CRT/DRE, MYB, MYC, T/G box, W-box, WRKY | ABRE, DPBF, GARE, CPBC |
| *BdTLP17* | BOXII, BOXL, GT1CONSENSUS,  GATA-box, I-box, SORLIP | ANAERO, DOFCORE, E-box, POLLEN1 | ACGT, BIHD1, MYB, MYC, W-box, WRKY | GARE, NDE |
| *BdTLP18* | GT1CONSENSUS,  GATA-box, I-box, SORLIP | DOFCORE, E-box, POLLEN1 | ACGT, ASF1, LTRE, IRO, MYB, MYC, RY-ELEMENTS, W-box, WRKY | ABRE, ARR1, CARE, NDE |
| *BdTLP19* | GT1CONSENSUS,  GATA-box, SORLIP | ANAERO, CAN, DOFCORE, E-box, POLLEN1 | ASF1, BIHD1, BOX-A, CuRE, MYB. MYC, RY-ELEMENTS, W-box, WRKY | ARF, ARR1 |
| *BdTLP20* | BOX-C, CIACADIAN, GT1CONSENSUS,  GATA-box, I-box, PRE, SORLIP | ANAERO, CAN, DOFCORE, POLEEN1 | BIHD1, CuRE, CTR/DRE, DRE, LTRE, W-box, WRKY | ARF, DPBF |
| *BdTLP21* | CIACADIAN, GT1CONSENSUS,  GATA-box, SORLIP | ANAERO, CAN, DOFCORE, E-box, POLLEN1 | ACGT, BIHD1, LTRE, MYB, MYC, TAAAG, W-box, WRKY | ABRE, DPBF |
| *BdTLP22* | GT1CONSENSUS,  GATA-box, I-box, T-box | DOFCORE, E-box, POLLEN1, NRR | B1HD1, CuRE, DRE, BOX-A, YB, MYC, SEBF, W-box, WRKY | ARF, ARR1, GARE |
| *BdTLP23* | GT1CONSENSUS,  GATA-box, I-box, PRE, SORLIP | ANAERO, CAN, DOFCORE, E-box, POLLEN1 | ACGT, ASF1, BIHD1, BOX-A, CTR/DRE, CuRE, DRE, GCC-box, LTRE, IRO, MYB, MYC, RY-ELEMENTS, W-box, WRKY | ABRE, ARF, ARR1, DPBF, GARE |
| *BdTLP24* | BOXC, BOXL, GT1CONSENSUS,  GATA-box, I-box, PRE | CARG, E-box | ACGT, BIHD1, BOX-A, CTR/DRE, MYB, MYC, T/G- box, RY-ELEMENTS. QAR, W-box, WRKY | ABRE, ARF, ARR1, GARE |
| *BdTLP25* | GT1CONSENSUS,  GATA-box, LRE-box, PRE, SORLIP | E-box, POLLEN1 | ACGT, BIHD1, CTR/DRE, CuRE, LTRE, MYB, MYC, W-box, WRKY | ABRE, GARE, NTBBF |
| *BdTLP26* | BOXC, BOXL, BOXII, GT1CONSENSUS,  GATA-box, I-box, PRE, SORLIP | CAN, DOFCORE, POLLEN1, Q-ELEMENT | ACGT, BIHD1, BOX-A, CTR/DRE, CuRE, DRE, LTRE, MYB, MYC, RY-ELEMENTS, W-box, WRKY | ABRE, ARF, ARR1, ERE, NTBBF, PROXB |
| *O. sativa* | | | | |
| *OsTLP1* | BOXL, I-box, GT1CONSENSUS,  GATA-box, SORLIP | ANAERO, DOFCORE, E-box, GTGA, GCN4, POLLEN1, PROLAMIN-box, Q-ELEMENT | ACGT, CuRE, MYB, MYC, W-box, WRKY | ABRE, ARF, ARR1, DPBF, S-box |
| *OsTLP2* | GT1CONSENSUS,  GATA-box, SORLIP, T-box | DOFCORE, E-box, GTGA, POLLEN1, RY-ELEMENTS, SURE | AGMOTIF, BIHD1, CTR/DRE, CuRE, GCC, MYB, MYC, SEBF, TAAAG, W-box, WRKY | ABRE, ERE, TATC-box, TCA-element |
| *OsTLP3* | BOXC, GT1CONSENSUS,  GATA-box, I-box, PRE, SORLIP, T-box, RBCS | ANAERO, CEREGLUE- box, DOFCORE, E- box, NAPIN, POLLEN1, RY-ELEMENT | ACGT, ASF1, BIHD1, CTR/DRE, CuRE, DRE, LTRE, MYB, MYC, SURE, TAAAG, W-box | ARR1, DPBF, NTBBF |
| *OsTLP4* | BOXL, BOXII, GT1CONSENSUS,  GATA-box, I-box, PRE, SORLIP | ANAERO, CIACADIAN, DOFCORE, E-box, POLLEN1, S1F, SRE | ACGT, ASF1, BIHD1, CTR/DRE, CuRE, DRE, GCC- box, LTRE, MYB, MYC, P1BS | ABRE, CARE, CGCG- box, DPBF, GARE |
| *OsTLP5* | GT1CONSENSUS,  GATA-box, I-box, SORLIP, T-box | DOFCORE, E-box, NAPIN, POLLEN1, SRE | ACGT, CuRE, MYB, MYC, P1BS | ABRE, CARE, NDE, DPBF, GARE, |
| *OsTLP6* | GT1CONSENSUS,  GATA-box, I-box, PRE, RBCS | CAN, DOFCORE, E-box, POLLEN1, SRE | ACGT, ASF1, BIHD1, CTR/DRE, MYB, MYC, T/G- box, QAR, W- box, WRKY | ABRE, NTBBF |
| *OsTLP7* | BOXL, BOXII, GT1CONSENSUS,  GATA-box, I-box, PRE, SORLIP | CAN, CEREGLUE- box, DOFCORE, E- box, NAPIN, POLLEN1, RY-ELEMENT, S1F, SRE | ACGT, ASF1, BIHD1, CTR/DRE, CuRE, DRE, LTRE, MYB, MYC, T/G- box, W-box | ABRE, CGCG- box, DPBF |
| *OsTLP8* | BOXL, GT1CONSENSUS,  GATA-box, I-box, RBCS | CIACADIAN, DOFCORE, E-box, POLLEN1, TGTCACA | BIHD1, CTR/DRE, MYB, MYC, W-box, WRKY | ARR1, CARE, NDE |
| *OsTLP9* | GT1CONSENSUS,  GATA-box, I-box, PRE, SORLIP | ANAERO, CAN, CELLCYLCE-box, E-box, POLLEN1, RY-ELEMENTS | ACGT, BIHD1, CuRE, MYB, MYC, W-box, WRKY | ABRE, CARE, CYTOSITE, DPBF, NTBBF, PROXB |
| *OsTLP10* | GT1CONSENSUS,  GATA-box, I-box, PRE, SORLIP | CAN, CIACADIAN, DOFCORE, E-box, POLLEN1 | ACGT, GCC- box, MYB, MYC, TAAAG, UPR | ABRE, ARR1, CGCG-box, NTBBF |
| *OsTLP11* | BOXL, GT1CONSENSUS,  GATA-box, I-box, SORLIP, T-box | DOFCORE, E-box, POLLEN1, SRE, TGATCA | ACGT, BIHD1, CuRE, GCC- box, CTR/DRE, DRE, LTRE, MYB, MYC, T/G- box, W-box, WRKY | ABRE, CGCG- box, DPBF, NTBBF |
| *OsTLP12* | GT1CONSENSUS,  GATA-box, I-box | ANAERO, DOFCORE, E-box, POLLEN1 | BIHD1, CuRE, MYB, MYC, P1BS, SEBF, W-box, WRKY | ARF, ARR1 |
| *OsTLP13* | BOXL, BOXII, GT1CONSENSUS,  GATA-box, I-box, SORLIP | ANAERO, DOFCORE, E-box, L1BOX, NAPIN, POLLEN1, SRE | ACGT, BIHD1, CuRE, LTRE, MYB, MYC, P1BS, W-box, WRKY | ABRE, ARR1, NDE |
| *OsTLP14* | GT1CONSENSUS,  GATA-box, I-box, PRE, SORLIP | DOFCORE, E-box, NAPIN, POLLEN1 | ACGT, BIHD1, MYB, MYC, W-box, WRKY | ARR1, CARE, NTBBF |
| *OsTLP15* | GT1CONSENSUS,  GATA-box, I-box, SORLIP | ANAERO, DOFCORE, E-box, POLLEN1, RY-ELEMENTS, S1F, SRE | CuRE, MYB, MYC, P1BS, W-box, WRKY | ARF, ARR1, NDE, NTBBF |
| *OsTLP16* | GT1CONSENSUS,  GATA-box, I-box, SORLIP | CACT, DOFCORE, E-box, POLLEN1, RY-ELEMENTS, SRE, SPBF | BIDH1, CuRE, MYB, MYC, W-box, WRKY | ARR1, CPB |
| *OsTLP17* | BOXII, GT1CONSENSUS,  GATA-box, I-box, PRE | ANAERO, CELLCYLCE-box, CIACADIAN, DOFCORE, E- box, POLLEN1, Q-ELEMENT | ACGT, BIHD1, CACG, IRO, MYB, MYC, SURE, TAAAG, AW-box, WRKY | ABRE, ARR1, CGCG- box, DPBF, ERE, NTBBF |
| *OsTLP18* | BOXL, GT1CONSENSUS,  GATA-box, I-box, PI, PRE, SORLIP, T- box | CAN, DOFCORE, POLLEN1, SRE | ACGT, ASF1, CuRE, MYB, MYC, P1BS, W-box, WRKY | ARR1, NDE, DPBF, ERE |
| *OsTLP19* | BOXL, GT1CONSENSUS,  GATA-box, I-box, H-box, PRE, SORLIP | ANAERO, DOFCORE, E-box, POLLEN1, RY-ELEMENTS, SRE | BIHD1, CuRE, GCC-box, MYB, MYC, W-box, WRKY | ABRE, ARR1, NDE, GARE, GRA |
| *OsTLP20* | BOXL, BOXII, GT1CONSENSUS,  GATA-box, I-box, PRE, SORLIP, T-box | ANAERO, CAN, CEREGLUE- box, DOFCORE, POLLEN1, RY-ELEMETS, SRE | BIHD1, CTR/DRE, CuRE, LTRE, MYB, MYC, W-box, WRKY | ARR1, DPBF |
| *OsTLP21* | GT1CONSENSUS,  GATA-box, I-box, SORLIP | CAN, DOFCORE, E-box | ACGT, BIHD1, BP5, CTR/DRE, ELRE, MYB, MYC. T/G- box, W-box, WRKY | ABRE, ARR1, CARE, DPBF, GARE |
| *OsTLP22* | BOXL, GT1CONSENSUS,  GATA-box, I-box, RBCS | CELLCYLCE-box, CIACADIAN, DOFCORE, E- box, POLLEN1, Q-ELEMENT, RY-ELEMENTS | CGT, CuRE, GCC-box, MYB, MYC, W-box, WRKY | ABRE, ARR1, NDE |
| *OsTLP23* | GATA-box, PRE, SORLIP | CAN, CIACADIAN, DOFCORE, E-box, POLLEN1, RY-ELEMENTS | ACGT, BIHD1, BP5, LTRE, MYB, MYC, P1BS, QAR, T/G- box | ABRE, CGCG- box, DPBF |
| *OsTLP24* | BOXII, GT1CONSENSUS,  GATA-box, I-box | ANAERO, DOFCORE, E-box, NAPIN, POLLEN1, RY-ELEMENTS | ACGT, CuRE, MYB, MYC, P1BS, W-box, WRKY | ARR1, CARE, DPBF, NTBBF |
| *OsTLP25* | GT1CONSENSUS,  GATA-box, I-box, PRE, SORLIP | CAN, CIACADIAN, DOFCORE, E-box, L1BOX, POLLEN1, RY-ELEMENTS | ACGT, CuRE, MYB, MYC, P1BS, W-box, WRKY | ABRE, ARR1, GARE |
| *OsTLP26* | BOXII, GT1CONSENSUS,  GATA-box, I-box, PRE, SORLIP | 56/59- box, CIACADIAN, ANAERO, DOFCORE, E-box, POLLEN1, RY-ELEMENTS | ACGT, BIHD1, CuRE, CTR/DRE, MYB, MYC, P1BS, W-box, WRKY | ABRE, ARR1, CGCG- box, DPBF, ERE, |
| *OsTLP27* | BOXII, GT1CONSENSUS,  GATA-box, I-box, PRE, SORLIP, T-box | DOFCORE, E-box, POLLEN1 | ACGT, BIHD1, CuRE, CTR/DRE, GCC-box, MYB, MYC, P1BS, W-box, WRKY | ABRE, ARR1, CARE, D4-ELEMENT, DPBF, GARE, NDE |
| *S. bicolor* | | | | |
| *SbTLP1* | GT1CONSENSUS,  GATA-box, PRE, SORLIP, T-box | ANAERO, CACT, DOFCORE, E-box, NAPINRY-ELEMENTS,  SITE II-ELEMENT | ACGT, BIHD1, CTR/DRE, DRE, CuRE, LTRE, MYB, MYC, SURE, TAAAG, W-box, WRKY | ARF, ARR1, CARE, CPB |
| *SbTLP2* | BOXL, GT1CONSENSUS,  GATA-box, I-box, PRE, SORLIP | CACT, CIACADIAN, DOFCORE, E-box, POLLEN1 | ACGT, CTR/DRE, CuRE, DRE, ELRE, LTRE, MYB, MYC, SURE, TAAAG, W-box, WRKY | ABRE, ARF, ARR1, CPB, DPBF, ERE |
| *SbTLP3* | BOXL, GT1CONSENSUS,  GATA-box, I-box, PRE, SORLIP | ANAERO, CACT, CIACADIAN, DOFCORE, E-box, POLLEN1, RY-ELEMENTS, UP2 | ACGT, ELRE, ASF1, LTRE, MYB, T/G-box, TAAAG, UPR, W-box, WRKY | ABRE, ARF, ARR1, CPB, AUX, CGCG-box, DPBF, ERE, GARE |
| *SbTLP4* | GT1CONSENSUS,  GATA-box, I-box, SORLIP | ANAERO, CACT, CEREGLUE-box DOFCORE, E-box, POLLEN1, RY-ELEMENTS | ACGT, BIHD1, CTR/DRE, CuRE, DRE, ELRE, LTRE, MYB, MYC, SURE, W-box, WRKY | ABRE, ARR1, CPB, CGCG-box |
| *SbTLP5* | BOXL, BOXII, GT1CONSENSUS,  GATA-box, I-box, PRE, RBCCS, SORLIP, T-box | ANAERO, CACT, DOFCORE, E-box, POLLEN1 | ACGT, ASF1, CuRE, ELRE, LTRE, MYB, MYC, SEBF, W-box, WRKY | ARR1, ARF, CGCG-box, GARE, NDE, NTBBF |
| *SbTLP6* | GT1CONSENSUS,  GATA-box, PRE, SORLIP, T-box | CACT, CEREGLUE-box DOFCORE, EVENING-ELEMENT, E-box, POLLEN1, RY-ELEMENTS | ACGT, ASF1, BIHD1, CTR/DRE, CuRE, DRE, LTRE, MYB, MYC, SURE, TAAAG-box, W-box, WRKY | ABRE, ARR1, CARE, CGCG-box, D1, DPBF, NTBBF |
| *SbTLP7* | BOXL, BOXII, GT1CONSENSUS,  GATA-box, I-box, PRE, SORLIP | ANAERO, CACT, DOFCORE, E-box, POLLEN1 | A-box, GCC box, HSE, LTR, MBS, TC-rich repeats | ABRE, ARR1, CGCG-box, NTBBF |
| *SbTLP8* | BOXL, GT1CONSENSUS,  GATA-box, I-box, PRE, SORLIP | CAT-box, CCGTCC-box, O2-site, circadian | ACGT, BIHD1, CTR/DRE, CuRE, DRE, LTRE, MYB, MYC, SURE, TAAAG-box, W-box, WRKY | ABRE, ARF, ARR1, CARE, CGCG-box, DPBF, GARE, NTBBF |
| *SbTLP9* | BOXL, BOXII, GT1CONSENSUS,  GATA-box, SORLIP | CACT, CEREGLUE-box, CIACADIAN, DOFCORE, E-box, POLLEN1, RY-ELEMENTS | ACGT, CTR/DRE, CuRE, DRE, LTRE, MYB, MYC, SURE, TAAAG-box, W-box, WRKY | ABRE, ARR1, CPB, DPBF, GARE, NTBBF |
| *SbTLP10* | GT1CONSENSUS,  GATA-box, I-box, PRE | CACT, CREGLUE-box, DOFCORE, RY-ELEMENT, SRE | CuRE, MYB, MYC, P1BS, TAAAG-box, W-box, WRKY | ARR1, CPB |
| *SbTLP11* | BOXL, GT1CONSENSUS,  GATA-box, I-box, PRE, SORLIP | ANAERO, CACT, CAN, CEREGLUE-box, DOFCORE, RY-ELEMENTS, Q-ELEMENTS, | ACGT, BIHD1, CTR/DRE, CuRE, MYB, MYC, SURE, T/G-box, TAAAG-box, QAR, W-box, WRKY | ABRE, ARR1, CARE, CGCG-box, DPBF, PROX B, NTBBF |
| *SbTLP12* | GT1CONSENSUS,  GATA-box, I-box, PRE, T-box | CACT, DOCORE, E-box, LEAFY, POLLEN1 | ASF1, CuRE, MYB, MYC, P1BS, TAAAG-box, W-box, WRKY | ARR1, CGCG-box, GARE, NDE |
| *SbTLP13* | GT1CONSENSUS,  GATA-box, I-box, PRE, SORLIP | ANAERO, CACT, CIACADIAN, DOFCORE, E-box, POLLEN1, RY-ELEMENTS, SRE | ACGT, ARE1, CTR/DRE, CuRE, DRE, ELRE, GCC-box, LTRE, MYB, MYC, P1BS, SEBF, SURE, TAAAG-box, W-box, WRKY | ABRE, AGC-box, ARR1, ARF, CGCG-box, DPBF, GARE, NTBBF |
| *SbTLP14* | BOX C, GT1CONSENSUS,  GATA-box, I-box, PRE, SORLIP, T- box | ANAERO, CACT, DOFOCRE, E-box, L1 BOX, Q-ELEMENT, RY-ELEMENT, SRE | ACGT, BIHD1, CTR/DRE, CuRE, DRE, GCC-box, LTRE, MYB, MYC W-box, WRKY | ABRE, ARR1, CGCG-box, DPBF |
| *SbTLP15* | GT1CONSENSUS,  GATA-box, I-box, RBCS, SORLIP | ANAERO, CACT, DOFCORE, E-box, POLLEN1, RY--ELEMENTS | ACGT, BIHD1, CTR/DRE, CuRE, DRE, GCC-box, LTRE, MYB, MYC SURE, W-box, WRKY, T/G-box, TAAAG-box | ABRE, ARR1, CGCG-box, DPBF, NTBBF |
| *SbTLP16* | BOX L, GT1CONSENSUS,  GATA-box, PRE, SORLIP | ANAERO, CACT, CAN, DOFOCRE, E-box, Q-ELEMENT, RY-ELEMENT | ACGT, BIHD1, CTR/DRE, CuRE, DRE, ELRE, GCC-box, LTRE, MYB, MYC, T/G-box, W-box, WRKY | ABRE, ARR1, CGCG-box, DPBF |
| *SbTLP17* | BOX L, GT1CONSENSUS,  GATA-box, I-box, SORLIP, T-box | ANAERO, CACT, CAN, CIACADIAN, DOFCORE, E-box, L1 BOX, SRE | ACGT, BIHD1, CTR/DRE, CuRE, DRE, MYB, MYC, W-box, WRKY | ABRE, ARR1, ARF, CARE, ERE, |
| *SbTLP18* | GT1CONSENSUS,  GATA-box, I-box, SORLIP | ANAERO, CACT, CAN, CIACADIAN, DOFCORE, E-box, POLLEN1, RY-ELEMENTS, SRE | ACGT, BIHD1, CuRE, MYB, MYC, SURE, TAAAG-box, W-box, WRKY | ARR1, CPB, NTBBF |
| *SbTLP19* | BOX C, BOX II, GT1CONSENSUS,  GATA-box, I-box, PRE, SORLIP | ANAERO, CACT, CAN, DOFCORE, E-box, RY-ELEMENTS, SRE | ACGT, BIHD1, CTR/DRE, CuRE, DRE, LTRE, MYB, MYC, SURE, TAAAG-box, W-box, WRKY | ARR1, NTBBF |
| *SbTLP20* | BOX II, GT1CONSENSUS,  GATA-box, I-box | ANAERO, CACT, CAN, DOFCORE, E-box, HDZIP, POLLEN1, | BIHD1, ELRE, MYB, MYC, TAAAG-box, WRKY | ARR1, DPBF, GARE |
| *SbTLP21* | GT1CONSENSUS,  GATA-box, I-box, PRE, SORLIP | CACT, POLLEN1, SRE | ACGT, BIHD1, BOX A, ELRE, MYB, MYC, SURE, TAAAG-box, W-box, WRKY | ABRE, ARR1, DPBF, GARE, NTBBF |
| *SbTLP22* | BOX L, GT1CONSENSUS,  GATA-box, I-box, PRE, SORLIP | ANAERO, CACT, DOFCORE, E-box, POLLEN1, UP1 | ACGT, BIHD1, CTR/DRE, CuRE, DRE, LTRE, MYB, MYC, TAAAG-box, W-box, WRKY | ABRE, ARR1, CGCG-box, DPBF, ERE, GARE |
| *SbTLP23* | GT1CONSENSUS,  GATA-box, I-box, PRE, SORLIP | ANAERO, CACT, DOFCORE, E-box, POLLEN1, RY-ELEMENTS | ACGT, ARE1, BIHD1, CTR/DRE, DRE, GCC-box, LTRE, IRO, MYB, MYC, SURE, T/G-box, UPR, W-box, WRKY | ABRE, AGC-box, ARR1, CGCG-box, DPBF, GARE, NDE, S-box |
| *SbTLP24* | GT1CONSENSUS,  GATA-box, I-box, RBCS, SORLIP | ANAERO, CACT, DOFCORE, E-box, GCN4, GLM, POLLEN1, Q ELEMENT, SRE | ACGT, CuRE, MYB, MYC, SURE, TAAAG-box, W-box, WRKY | ABRE, ARR1, CPB, DPBF, GARE, NDE |
| *SbTLP25* | GT1CONSENSUS,  GATA-box, I-box, PRE, RBCS | CACT, CIACADIAN, DOFCORE, E-box, POLLEN1, RY-ELEMENTS, SRE | ACGT, BIHD1, CTR/DRE, CuRE MYB, MYC, SURE, TAAAG-box, W-box, WRKY | ARF, ARR1, CGCG-box, DPBF, NTBBF |
| *SbTLP26* | BOX II, GT1CONSENSUS,  GATA-box, SORLIP | CACT, CAN, CEREGLUE-box, CIACADIAN, DOFCORE, E-box, POLLEN1, | ACGTA, BIHD1, CuRE, MYB, MYC, TAAAG, WRKY | ABRE, ARR1, CGCG-box, CPB, DPBF |
| *SbTLP27* | BOX II, SORLIP, T-box | ANAERO, CACT, CAN, CIACADIAN, DOFCORE, E-box, POLLEN1, Q-ELEMENT, RY-ELEMENTS | ACGT, BIHD1, CuRE, ELRE, LTRE, MYB, MYC, W-box, WRKY | ABRE, ARR1, AUX, DPFB, ERE, |
| *SbTLP28* | GT1CONSENSUS,  GATA-box, I-box, PRE, RBCS, SORLIP | ANAERO, CACT, CIACADIAN, DOFCORE, E-box, POLLEN1 | ACGT, BIHD1, CuRE, MYB, MYC, QAR, W-box, WRKY | ABRE, ARR1, DPBF, |
| *SbTLP29* | GT1CONSENSUS,  GATA-box, PRE, SORLIP | ANAERO, CACT, CAN, CIACADIAN, DOFCORE, E-box, POLLEN1, RY-ELEMENTS | ACGT, BIHD1, CuRE, ELRE, SEBF, SURE, T/G- box, TAAAG-box, MYB, MYC, W-box, WRKY | ABRE, ARR1, ARF, DPBF, NDE, NTBBF |
| *SbTLP30* | BOX II, GT1CONSENSUS,  GATA-box, SORLIP, T-box | CACT, CIACADIAN, DOFCORE, POLLEN1, XYL | BIHD1, MYB, SEBF, SURE, TAAAG-box, W-box, WRKY | ARF, ARR1, CARE, DPBF, NTBBF |
| *SbTLP31* | BOX II, GT1CONSENSUS,  GATA-box, I-box, SORLIP, T-box | ANAERO, CACT, CELLCYLCE-box, CIACADIAN, DOFCORE, E-box, L1 BOX, NAPIN, POLLEN1, RY-ELEMENTS, SRE | ACGT, ASF1, BIHD1, CuRE, ELRE, GCC-box, LTRE, MYB, MYC W-box, WRKY | ABRE, ARR1, CARE, DPBF, ERE |
| *SbTLP32* | GT1CONSENSUS,  GATA-box, I-box, RBCS, SORLIP | ANAERO, CACT, CIACADIAN, DOFCORE, E-box, L1 BOX, POLLEN1, Q-ELEMENT, RY-ELEMENTS | ACGT, BIHD1, CTR/DRE, CuRE, MYB, MYC, SEBF, SURE, W-box, WRKY | ABRE, ARR1, ARF, CARE, GARE |
| *SbTLP33* | GT1CONSENSUS,  GATA-box, I-box, SORLIP | CACT, CIACADIAN, DOFCORE, E-box, L1 BOX, POLLEN1, Q-ELEMENT, RY-ELEMENTS, SRE | ACGT, BIHD1, CTR/DRE, CuRE, MYB, MYC, SURE, W-box, WRKY | ARR1, CARE, NTBBF |
| *SbTLP34* | BOX II, GT1CONSENSUS,  GATA-box, I-box | ANAERO, CACT, CAN, DOFCORE, E-box, POLLEN1, RY-ELEMENTS, SRE | BIHD1, CuRE, DRE, ELRE, MYB, MYC, W-box, WRKY | ABRE, ARR1, CGCG-box, CPB, DPBF, GARE, |
| *SbTLP35* | BOX L, GT1CONSENSUS,  GATA-box, SORLIP | ANAERO, CACT, CIACADIAN, DOFCORE, E-box, POLLEN1 | BIHD1, CuRE, GCC-box, MYB, MYC, SEBF, W-box, WRKY | ARF, ARR1, CARE, CPB, DPBF, |
| *SbTLP36* | BOX II, GT1CONSENSUS,  GATA-box, SORLIP, T-box | ANAERO, CACT, DOFCORE, E-box, POLLEN1 | ACGT, BIHD1, CTR/DRE, CuRE, DRE, GCC-box, LTRE, MYB, MYC, P1BS, SURE, W-box, WRKY | ABRE, ARR1, CGCG-box, CPB, DPBF |
| *SbTLP37* | BOX II, GT1CONSENSUS,  GATA-box, SORLIP, T-box | ANAERO, CACT, CAN, DOFCORE, SRE | ACGT, BIHD1, BP5, CuRE, GCC-box, MYB, MYC, QAR, T/G-box, W-box, WRKY | ABRE, ARR1, GARE, NTBBF |
| *SbTLP38* | BOX II, BOX L, GT1CONSENSUS,  GATA-box, RBCS, SORLIP | ANAERO, CACT, DOFCORE, E-box, POLLEN1, RY-ELEMENTS | BIHD1, CTR/DRE, DRE, ELRE, LTRE, MYB, MYC, SURE, TAAAG-box, W-box, WRKY | ARR1, ARF, CPB, DPBF |
| *SbTLP39* | BOX II, BOX C, BOX L, GATA-box, PRE, RBCS, SORLIP | ANAERO, CACT, DOFCORE, E-box, NAPIN, POLLEN1, Q-ELEMENT, RY-ELEMENTS, SPH | ACTG, BIHD1, GCC-box, LTRE, MYB, MYC, T/G-box, UPR, W-box, WRKY | ABRE, ARR1, CARE, CGCG-box, DPBF |
| *T. aestivum* | | | | |
| *TaTLP1-A* | GT1CONSENSUS,  GATA-box, I-box, PRE, SORLIP, SV40 | ANAERO, CACT, DOFCORE, E-box, POLLEN1, RY-ELEMENTS, SPH | ACGT, ASF1, BIHD1, CTR/DRE, CuRE, DRE, ELRE, LTRE, MYB, MYC, QAR, TAAAG-box, T/G-box, UPR, W-box, WRKY | ABRE, ARR1, CGCG-box, CPB, DPBF, NTBBF |
| *TaTLP1-B* | BOX L, GT1CONSENSUS,  GATA-box, I-box, PRE, SORLIP, SV40 | ANAERO, CACT, DOFCORE, E-box, POLLEN1, Q-ELEMENT, RY-ELEMENTS, SPH | ACGT, ASF1, BIHD1, BOX-A, CTR/DRE, CuRE, DRE, GCC-box, LTRE, MYB, MYC, QAR, TAAAG-box, W-box, WRKY | ABRE, ARR1, CARE, DPBF, NDE |
| *TaTLP2-A* | BOX II, GT1CONSENSUS,  GATA-box, I-box, PRE, RBCS, SORLIP | ANAERO, CACT, CAN, DOFCORE, E-box, POLLEN1 | CuRE, GCC-box, LTRE, MYB, MYC, P1BS, SURE, TAAAG-box, W-box, WRKY | ARF, ARR1 |
| *TaTLP2-B* | BOX II, GT1CONSENSUS,  GATA-box, I-box, PRE, RBCS, SORLIP | ANAERO, CACT, CAN, DOFCORE, E-box, POLLEN1, RY-ELEMEMNTS | CuRE, GCC-box, LTRE, MYB, MYC, P1BS, SURE, TAAAG-box, W-box, WRKY | ARF, ARR1 |
| *TaTLP2-D1* | BOX II, GT1CONSENSUS,  GATA-box, PRE, RBCS, SORLIP, T-box | ANAERO, CACT, CAN, DOFCORE, E-box, POLLEN1, RY-ELEMEMNTS | CTR/DRE, CuRE, DRE, LTRE, MYB, MYC, SURE, TAAAG-box, W-box, WRKY | ARR1, CARE, DPBF |
| *TaTLP2-D2* | GT1CONSENSUS,  GATA-box, PRE, RBCS, SORLIP, T-box | ANAERO, CACT, CAN, DOFCORE, E-box, POLLEN1, RY-ELEMEMNTS | MYB, MYC, P1BS, SURE, TAAAG-box, W-box, WRKY | ARF, ARR1, CARE |
| *TaTLP3-A1* | BOX L, GT1CONSENSUS,  GATA-box, I-box, PRE, RBCS, SORLIP, SV40 | ANAERO, CACT, CIACADIAN, DOFCORE, E-box, POLLEN1, SRE | ACGT, IRO-, MYB, MYC, BOX-A, SEBF, SURE, TAAAG-box, W-box, WRKY | ABRE, ARF, ARR1, CARE, CGCG-box, DPBF, ERE, NTBBF |
| *TaTLP3-A2* | GT1CONSENSUS,  GATA-box, I-box, SORLIP, SV40 | CACT, CAN, DOFCORE, E-box, POLLEN1, RY-ELEMEMNTS, SRE | ACGT, ASF1, CuRE, IRO, MYB, MYC, SURE, TAAAG-box, W-box, WRKY | ABRE, ARR1, CARE, DPBF, ERE, NTBBF |
| *TaTLP3-B* | GT1CONSENSUS,  GATA-box, I-box, PRE, SORLIP, SV40 | CACT, DOFCORE, E-box, POLLEN1, Q-ELEMENT | ACGT, ASF1, BIHD1, BOX-A, CTR/DRE, CuRE, DRE, GCC-box, IRO, LTRE, MYB, MYC, SURE, W-box, WRKY | ABRE, ARR1, DPBF |
| *TaTLP3-U* | BOX L, GT1CONSENSUS,  GATA-box, I-box, PRE, SORLIP, SV40 | ANAERO, CACT, DOFCORE, E-box, POLLEN1, RY-ELEMEMNTS, SRE | ACGT, ASF1, BIHD1, CuRE, IRO, MYB, MYC, SEBF, SURE, TAAAG-box, W-box, WRKY | ARF, ARR1, CARE, DPBF, NTBBF |
| *TaTLP4-A* | BOX II, GT1CONSENSUS,  GATA-box, SORLIP, SV40 | ANAERO, CACT, DOFCORE, E-box, POLLEN1 | ACGT, ASF1, BIHD1, CuRE, DRE, GCC-box, MYB, MYC, TAAAG-box WRKY | ABRE, ARR1, CARE, CGCG-box, DPBF, GARE ERE, NTBBF |
| *TaTLP4-B* | BOX II, GT1CONSENSUS,  GATA-box, SORLIP | ANAERO, CACT, DOFCORE, E-box, POLLEN1, RY-ELEMEMNTS | ACGT, BIHD1, BOX-A, CTR/DRE, CuRE, DRE, LTRE, MYB, MYC, TAAAG-box, W-box, WRKY | ABRE, ARR1, DPBF, NTBBF |
| *TaTLP4-D* | BOX L, GT1CONSENSUS,  GATA-box, I-box, PRE, SORLIP, T-box | ANAERO, CACT, CAN, DOFCORE, E-box, POLLEN1, RY-ELEMEMNTS | ACGT, BIHD1, BOX-A, CTR/DRE, CuRE, DRE, LTRE, MYB, MYC, P1BS, SEBF, SURE, TAAAG-box, T/G-box, W-box, WRKY | ABRE, ARR1, CGCG-box, NDE, NTBBF |
| *TaTLP5-A* | GT1CONSENSUS,  GATA-box, I-box, SORLIP | ANAERO, CACT, CIACADIAN, DOFCORE, E-box, POLLEN1, Q-ELEMEMNT | ACGT, BIHD1, CTR/DRE, CuRE, DRE, ELRE, GCC-box, LTRE, MYB, MYC, SEBF, SURE, TAAAG-box, W-box, WRKY | ARF, ARR1, CGCG-box |
| *TaTLP5-D* | GT1CONSENSUS,  GATA-box, I-box, SORLIP | ANAERO, CACT, CIACADIAN, DOFCORE, E-box, POLLEN1, Q-ELEMEMNT | ACGT, BIHD1, CTR/DRE, CuRE, DRE, ELRE, LTRE, MYB, MYC, SEBF, SURE, TAAAG-box, W-box, WRKY | ARF, ARR1 |
| *TaTLP5-U* | GT1CONSENSUS,  GATA-box, PRE, SORLIP, T-box | ANAERO, CACT, CIACADIAN, DOFCORE, E-box, POLLEN1, Q-ELEMEMNT | ACGT, CTR/DRE, CuRE, DRE, GCC-box, LTRE, MYB, MYC, SEBF, SURE, TAAAG-box, W-box, WRKY | ARF, ARR1, CGCG-box, DPFB |
| *TaTLP6-A* | GT1CONSENSUS,  PRE, SORLIP, T-box | CACT, DOFCORE, E-box, POLLEN1 | ACGT, BIHD1, CuRE, MYB, MYC, SURE, TAAAG-box, W-box, WRKY | CGCG-box, DPBF, GARE, NTBBF |
| *TaTLP6-B* | GT1CONSENSUS,  PRE, SORLIP, T-box | ANAERO, CACT, CIACADIAN, DOFCORE, E-box, POLLEN1 | BIHD1, BOX-A, CTR/DRE, CuRE, LTRE, MYB, MYC, P1BS, SURE, TAAAG-box, W-box, WRKY | ARR1, DPBF, |
| *TaTLP6-D* | GT1CONSENSUS,  GATA-box, SV40, T-box | ANAERO, CACT, DOFCORE, POLLEN1 | ACGT, ASF1, BIHD1, CuRE, MYB, SURE, TAAAG-box, W-box, WRKY | ABRE, AAR1, CGCG-box, DPBF, GARE |
| *TaTLP7-A* | GT1CONSENSUS,  GATA-box, I-box, PRE, RBCS, SORLIP, SV40, T-box | ANAERO, CACT, DOFCORE, POLLEN1, RY-ELEMENTS | BIHD1, CTR/DRE, CuRE, MYB, MYC, TAAG-box, W-box, WRKY | ARR1, CPB, DPBF, ERE, NTBBF |
| *TaTLP7-B* | BOX II, GT1CONSENSUS,  GATA-box, I-box, RBCS, T-box | ANAERO, CACT, CAN, DOFCORE, E-box, POLLEN1, RY-ELEMENTS | BIHD1, BOX-A, CTR/DRE, CuRE, MYB, MYC, TAAG-box, W-box, WRKY | ARR1, NTBBF |
| *TaTLP7-D* | BOX II, GT1CONSENSUS,  GATA-box, I-box, RBCS, T-box | ANAERO, CACT, CAN, DOFCORE, E-box, POLLEN1, RY-ELEMENTS | BIHD1, BOX-A, CTR/DRE, CuRE, MYB, MYC, TAAG-box, W-box, WRKY | ARR1, NTBBF |
| *TaTLP8-A* | BOXC, BOXL, GT1CONSENSUS,  GATA-box, PRE, SORLIP | CACT, DOFCORE, E-box, POLLEN1 | ACGT, ASF1, BOX-A, BIHD1, CuRE, GCC-box, MYB, MYC, SEBF, SAUR, TAAAG-box, WRKY | ABRE, ARR1, CGCG-box, DPBF |
| *TaTLP8-B* | BOX II, GT1CONSENSUS,  GATA-box, SORLIP | ANAERO, CACT, DOFCORE, E-box, NAPIN, POLLEN1, Q-ELEMENT | ACGT, BIHD1, CuRE, GCC-box, IRO, MYB, MYC, P1BS, SURE, W-box, WRKY | ABRE, ARR1, CGCG-box, DPBF, EMBP |
| *TaTLP8-D* | BOXC, BOXL, GT1CONSENSUS,  GATA-box, I-box, PRE, SORLIP, T-box | ANAERO, CACT, DOFCORE, E-box, POLLEN1, RY-ELEMENTS | ACGT, BOX-A, CuRE, GCC-box, MYB, MYC, QAR, T/G-box, TAAAG-box, UPR | ABRE, ARR1, CARE, CGCG-box, DPBG |
| *TaTLP9-A* | BOXL, GT1CONSENSUS,  GATA-box, SORLIP | CACT, CAN, DOFCORE, E-box, POLLEN1 | ACGT, ASF1, BOX-A, CTR/DRE, DRE, GCC-box, LTRE, MYB, MYC, SURE, W-box, WRKY | ABRE, ARR1, AUX, CGCG-box |
| *TaTLP9-D* | BOXL, GT1CONSENSUS,  GATA-box, I-box, PRE, SORLIP, T-box | ANAERO, CACT, CAN, CIACADIAN, DOFCORE, E-box, POLLEN1, Q-ELEMENT, SRE | ACGT, ARE, BIHD1, BOX-A, CTR/DRE, CuRE, DRE, ELRE, GCC-box, LTRE, MYB, MYC, SURE, TAAAG-box, UPR, W-box, WRKY | ABRE, ARR1, CARE, CGCG-box, DPBF, ERE, NTBBF |
| *TaTLP10-A* | BOXC, GT1CONSENSUS,  GATA-box, I-box, RBCS, SORLIP | CACT, DOFCORE, E-box, POLLEN1, Q-ELEMENT, SRE | ACGT, ASF1, BIHD1, CuRE, DPBF, DRE, MYB, MYC, SURE, TAAA-box, W-box, WRKY | ARF, ARR1, CGCG-box, DPBF, |
| *TaTLP10-B* | BOXC, GT1CONSENSUS,  GATA-box, I-box, PRE, RBCS, SORLIP | ANAERO, CACT, CIACADIAN, DOFCORE, E-box, POLLEN1, RY-ELEMENTS, SREA | ACGT, ASF1, BIHD1, CTR/DRE, CuRE, DRE, LTRE, MYB, MYC, SEBF, SURE, TAAA-box, W-box, WRKY | ARF, ARR1, CARE, CGCG-box, DPBF, NTBBF |
| *TaTLP10-D* | BOXC, GT1CONSENSUS,  GATA-box, I-box, PRE, RBCS, SORLIP, T-box | NAERO, CACT, CEREGLUE-box, CIACADIAN, DOFCORE, E-box, POLLEN1, Q-ELEMENT, RY-ELEMENTS, SRE | ACGT, ASF1, BIHD1, CTR/DRE, CuRE, DRE, IRO, LTRE, MYB, MYC, SEBF, SURE, TAAA-box, W-box, WRKY | ABRE, ARF, ARR1, CGCG-box, CARE, DPBF, GARE, NTBBF |
| *TaTLP11-A* | GT1CONSENSUS,  GATA-box, I-box, PRE, T-box | ANAERO, CACT, CIACADIAN, DOFCORE, E-box, POLLEN1, Q-ELEMENT, RY-ELEMENTS, SRE | ACGT, BIHD1, CTR/DRE, CuRE, DRE, LTRE, MYB, MYC, T/G-box, TAAAG-box, W-box, WRKY | ABRE, ARR1, DPBF |
| *TaTLP11-B* | GT1CONSENSUS,  GATA-box, I-box, PRE | ANAERO, CACT, DOFCORE, E-box, HDZIP, POLLEN1, SRE | ACGT, BIHD1, CTR/DRE, CuRE, DRE, LTRE, MYB, MYC, TAAAG-box, W-box, WRKY | ARR1. DPBF, GARE |
| *TaTLP11-D* | GT1CONSENSUS,  GATA-box, I-box, PRE, T-box | CACT, CIACADIAN, DOFCORE, E-box, POLLEN1, RY-ELEMENTS, SRE | ACGT, BIHD1, CTR/DRE, CuRE, DRE, LTRE, MYB, MYC, TAAAG-box, SURE, W-box, WRKY | ABRE, ARF, ARR1, CGCG-box, DPBF, |
| *TaTLP12-A* | GT1CONSENSUS,  GATA-box, I-box, SORLIP | ANAERO, CACT, CIACADIAN, DOFCORE, E-box, RY-ELEMENTS | ACGT, ASF1, BIHD1, CTR/DRE, CuRE, DRE, ELRE, LTRE, MYB, MYC, QAR, SURE, T/G-box, TAAAG-box, -box, WRKY | ABRE, ARF, ARR1, DPBF, ERE, |
| *TaTLP12-B* | GT1CONSENSUS,  GATA-box, SORLIP, T-box | ANAERO, CACT, DOFCORE, E-box, POLLEN1, | ASF1, BIHD1, CTR/DRE, CuRE, GCC-box, DRE, MYB, MYC, SEB, SURE, T/G-box, TAAAG-box, W-box, WRKY | ARF, ARR1, CGCG-box, DPBF, |
| *TaTLP12-D* | GT1CONSENSUS,  PRE, SORLIP, T-box | ANAERO, CACT, COFCORE, E-box, POLLEN1, | ACGT, ASF1, BIHD1, CTR/DRE, CuRE, DRE, LTRE, MYB, MYC, SURE, W-box, WRKY | ABRE, ARF, ARR1, CARE, CGCG-box, DPBF |
| *TaTLP13-D* | GT1CONSENSUS,  GATA-box, I-box, PRE, SORLIP | CACT, CAN, DOFCORE, E-box, POLLEN1, Q-ELEMENT, RY-ELEMENTS | ACGT, ASF1, BIHD1, CuRE, ELRE, MYB, MYC, PIBS, T/G-box, W-box, WRKY | ABRE, ARR1 |
| *TaTLP14-A1* | BOX II, GT1CONSENSUS,  GATA-box, I-box, PRE, RBCS, SORLIP | ANAERO, CACT, CAN, DOFCORE, E-box, POLLEN1, RY-ELEMENTS | ACGT, ASF1, BIHD1, BOX-A, CuRE, MYB, MYC, P1BS, T/G-box, W-box, WRKY | ABRE, ARR1, CGCG-box, DPBF, NTBBF |
| *TaTLP14-A2* | BOX II, GT1CONSENSUS,  GATA-box, I-box, PRE, SORLIP | ANAERO, CACT, DOFCORE, E-box, POLLEN1, RY-ELEMENTS | ACGT, BIHD1, CTR/DRE, DRE, LTRE, MYB, MYC, W-box, WRKY | ABRE, ARR1, CARE, CGCG-box, CPB, DPBF, |
| *TaTLP14-B1* | GT1CONSENSUS,  GATA-box, I-box, RBCS, SORLIP | ANAERO, CACT, CAN, DOFCORE, E-box, POLLEN1, RY-ELEMENTS | ACGT, ASF1, BIHD1, CuRE, ELRE, LTRE, MYB, MYC, PIBS, SURE, W-box, WRKY | ARR1, CPB, DPBF, |
| *TaTLP14-B2* | BOX L, GT1CONSENSUS,  GATA-box, SORLIP | ANAERO, BOX-B, CACT, DOFCORE, E-box, POLLEN1, Q-ELEMENT | ACGT, ASF1, BIHD1, CuRE, CRT/DRE, ELRE, GCC-box, MYB, MYC, PIBS, SURE, TAAAG-box, W-box, WRKY | ABRE, ARR1, CGCG-box, DPBF, |
| *TaTLP14-B3* | BOX II, GATA-box, SORLIP, T-box | CACT, CIACADIAN, DOFCORE, E-box, LEAFY, POLLEN, Q-ELEMENT, RY-ELEMENTS, | ACGT, ASF1, BIHD1, CuRE, LTRE, MYB, MYC, SURE, TAAAG-box, W-box, WRKY | ABRE, ARR1, AUX, CARE, CGCG-box, DPBF, |
| *TaTLP14-B4* | BOX II, GT1CONSENSUS,  GATA-box, I-box, SORLIP | ANAERO, CACT, DOFCORE, POLLEN1, RY-ELEMENTS, SRE | ACGT, BIHD1, CuRE, GCC-box, MYB, MYC, SEBF, SURE, TAAAG-box, WRKY | ABRE, ARR1, CGCG-box |
| *TaTLP14-B5* | GT1CONSENSUS,  GATA-box, I-box, PRE, RBCS, SORLIP | ANAERO, CACT, CAN, DOFCORE, E-box, RY-ELEMENTS | ACGT, BIHD1, BOX-A, CTR/DRE, CuRE, DRE, ELRE, LTRE, MYB, MYC, TAAAG-box, W-box, WRKY | ARR1, CARE, CGCG-box, DPBF, NDE, |
| *TaTLP15-A* | GATA box, GT1CONSENSUS, I-box, PRE, RBCS, SORLIP, T BOX | ANAERO, CACT, CIACADIAN, DOFCORE, E-box, EVE, GTG, POLLEN1, | ACGT, ASF1, BIHD1, CGAC, CuRE, ELRE, MYB, MYC, SURE, WBOX, WRKY | ARR1, ERE |
| *TaTLP15-B1* | GATA, GT1CONSENSUS, SORLIP | ANAERO, CACT, DOFCORE, E-box, GCN, GLMH, GTGA, POLLEN1, | CGAC, MYC, WBOX, WRKY | ARR1, CPB, DPBF |
| *TaTLP15-B2* | GT1COSENSUS, PRE, SORLIP | CACT, CIACADIAN, DOFCORE, E-box, GTGA, RY elements, WUSAT | ACGT, ASF1, BIHD1, CuRE, GCC, MYC, SEBF, SURE, TAAAG-BOX, WBOX, WRKY | ARR1, CARE, DPBF, GRAZ |
| *TaTLP15-B3* | GT1CONSENSUS,  GATA-box, I-box, PRE, SORLIP | ANAERO, CACT, CIACADIAN, DOFCORE, EVE, GTGA, RY elements, S1F, | ACGT, ASF1, BIHD1, CuRE, EECC, LTRE, MYB, SURE, TAAAG-BOX, W-box, WRKY | ARR1, NTBBF |
| *TaTLP16-A* | GT1CONSENSUS, GATA, RBCS, SORLIP | CACT, DOFCORE, E-box, GTGA, POLLEN1, RY-elements, TGAC, UP1 | ACGT, ASF1, CACG, CTR/DRE, CGAC, CRTD, CuRE, IRO, LTRE, MYB, MYC, W-box, WRKY | ABRE, ARR1, ERE, GARE, TATC |
| *TaTLP17-A* | GT1CONSENSUS,  GATA-box, I-box, PRE, SORLIP | ANAERO, CACT, DOFCORE, E2F, E-box, GTGA, POLLEN1, RHER, SREA | ACGT, BIHD1, CGAC, CuRE, GCC, LTRE, MYB, MYC, SEBF, TAAAG-BOX, WBOX, WRKY | ABRE, ARR1, CATA, CGCG-BOX-BOX, |
| *TaTLP18-A* | BOX II, GATA, GT1CONSENSUS, I-box, SORLIP, T BOX | ANAERO, CACT, CIADIAN, DOFCORE, E2F, E-box, GTGA, POLLEN1, , TGAC | ACGT, ASF1, MYB, MYC, WBOX, WRKY | ARR1, CPB |
| *TaTLP18-B* | BOX II, GATA, GT1CONSENSUS, I-box, SORLIP, T BOX | CACT, CIADIAN, DOFCORE, E2F, E-box, GTGA, POLLEN1, , RY elements, , , , TE2F, TGAC | ACGT, ASF1, CuRE, MYB, MYC, P1BS, TAAAG-BOX, WBOX, WRKY | ARR1, CPB, TATC |
| *TaTLP19-A* | GATA, GT1CONSENSUS, I-box, SORLIP, T BOX | ANAERO, CACT, DOFCORE, E-box, GTGA, POLLEN1, RY elements | ACGT, BIHD1, CuRE, MYB, MYC, TAAAG-BOX, UPR, WBOX, WRKY | ARR1, CATA, GARE, |
| *TaTLP20-A1* | BOXL, GT1CONSENSUS,  GATA-box, I-box SORLIP, T BOX | CACT, DOFCORE, E2F, GTGA, POLLEN1, RHER, RY elements, S1F | BIHD1, CTR/DRE, CuRE, EECC, LTRE, MYB, W-box. WRKY | ARR1, CPB, DPBF, ERE, |
| *TaTLP20-A2* | BOX C, BOX II, BOX L, GATA, IBOX, PRE, T BOX | ANAERO, CACT, CIACADIAN, DOFCORE, E-box, GTGA, POLLEN1, RHER, SREA | BIHD1, CTR/DRE, CGAC, DRE, EECC, LTRE, MYB, MYC, WBOX, WRKY | , ARR1, CATA, CGCG-BOX-BOX, CPB, DPBF, ERE, GARE, TATC |
| *TaTLP20-A3* | BOXL, GT1CONSENSUS, GATA-box, I-box, T BOX | ANAERO, CACT, CIACADIAN, DOFCORE, E-box, GTGA, POLLEN1, RHER, SREA | ACGT, BIHD1, CTR/DRE, DRE, LTRE, MYB, MYC, UPR, W-box, WRKY | , ARR1, CPB, DPBF, ERE, GARE, |
| *TaTLP20-A4* | BOXII, BOXL, GT1CONSENSUS, GATA-box, I-box, T BOX | ANAERO, CACT, CIACADIAN, DOFCORE, E-box, GTGA, POLLEN1, RHER, SREA | BIHD1, CTR/DRE, DRE, LTRE, MYB, MYC, W-box, WRKY | ABRE, ARR1, CATA, CGCG-BOX-BOX, CPB, DPBF, ERE, GARE, |
| *TaTLP20-A5* | BOX C, BOX II, BOX L, GATA-box, I-box, PRE, T BOX | ANAERO, CACT, CIACADIAN, DOFCORE, E-box, GTGA, POLLEN1, RHER, SREA | BIHD1, CTR/DRE, CGAC, DRE, LTRE, MYB, MYC, W-box, WRKY | , ARR1, CATA, CGCG-BOX-BOX, CPB, DPBF, ERE, GARE, TATC |
| *TaTLP20-A6* | BOX C, BOX L, GATA, GT1CONSENSUS, I-box, T BOX | AACA, ANAERO, CACT, CIACADIAN, DOFCORE, E-box, GTGA, POLLEN1, RHER, SREA | ACGT, BIHD1, CTR/DRE, CGAC, LTRE, MYB, MYC, W-box, WRKY | , ARR1, CATA, CGCG-BOX-BOX, CPB, DPBF, ERE, GARE, |
| *TaTLP21-A* | GATA, GT1CONSENSUS, I-box, PRE, SORLIP, SV40 | ANAERO, CACT, DOFCORE, E-box, GTGA, POLLEN1, RHER, RY elements, | ASF1, BIHD1, CTR/DRE, CuRE, DRE, LTRE, MYB, MYC, SURE, TAAAG-BOX, W-box, WRKY | ARR1, DPBF, |
| *TaTLP22-A* | , PRE, SORLIP, SV40 | ANAERO, CACT, DOFCORE, E-box, GTGA, RY elements, , XYLAT | ACGT, CTR/DRE, CuRE, ELRE, GCC, LTRE, MYB, MYC, SURE, TAAAG-BOX, W-box, WRKY | ABRE, ARR1, CGCG-BOX-BOX, GARE |
| *TaTLP22-B* | GATA, PRE, SORLIP | CACT, DOFCORE, E-box, GTGA, POLLEN1, Q ELEMENT, RHER, RY elements, , XYLAT | ACGT, CTR/DRE, CGAC, DRE, EECC, ELRE, GCC, LTRE, MYB, MYC, SURE, TAAAG-BOX, W-box, WRKY | ABRE, ARR1, CGCG-BOX-BOX, DPBF, GARE |
| *TaTLP22-C* | BOX L, PIAT, SORLIP, SV40 | ANAERO, CACT, DOFCORE, E-box, GTGA, POLLEN1, RHER, XYLAT | ACGT, CGAC, CuRE, DRE, ELRE, LTRE, MYB, MYC, SURE, TAAAG-BOX, W-box, WRKY | ARR1, CGCG-BOX-BOX, GARE |
| *TaTLP23-A* | BOX L, GATA, GT1CONSENSUS, I-box, PRE, SV40 | CACT, CIACADIAN, DOFCORE, E-box, GTGA, L1 BOX, POLLEN1, Q ELEMENT, RHER | ACGT, ASF1, CTR/DRE, CGA, CuRE, DRE, EECC, LTRE, MYB, MYC, TAAAG-BOX, W-box, WRKY | ABRE, ARR1, CARE, NTBBF, , TATC |
| *TaTLP23-B* | BOX L, GATA, GT1CONSENSUS, I-box, SV40, T BOX | AACA, CACT, CIACADIAN, DOFCORE, E-box, GTGA, POLLEN1, RHER, RY elements, SREA, TGAC | ACGT, ASF1, BIHD1, CTR/DRE, CuRE, DRE, EECC, LTRE, MYB, MYC, SURE, W-box, WRKY | ABRE, ARR1, CARE, CGCG-BOX-BOX, GARE, TATC |
| *TaTLP23-D* | GATA, GT1CONSENSUS, I-box, PRE, SORLIP, SV40 | CACT, CIACADIAN, DOFCORE, E2F, E-box, GTGA, POLLEN1, RHER, RY elements, SREA | ACGT, ASF1, BIHD1, CTR/DRE, CGA, CRTD, CuRE, DRE, EECC, LTRE, MYB, MYC, SURE, TAAAG-BOX, W-box, WRKY | ABRE, ARF, ARR1, CARE, CGCG-BOX-BOX, NTBBF, TATC |
| *TaTLP24-A* | BOX L, GATA, GT1CONSENSUS, SORLIP | ANAERO, CACT, DOFCORE, E-box, GTGA, POLLEN1, RHER, RY elements, S1F, | ACGT, BIHD1, CuRE, LTRE, MARABOX, MYB, MYC, SURE, TAAAG-BOX, W-box, WRKY | ABRE, ARR1, CATA, CGCG-BOX-BOX, DPBF, NTBBF |
| *TaTLP24-B* | GATA, GT1CONSENSUS, PRE, SORLIP | ANAERO, CACT, DOFCORE, GTGA, POLLEN1, RHER, RY elements, | ACGT, ASF1, BIHD1, CTR/DRE, CGAC, CRTD, CuRE, DRE, EECC, LTRE, MARABOX, MYB, SURE, W-box, WRKY | ABRE, ARR1, CARE, CGCG-BOX-BOX, DPBF |
| *TaTLP24-D* | BOX L, GATA, GT1CONSENSUS, SORLIP | ANAERO, CACT, DOFCORE, E-box, GTGA, POLLEN1, Q ELEMENT, RHER, RY elements, | ACGT, ASF1, BIHD1, CTR/DRE, DRE, EECC, LTRE, MYB, MYC, SURE, W-box, WRKY | ABRE, ARR1, CATA, CGCG-BOX-BOX, DPBF, NTBBF |
| *TaTLP25-A* | BOX II, GATA, GT1CONSENSUS, I-box, PRE, SORLIP, SV40 | ANAERO, CACT, DOFCORE, E2F, E-box, EVE, GTGA, POLLEN1, RHER, S1F, SEF1, TGAC | ACGT, ASF1, CACG, CuRE, EECC, GCC, IRO, MYB, MYC, SURE, UPR, WRKY | ABRE, ARR1, CACG, DPBF, GADOWN, TATC |
| *TaTLP25-B* | BOX C, BOX II, GATA, GT1CONSENSUS, I-box, LREN, PRE, SORLIP | ANAERO, CACT, CIACADIAN, DOFCORE, E2F, E-box, EVE, GTGA, POLLEN1, RHER, S1F, TGAC | ACGT, ASF1, BIHD1, CACG, CGAC, CuRE, EECC, IRO, MYB, MYC, TAAAG-BOX, WRKY | ABRE, ARR1, CARE, DPBF, EMBP, GADOWN, |
| *TaTLP25-D* | BOX C, BOX II, GATA, GT1CONSENSUS, I-box, LREN, PRE, SORLIP | ANAERO, CACT, CAN, CIACADIAN, DOFCORE, E2F, E-box, EVE, GTGA, POLLEN1, RHER, S1F, SEF1, TGAC | ACGT, ASF1, CACG, CGAC, CuRE, EECC, IRO, LTRE, MYB, MYC, SURE, WBOX, WRKY | ABRE, ARF, ARR1, CARE, DPBF, EMBP, GADOWN |
| *TaTLP26-B* | GATA, GT1CONSENSUS, I-box, SORLIP | AACA, ANAERO, CACT, CAN, CIACADIAN, DOFCORE, E-box, GTGA, POLLEN1, | ASF1, BIHD1, EECC, MARABOX, MYB, MYC, SEBF, SURE, TAAAG-BOX, WBOX, WRKY | AMYBOX, ARF, ARR1, CATA, CPB, DPBF, GARE, NTBBF, |
| *TaTLP26-D* | BOX L, GATA, GT1CONSENSUS, I-box, PRE, SORLIP | CACT, CIACADIAN, DOFCORE, E2F, E-box, GTGA, POLLEN1, RY elements | ACGT, BIHD1, CTR/DRE, CGAC, CuRE, DRE, ELRE, LTRE, MYB, MYC, P1BS, SURE, TAAAG-BOX, WBOX, WRKY | ABRE, ARR1, DPBF, TATC |
| *TaTLP27-A* | BOX II, GATA, GT1CONSENSUS, I-box, SORLIP | ANAERO, CACT, DOFCORE, E-box, GTGA, POLLEN1, RHER, SEF1, WUSAT | ACGT, BIHD1, CACG, CTR/DRE, CGAC, CRTD, CuRE, EECC, MYB, MYC, SURE, TAAAG-BOX, WBOX, WRKY | ABRE, AMYBOX, ARR1, CATA, CGCG-BOX-BOX, DPBF, SBOX, TATC |
| *TaTLP27-B* | BOX L, GATA, GT1CONSENSUS, PRE, SORLIP, T BOX | ANAERO, CACT, DOFCORE, E-box, GTGA, POLLEN1, RHER, RY elements, SEF1, , WUSAT | ACGT, CTR/DRE, CGAC, CuRE, DRE, EECC, GCC, LTRE, MYB, MYC | ABRE, ARR1, CATA, CGCG-BOX-BOX, DPBF, SBOX |
| *TaTLP27-D* | GATA, GT1CONSENSUS, PRE, SORLIP | ANAERO, CACT, DOFCORE, E-box, GTGA, POLLEN1, SEF1 | ACGT, BIHD1, CGAC, CuRE, GCC, MYB, MYC, SURE, TAAAG-BOX, WBOX, WRKY | ABRE, ACGT, AGC, ARR1, CGCG-BOX-BOX, DPBF, GADOWN, |
| *TaTLP28-A* | BOX II, BOX L, GATA, GT1CONSENSUS, SORLIP | ANAERO, CACT, CIACADIAN, DOFCORE, E-box, GCN, GTGA, POLLEN1, , RHER, RY elements, S1F, | BIHD1, CuRE, EECC, ELRE, MYB, MYC, SEBF, TAAAG-BOX, WBOX, WRKY | ABRE, ARR1, AUX, CARE, CPB, DPBF, NTBBF, TATC |
| *TaTLP28-B* | GATA, GT1CONSENSUS, RBCS, SORLIP | ANAERO, CACT, CIACADIAN, DOFCORE, E2F, E-box, GCN, GTGA, POLLEN1, RY elements, S1F, | ASF1, BIHD1, CuRE, EECC, ELRE, MYB, MYC, SEBF, TAAAG-BOX, WBOX, WRKY | ARR1, DPBF |
| *TaTLP28-D* | BOX II, BOX L, GATA, GT1CONSENSUS, I-box, RBCS, SORLIP | ANAERO, CACT, CIACADIAN, DOFCORE, E-box, GTGA, POLLEN1, RY elements, S1F, | BIHD1, CuRE, EECC, ELRE, MYB, MYC, SEBF, TAAAG-BOX, WBOX, WRKY | ARR1, CARE, DPBF, NTBBF |
| *TaTLP29-A* | GT1CONSENSUS, PRE, SORLIP, SV 40 | CACT, CAN, DOFCORE, E2F, E-box, GTGA, POLLEN1, RHER, , | ACGT, ASF1, BIHD1, CTR/DRE, CGAC, DRE, EECC, ELRE, GCC, LTRE, MYB, MYC, SEBF, SURE, WBOX, WRKY | ARR1, AUX, CGCG-BOX-BOX, DPBF, PROX, |
| *TaTLP29-B* | BOX L, GATA, GT1CONSENSUS, SORLIP | CACT, DOFCORE, E2F, E-box, GTGA, POLLEN1, Q ELEMENT, RHER, , | ACGT, BIHD1, CACG, CGAC, CURE, GCC, LTRE, MYB, MYC, TAAAG-BOX, WBOX, WRKY | ABRE, ARR1, CGCG-BOX-BOX, DPBF, NTBBF, |
| *TaTLP29-D* | BOX C, BOX L, GT1CONSENSUS, PRE, SORLIP, T BOX | CACT, CAN, DOFCORE, E2F, E-box, GTGA, POLLEN1, , RHER, | BIHD1, CTR/DRE, CuRE, DRE, EECC, ELRE, GCC, LTRE, MYB, MYC, SURE, WBOX, WRKY | ARR1, CARE, CGCG-BOX-BOX, DPBF, PROX, |
| *TaTLP30-A* | BOX L, GATA, GT1CONSENSUS, RBCS, PRE, SORLIP, SV 40 | CACT, CAN, DOFCORE, E2F, E-box, GTGA, POLLEN1, RY elements, | ACGT, BIHD1, BP5O, CTR/DRE, CuRE, DRE, EECC, LTRE, MYB, MYC, QARB, SEBF, SURE, T/G BOX, TAAAG-BOX, WBOX, WRKY | ABRE, ARF, ARR1, CGCG-BOX-BOX, DPBF, GADOWN, NTBBF, |
| *TaTLP30-B* | BOX C, BOX II, BOX L, GATA, GT1CONSENSUS, RBCS, PRE, SORLIP, SV 40 | ANAERO, CACT, CAN, DOFCORE, E-box, GTGA, POLLEN1, RHER, RY elements, , | ACGT, ASF1, CACG, BIHD1, BP5O, CTR/DRE, CGAC, CuRE, DRE, IRO, LTRE, MYB, MYC, QARB, SEBF, SURE, T/G BOX, TAAAG-BOX, UPR, WBOX, WRKY | ABRE, ARF, ARR1, CARE, CGCG-BOX-BOX, DPBF, EMBP, NTBBF, , TATC |
| *TaTLP30-D* | BOX L, GATA, GT1CONSENSUS, PRE, SORLIP, SV 40 | ANAERO, CACT, CERE, DOFCORE, E-box, GTGA, POLLEN1, RY elements, , , | ACGT, ASF1, BIHD1, BP5O, CuRE, EECC, MYB, MYC, QARB, T/G BOX, WBOX, WRKY | ABRE, ARR1, CGCG-BOX-BOX, DPBF, GADOWN |
| *TaTLP31-A* | GATA, GT1CONSENSUS, I-box, SORLIP, SV 40, T BOX | CACT, DOFCORE, E-box, GTGA, POLLEN1, Q ELEMENT, , RY elements, | ACGT, ASF1, BIHD1, CuRE, EECC, MYB, MYC, SURE, WBOX, WRKY | ABRE, ARR1, CARE |
| *TaTLP31-B* | GT1CONSENSUS, RBCS, SORLIP, T BOX | ANAERO, CACT, CIACADIAN, DOFCORE, GCN, GTGA, POLLEN1, , RY elements, SEF1, , , , SPH | BIHD1, CTR/DRE, DRE, EECC, LTRE, MYB, SURE, TAAAG-BOX, WBOX, WRKY | ARR1, CARE, CGCG-BOX-BOX, |
| *TaTLP31-D1* | GT1CONSENSUS, RBCS, SORLIP, T BOX | ROOTMOTIF, ANAERO, CACT, E-box, GCN, GTGA, POLLEN1, , RHER, RY elements, , SPH | BIHD1, CuRE, EECC, MYB, MYC, SURE, WBOX, WRKY | ARR1, CARE, ERE |
| *TaTLP31-D2* | BOX II, CPR, GATA, G BOX, LRE BOX, GT1CONSENSUS, SORLIP, SV40, T BOX | CACT, DOFCORE, E-box, GTGA, POLLEN1, RHER, RY elements, SEF 4, | ACGT, BIHD1, CACG, EECC, IRO, MYB, MYC, SURE, TAAAG-BOX, WBOX, WRKY | ABRE, ARR1, CPB, EMBP, |
| *TaTLP32-A* | GATA, GT1CONSENSUS, I-box, SORLIP, T BOX | ANAERO, CACT, DOFCORE, E2F, E-box, GTGA, POLLEN1, RY elements, SREA | EECC, ELRE, MYB, MYC, TAAAG-BOX, WBOX, WRKY | , ARR1, DPBF, GARE |
| *TaTLP32-B* | GATA, GT1CONSENSUS, I-box, SORLIP, T BOX | ANAERO, CACT, CAN, DOFCORE, E2F, E-box, GTGA, POLLEN1, , RHER, | ACGT, CuRE, EECC, ELRE, MYB, MYC, WBOX, WRKY | ABRE, ARR1, CACG, GARE |
| *TaTLP32-D* | GATA, GT1CONSENSUS, I-box, RBCS, SORLIP, T BOX | ANAERO, CACT, CAN, DOFCORE, E-box, GTGA, POLLEN1, , RHER | ACGT, CuRE, EECC, ELRE, MYB, MYC, TAAAG-BOX, WBOX, WRKY | ABRE, , ARR1, D4GM, GARE |
| *Z. mays* | | | | |
| *ZmTLP1* | GATA box, GT1CONSENSUS, I-box, PRE, INR, SORLIP, BOXL | DOFCORE, E-box, POLLEN1, ANAERO, CIACADIAN, RY elements | ACGT, ASF1, BIHD1, DRE, CuRE, MYB, MYC, W-box, WRKY, P1BS, SURE, TAAG | ABRE, ARR1, ARF, ERE, NTBBF, DPBF |
| *ZmTLP2* | BOXL, GATA box, GT1CONSENSUS, INR, PRE, SORLIP, T-box | ANAERO, CACT, CIACADIAN, DOFCORE, E-box, EVENING, GTGA, Q ELEMENT | ACGT, ASF1, CRT/DRE, DRE, ELRE, LTRE, MYB, MYC, TAAAG-BOX, UPR, W-box, WRKY | ABRE, ARR1, CGCG-BOX-BOX, DPBF, NTBBF |
| *ZmTLP3* | BOXL, GATA box, GT1CONSENSUS, I-box, PRE, SORLIP | ANAERO, CACT, DOFCORE, E-box, HDZIP, POLLEN1 | ASF-1, BIHD1, CGAC, CuRE, MYB, MYC, PREAT, SEBF, SURE, TAAAG-BOX, W-box, WRKY | ABRE, ARF, CGCG-BOX-BOX-box, ARR1, DPBF, NTBBF, |
| *ZmTLP4* | GT1CONSENSUS,  GATA-box, I-box, PRE, SORLIP, SV40, T-box | ANAERO, CACT, DOFCORE, E-box, POLLEN1 | ACGT, ASF-1, BIHD1, CuRE, EECC, ELRE, MYB, MYC, SURE, TAAAG-BOX, W-box, WRKY | ABRE, ARR1, AUX, CGCG-BOX-BOX-box, CPB, NTBBF |
| *ZmTLP5* | BOX-II, GT1CONSENSUS, GATA-box, I-box, SORLIP | ANAERO, CACT, DOFCORE, E-box, POLLEN1, RHER, RY-elements | ACGT, CTR/DRE CuRE, MYB, MYC, SURE, TAAAG-BOX, W-box, WRKY | ABRE, ARR1, CARE, NTBBF, |
| *ZmTLP6* | BOX-II, GT1,  GATA-box, I-box, SORLIP | CACT, , DOFCORE E2F, POLLEN1, Q ELEMENT, RY-elements | ACGT, BIHD1, CGAC, CuRE, EECC R, GCC-box, LTRE, SEBF, TAAAG-BOX, MYB, W-box, WRKY | ABRE, ARR1, CGCG-BOX-BOX, CPB, DPBF, NTBBF, ARF |
| *ZmTLP7* | BOX-II, BOXL, GT1,  GATA-box, RBCS, SORLIP | CACT, DOFCORE, E2F, POLLEN1, Q ELEMENT, RY-elements | ACGT, BIHD1, CuRE, GCC-box, LTRE, MYB, W-box, WRKY | ABRE, ARR1, CGCG-BOX-BOX-box, CPB, DPBF, GARE |
| *ZmTLP8* | GT1CONSENSUS,  GATA-box, I-box, PRE, SORLIP, T-box | ANAERO, CACT, CIACADIAN, DOFCORE, E-box, POLLEN1, RY-elements, SRE | ACGT, ASF-1, CTR/DRE, CuRE, DRE, ELRE, LTRE. MYB, MYC, TAAAG-BOX, W-box, WRKY | ARR1, CATA, CGCG-BOX-BOX-box, CPB, DPBF, GARE |
| *ZmTLP9* | BOX-II, GT1CONSENSUS, GATA-box, I-box, PRE, SORLIP | CACT, DOFCORE, E-BOX, POLLEN 1, | ACGT, ASF1, BIHD1, BOX-A, CRT/DRE, CuRE, DRE, GCC-box, LTRE, MYB, MYC, SURE, T/G box, TAAAG-BOX, W-box, WRKY | ABRE, ARR1, ERE, S-BOX |
| *ZmTLP10* | GT1CONSENSUS,  GATA-box, I-box, PRE, SORLIP, T-box | CACT, DOFCORE, E2F, POLLEN1, Q ELEMENT, SRE, XYLAT | ACGT, ARE1, ASF1, BIHD1, MYB, MYC, SURE, TAAAG-BOX, W-box. WRKY | ABRE, ARR1, CGCG-BOX-BOX-box, DPBF, GARE |
| *ZmTLP11* | GT1CONSENSUS, GATA-box, | ANAERO, CACT, CIACADIAN, DOFCORE, E-box, POLLEN1, Q ELEMENT | ACGT, ELRE, MYB, MYC, W-box, WRKY | ARR1, CPB |
| *ZmTLP12* | BOXL, GT1CONSENSUS,  GATA-box, I-box, PRE, SORLIP | ANAERO, CACT, DOFCORE, E-box, L1BOX, SREAT | ACGT, ASF1, A-box, BIHD1, CRT/DRE, CuRE, DRE, GCC-box, LTRE, MYB, MYC, TAAAG-BOX, W-box, WRKY | ABRE, ARR1, CGCG-BOX-BOX-box |
| *ZmTLP13* | BOXII, GT1CONSENSUS,  GATA-box, SV40 | CACT, DOFCORE, E-box, LEAFY, POLLEN1, | ACGT, BIHD1, CTR/DRE, CuRE, EECC, GARE, GCC, LTRE, MYB, MYC, TAAAG-BOX, UPR, W-box, WRKY | ABRE, ARR1, CGCG-BOX-BOX-box, CPB, ERE |
| *ZmTLP14* | GT1CONSENSUS,  GATA-box, I-box, PRE, SORLIP, T-box | CACT, CAN, DOFCORE, E-box, HDZIP, POLLEN1 | ACGT, ASF1, BP5, CuRE, EECC, MYB, MYC, P1BS, T/G box, TAAAG-BOX, WRKY | ABRE, ARR1, CARE, CPB |
| *ZmTLP15* | BOX L, GT1CONSENSUS,  GATA-box, I-box, T-box | ANAERO, CACT, CAN, CIACADIAN, DOFCORE, E2F, POLLEN1, Q ELEMENT, SRE | ACGT, BIHD1, CTR/DRE, CGAC, CuRE, DRE, ELRE, LTRE, MARABOX, MYB, MYC, QARB, SURE, T/G BOX, WBOX, WRKY | ABRE, ARR1, CACG, DPBF, AUX, GARE |
| *ZmTLP16* | BOX L, GT1CONSENUS | ANAERO, E2F, | ACGT, ASF1, CTR/DRE, CGAC, DRE, GCC, LTRE, MYB, SURE, UPR, WBOX, WRKY | ABRE, ARR1, CGCG-BOX-BOX |
| *ZmTLP17* | BOXII, GATA, GT1CONSENSUS, INRN, SORLIP | ANAERO, CACT, E2F, E BOX, NAP, DOFCORE, | ACGT, ASF1, BIHD1, CuRE, EECC, GCC, LTRE, MYB, MYC, TAAAG-BOX, WRKY | ABRE, AMYBOX, ARR1, DPBF, GARE, |
| *ZmTLP18* | GT1CONSENSUS,  GATA-box, I-box, PRE, SV40 | AACA, ANAERO, CACT, DOFCARE, E-box, GCN, GLM, L1 BOX, POLLEN1, Q ELEMENT, , RY elements, S1F, , TGAC | ACGT, ASF1, BIHD1, CGAC, CuRE, MARABOX, MYB, MYC, P1BS, TAAAG-BOX, WBOX, WRKY | ABRE, ARR1, CATA, DPBF, GARE, |
| *ZmTLP19* | GATA, GT1CONSENSUS, INRN, PRE, SORLIP, SV40 | ANAERO, CAC T, DOFCORE, E2F, E-box, POLLEN1, Q ELEMENT, TGAC | ACGT, ASF1, BIHD1, CTR/DRE, CGAC, CuRE, GCC, LTRE, MYB, MYC, SURE, WBOX, WRKY | ARR1, CGCG-BOX-BOX, DPBF, SBOX |
| *ZmTLP20* | BOX L, GATA, GT1CONSENSUS, HBOX, I-box, PRE, SORLIP | ANAERO, CACT, DOFCORE, E2F, E-box, L1 BOX, POLLEN1, S1F, SEF1, | ACGT, ASF1, CuRE, EECC, MYB, MYC, TAAAG-BOX, UPR, WBOX, WRKY | AMYBOX, ARR1, CGCG-BOX-BOX, DPBF, GARE, NTBBF, |
| *ZmTLP21* | BOX L, GATA, GT1CONSENSUS, I-box, INRN, PRE, SORLIP | CACT, DOFCORE, E2F, E-box, POLLEN1, RY elements, S1F, | EECC, LTRE, MYB, MYC, SURE, TAAAG-BOX, WBOX, WRKY | ARR1, CARE, CPB, DPBF, TATC |
| *ZmTLP22* | BOX II, GATA, GT1CONSENSUS, I-box, INRN, PRE | ANAERO, CACT, CAN, CIACADIAN, DOFCORE, E-box, POLLEN1, S1F, | ASF1, BIHD1, CGAC, CuRE, LTRE, MYB, MYC, SURE, TAAAG-BOX | AFR, ARR1, CATA, CPB, DPBF, NTBBF, PROX |
| *ZmTLP23* | BOX C, BOX II, BOX L, GATA, GT1CONSENSUS, I-box, INRN, PRE, SORLIP | CACT, CIACADIAN, DOFCORE, E2F, E-box, GCN, L1 box, POLLEN1, Q ELEMENT, | ASF1, BIHD1, CGAC, CuRE, EECC, ELRE, GCC, MYB, MYC, SEBF, WBOX, WRKY | ABRE, ARR1. CATA, CGCG-BOX-BOX, DPBF, |
| *ZmTLP24* | BOX II, GATA, GT1CONSENSUS, INRN, RBCS | ANAERO, CACT, DOFCORE, E-box, POLLEN1, RY elements, S1F, , SPH | ACGT, ASF1, BIHD1, CuRE, LTRE, MYB, MYC, QARB, T/G box, WBOX, WRKY | ABRE, ARR1, CATA, GADOWN |
| *ZmTLP25* | GATA, GT1CONSENSUS, PRE, SORLIP | ANAERO, CACT, CAN, CIACADIAN, DOFCORE, E-box, POLLEN1, RY elements, TGACG | ACGT, ASF1, CTR/DRE, CGAC, CuRE, GCC, LTRE, MYB, MYC, SURE, WRKY | AGC, ARF, ARR1, CGCG-BOX-BOX, CPB, DPBF, PROX |
| *ZmTLP26* | BOX L, GATA, GT1CONSENSUS, I-box, INRN, PRE, SORLIP | CACT, DOFCORE, E2F, E-box, L1 box, POLLEN1, Q ELEMENT, , , WUSAT | ACGT, ASF1, BIHD1, BP5O, EECC, ELRE, MYB, MYC, SEBF, SURE, T/G box, TAAAG-BOX, W BOX, WRKY | ABRE, AMYBOX, ARF, ARR1, DPBF, GADOWN, NTBBF, TATC |
| *ZmTLP27* | BOX L, GATA, INRN, PRE, SORLIP, T BOX | ANAERO, CACT, CIACADIAN, DOFCORE, E2F, E-box, POLLEN1, RY elements, S1F, TGAC | ACGT, ASF1, CTR/DRE, CGAC, CRTD, CuRE, EECC, ELRE, LTRE, MYC, SURE, WBOX, WRKY | ABRE, ARR1, CGCG-BOX-BOX, DPBF |
| *ZmTLP28* | GT1CONSENSUS, SORLIP, SV40 | ANAERO, CACT, DOFCORE, E-box, POLLEN1, RY elements, S1F, SPH | ACGT, BIHD1, CACG, CuRE, ELRE, MYB, MYC, SURE, WBOX, WRKY | ABRE, ARR1, CATA, CPBSC, DPBF, GARE, |
| *ZmTLP29* | GATA, GT1CONSENSUS, PRE, SORLIP, T-box | AACA, ANAERO, CACT, DOFCORE, E-box, POLLEN1, Q ELEMENT, RY elements, WUSAT | ACGT, ASF1, BIHD1, CTR/DRE, CuRE, DRE, EECC, LTRE, MYB, MYC, SURE, WBOX, WRKY | ABRE, ARR1, CATA |
| *ZmTLP30* | BOX L, GATA, GT1CONSESUS, I-box, PRE, SORLIP | CACT, DOFCORE, E-box, LEAF, POLLEN1, SRE, TGAC, TGTC, XYLAT | ACGT, ASF1, BIHD1, CGAC, CuRE, DRE, EECC, LTRE, MYB, MYC, P1BS, SEBF, TAAAG-BOX, WBOX, WRKY | ARR1, CGCG-BOX-BOX, CPB, GARE, NTBBF, TATC |
| *ZmTLP31* | BOX II, GATA, GT1CONSENSUS, I-box, SORLIP | ANAERO, CACT, DOFCORE, E2F, E-box, POLLEN1, Q ELEMENT, , SRE, WUSAT | ACGT, ASF1, BIHD1, CTR/DRE, CGAC, CuRE, DRE, EECC, ELRE, GCC, LTRE, MARABOX, MYB, MYC, SURE, TAAAG-BOX, W BOX, WRKY | ABRE, ARF, ARR1, DPBF |
| *ZmTLLP32* | BOX II, GATA, GT1CONSENSUS, I-box, | ANAERO, CACT, DOFCORE, E2F, E-box, POLLEN1, SEF1, , TGAC, UPR | ACGT, ASF1, BIHD1, CTR/DRE, CuRE, ELRE, MYB, MYC, SURE, T/G box, TAAAG-BOX, W BOX, WRKY | ABRE, ARF, ARR1, CATA, CGCG-BOX-BOX, GARE, |
| *ZmTLP33* | GATA, GT1CONSENSUS, I-box, PRE | ANAERO, DOFCORE, E2F, E-box, | ACGT, ASF1, CTR/DRE, CGAC, CRTD, CuRE, DRE, LTRE, MYB, MYC, UPR, WBOX, WRKY | ABRE, ARR1, CGCG-BOX-BOX |
| *ZmTLP34* | BOX II, BOX L, GATA, GT1CONSENSUS, I-box, PRE, SORLIP | CACT, CIACADIAN, DOFCORE, E2F, E-box, POLLEN1, S1F, , | ACGT, ASF1, BIHD1, CTR/DRE, CGAC, CuRE, DRE, EECC, GCC, LTRE, MARABOX, MYB, MYC, SEBF, SURE, WBOX, WRKY | ABRE, ARF, ARR1, CARE, CGCG-BOX-BOX, DPBF, GARE |
| *ZmTLP35* | BOX C, GATA, GT1CONSENSUS, PRE, SORLIP | ANAERO, CACT, CIACADIAN, DOFCORE, E2F, E-box, POLLEN1, S1F, | ACGT, ASF1, BIHD1, CuRE, EECC, GARE, LTRE, MYB, MYC, SURE, TAAAG-BOX, WBOX, WRKY | ABRE, ARR1, CACG, CARE, CGCG-BOX-BOX, DPBF, |
| *ZmTLP36* | BOX 1, GATA, GT1CONSENSUS, SORLIP, T BOX | ANAERO, CACT, DOFCORE, E2F, E-box, POLLEN1, UP1, XLAT | ACGT, ASF1, BIHD1, CGAC, CuRE, GCC, LTRE, MYB, MYC, TAAAG-BOX, UPR, WBOX, WRKY | ABRE, ARR1, CGCG-BOX-BOX, DPBF, NTBBF |
| *ZmTLP37* | BOX II, GATA, GT1CONSENSUS, I-box, PRE, SORLIP | AACA, ANAERO, CACT, DOFCORE, E2F, E-box, POLLEN1, RY elements | ACGT, ASF1, CTR/DRE, CuRE, DRE, EECC, LTRE, MYB, MYC, P1BS, SEBF, SURE, TAAAG-BOX, WBOX, WRKY | ABRE, ARF, ARR1, |
