## Supplementary figures and images for "Molecular characterization revealed the role of thaumatin-like proteins in stress response in bread wheat"

### Supplementary figure 1.pdf

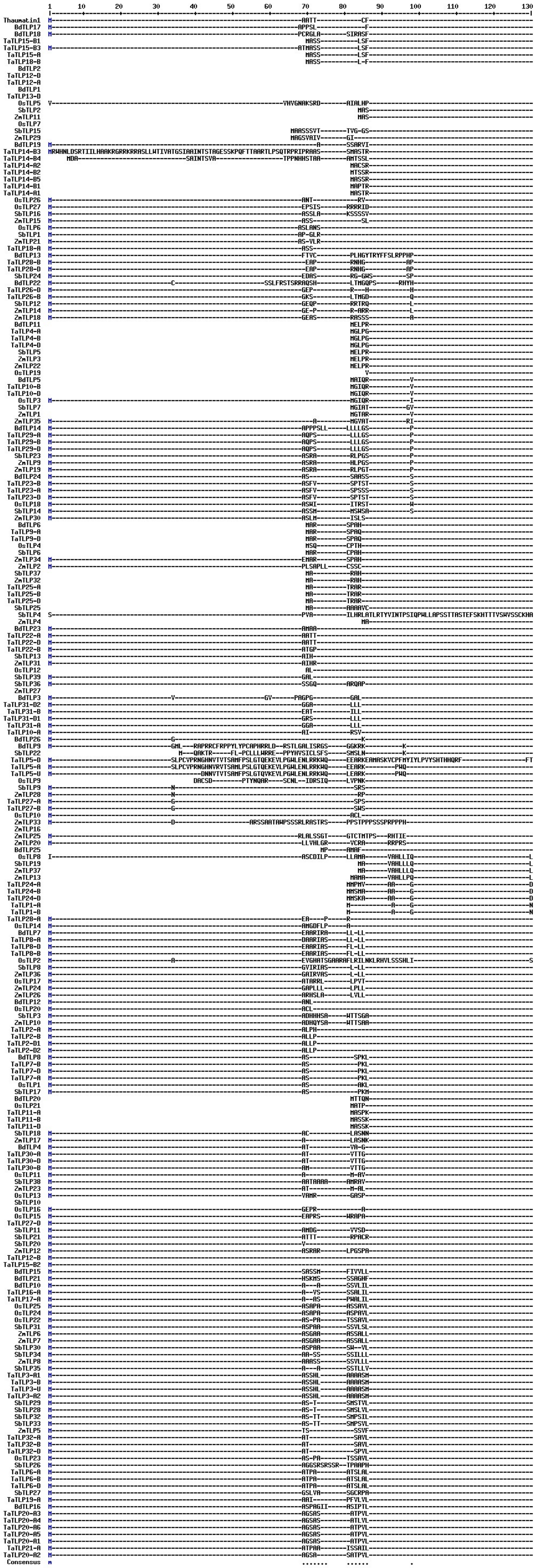

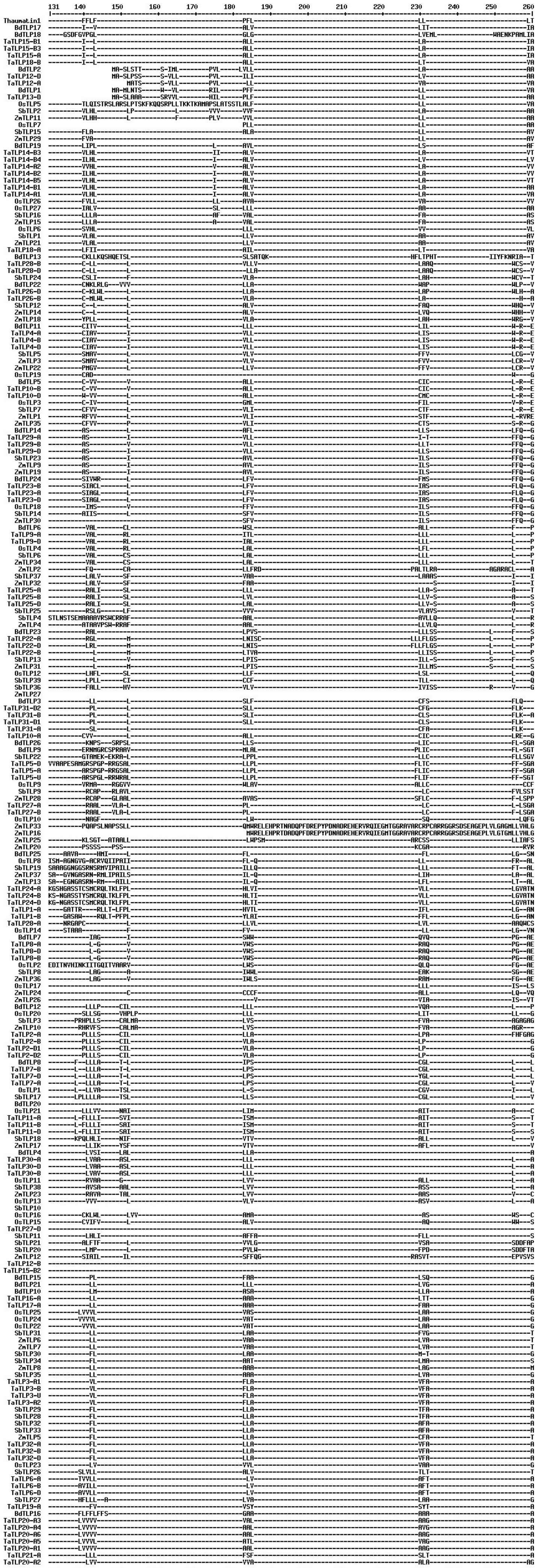

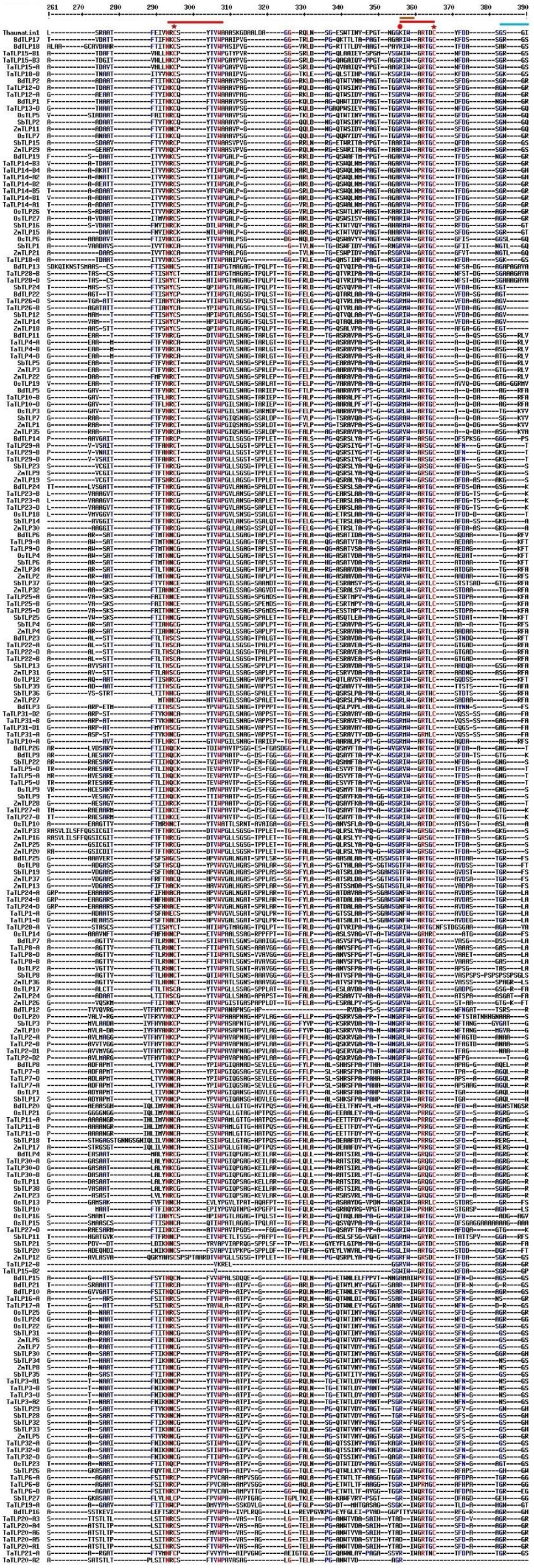

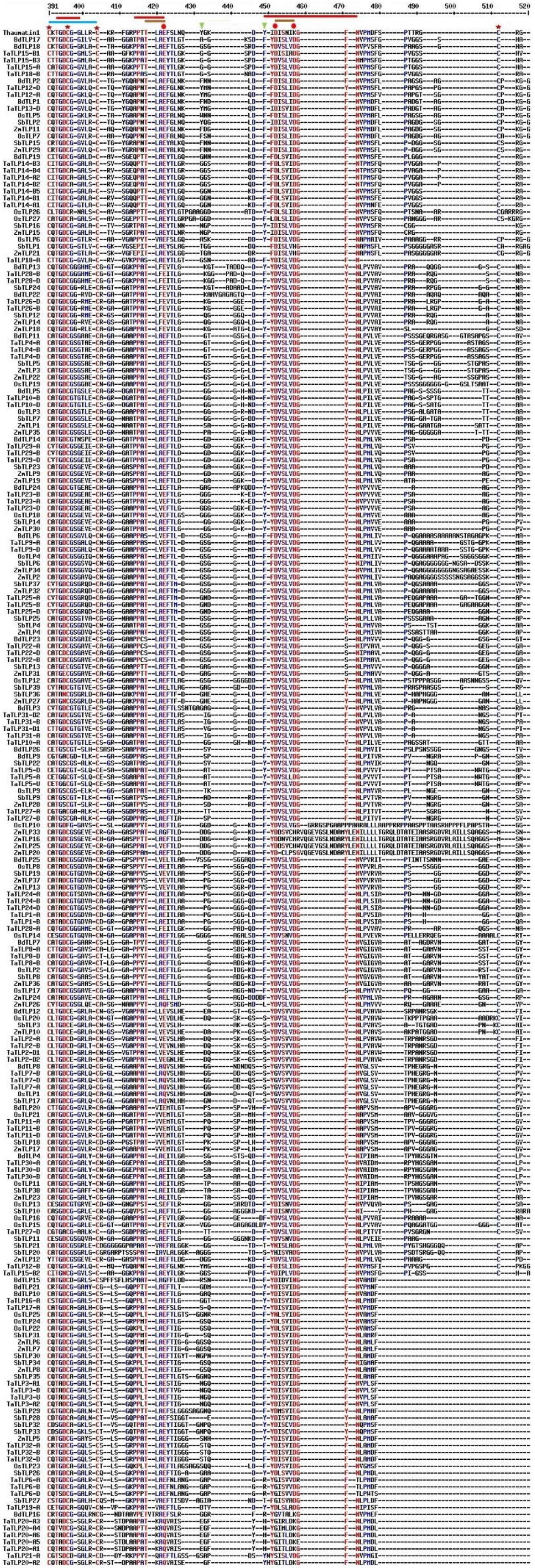

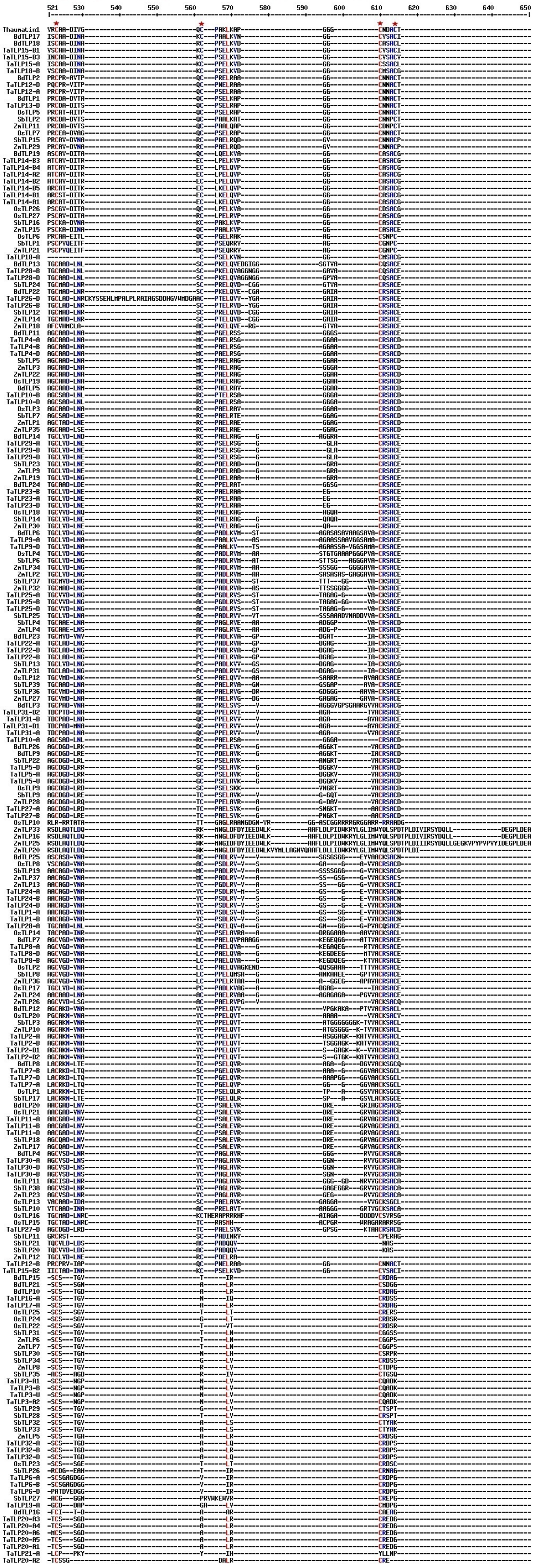

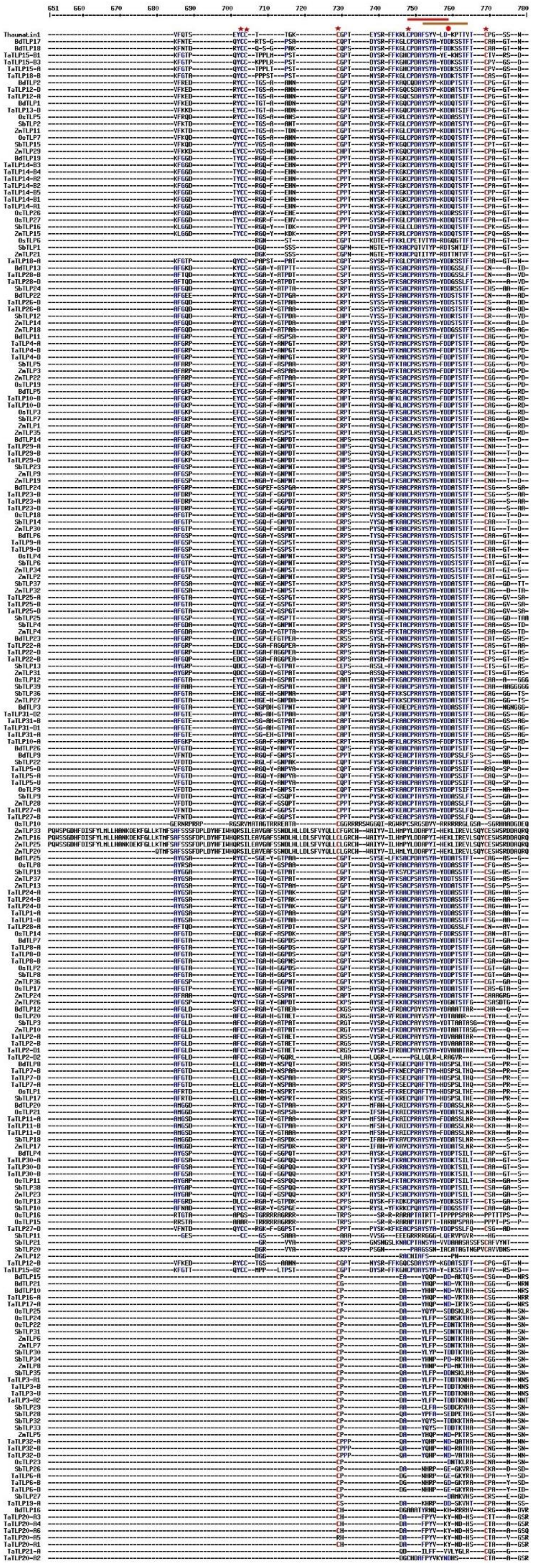

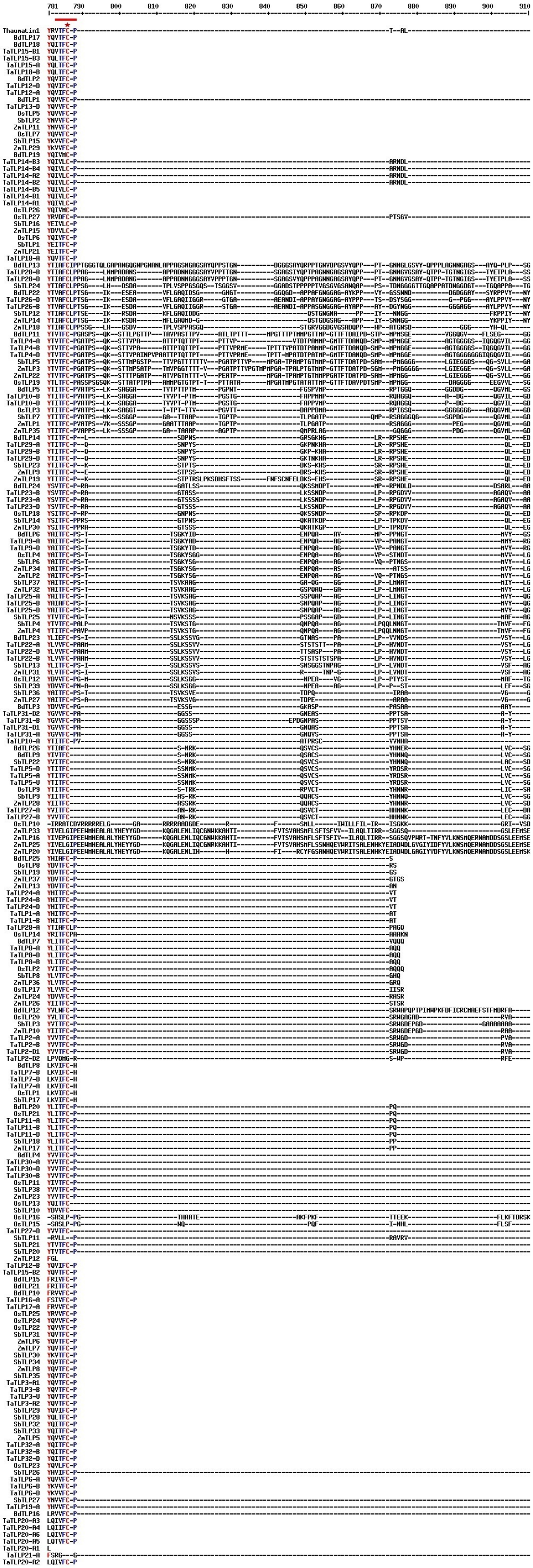

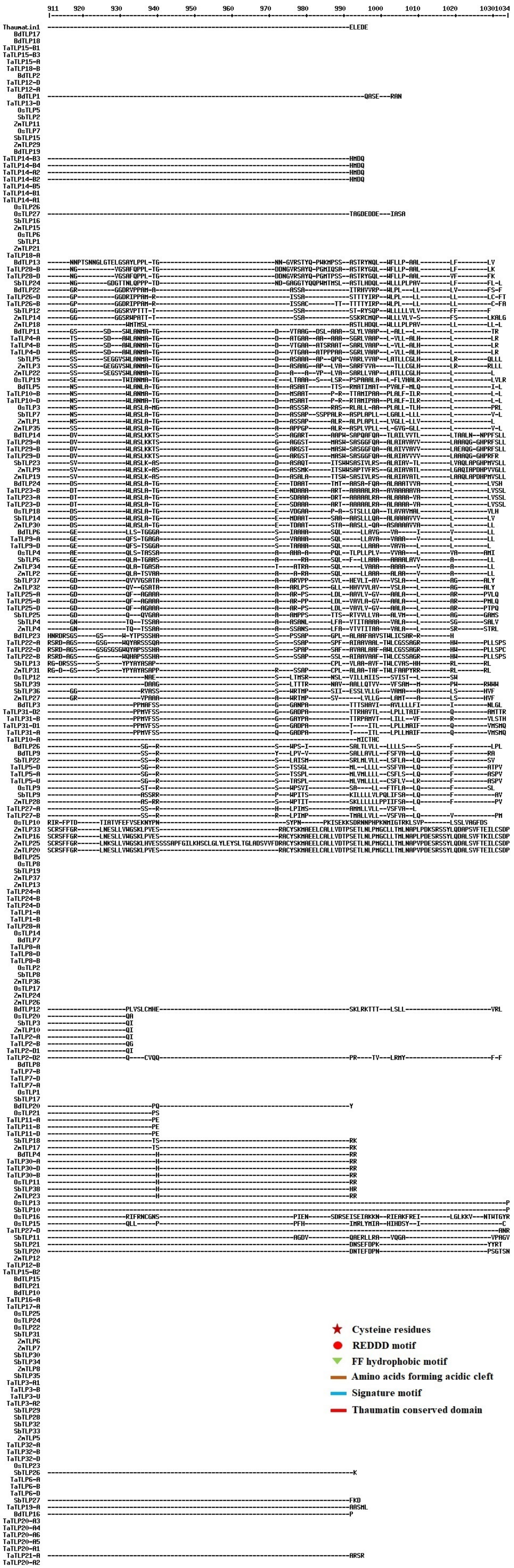

### Supplementary figure 2.pdf

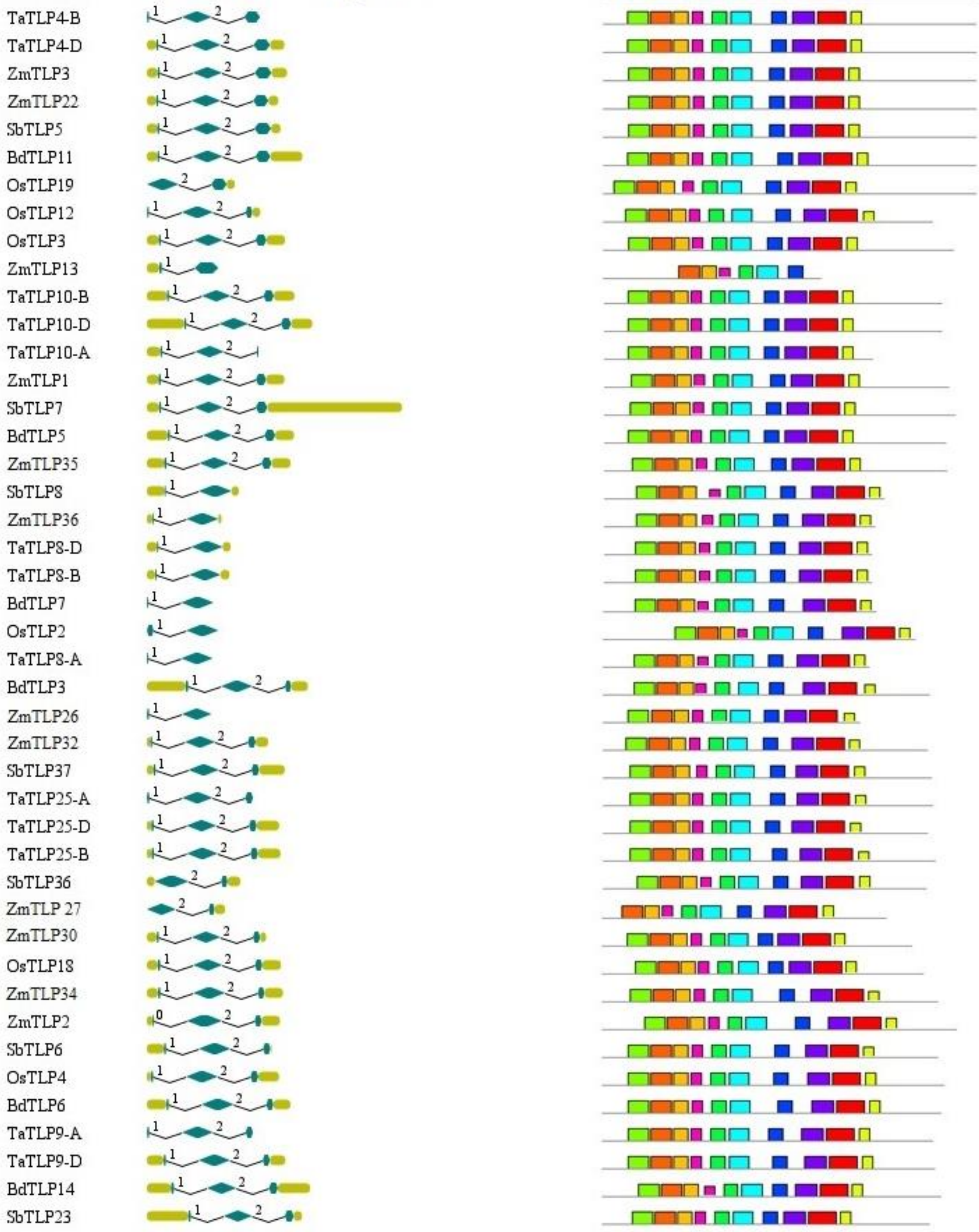

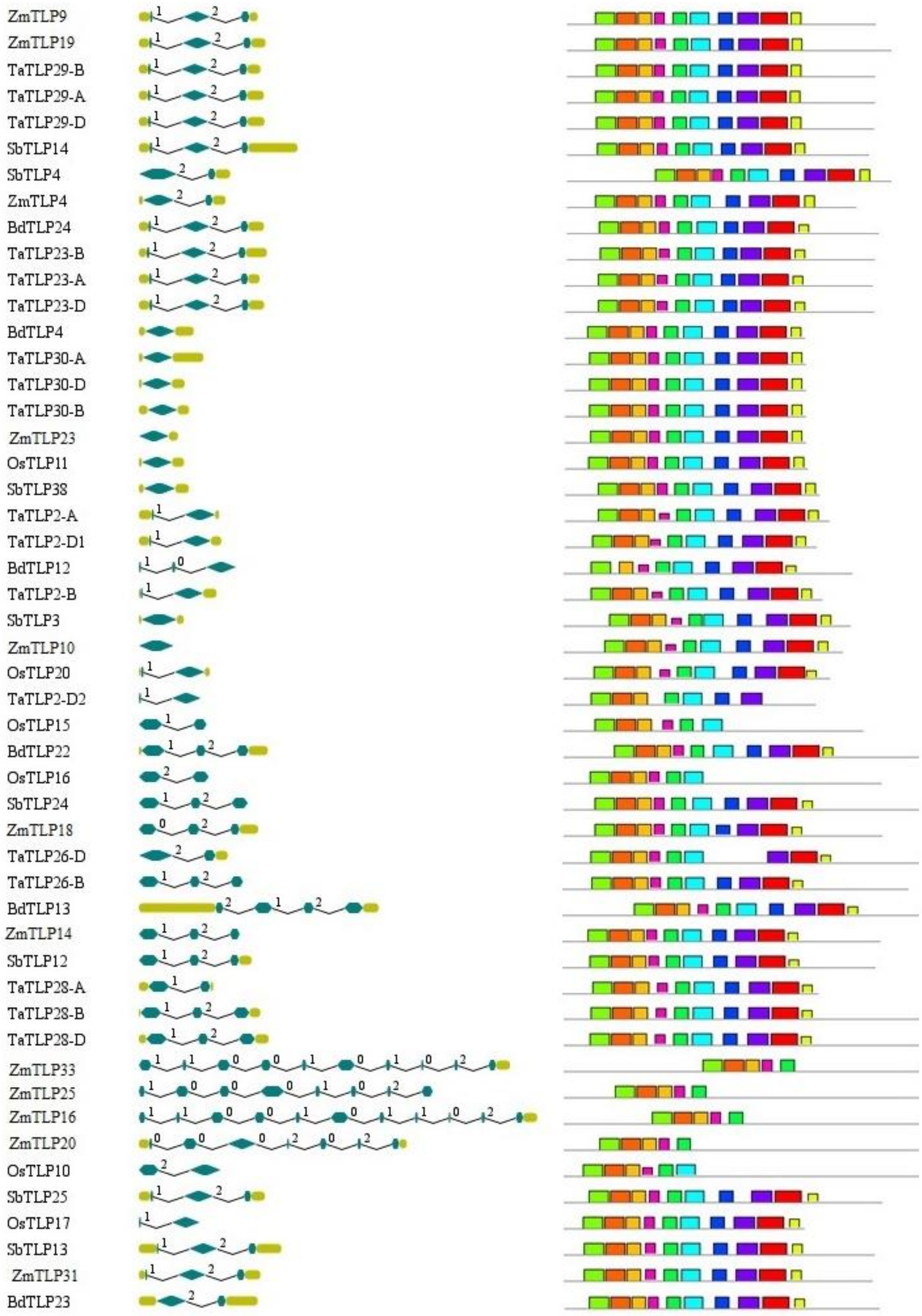

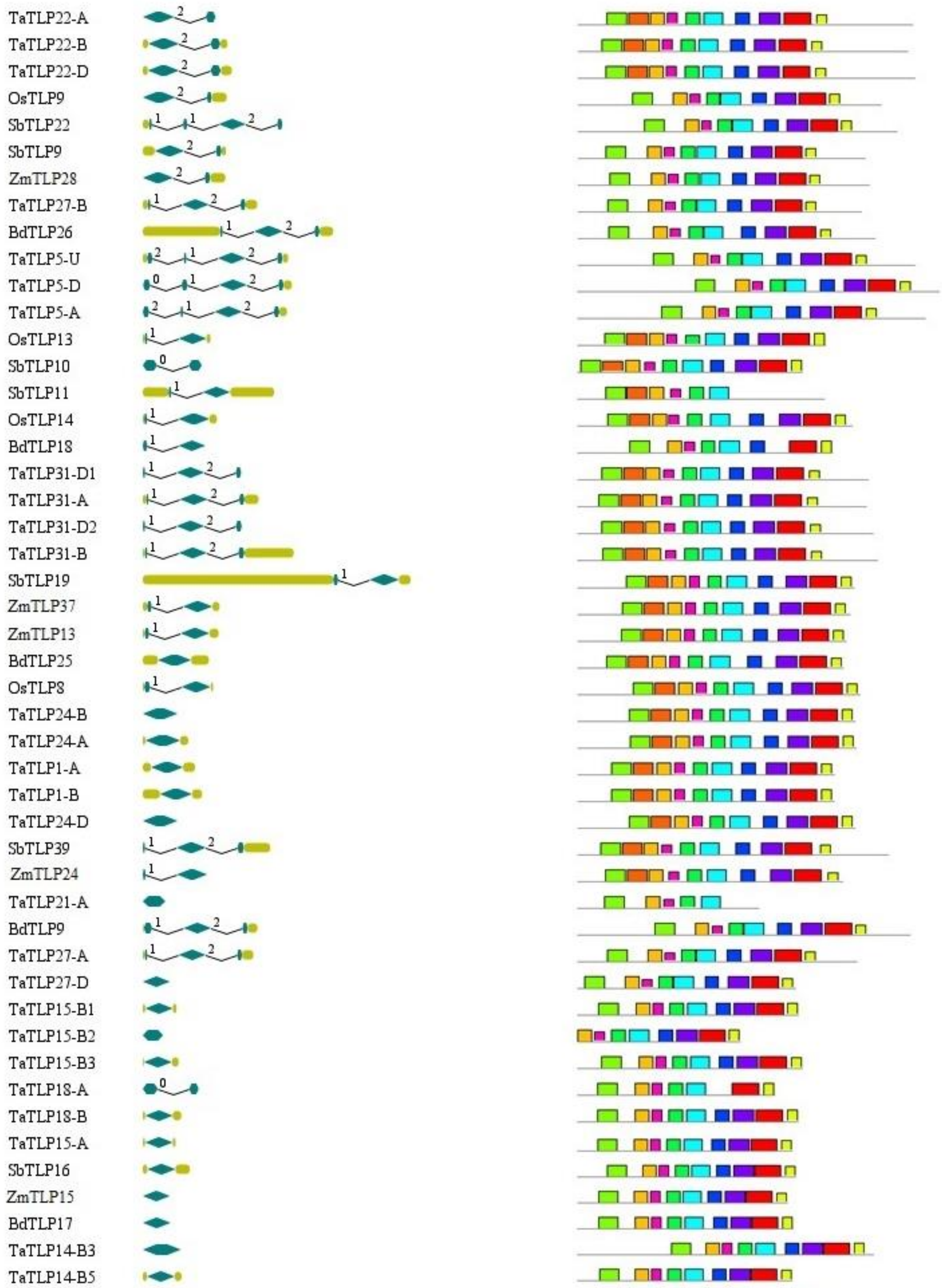

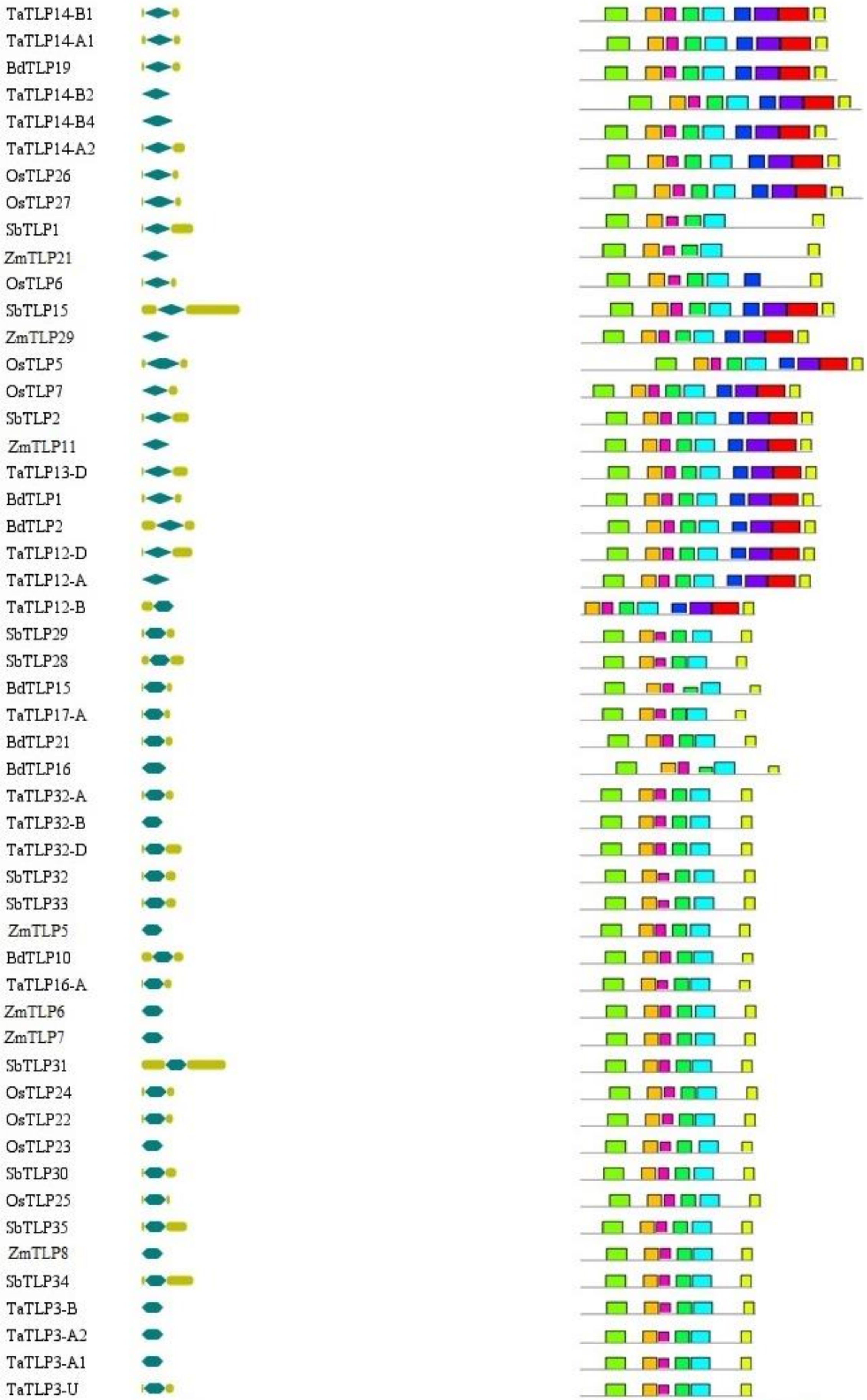

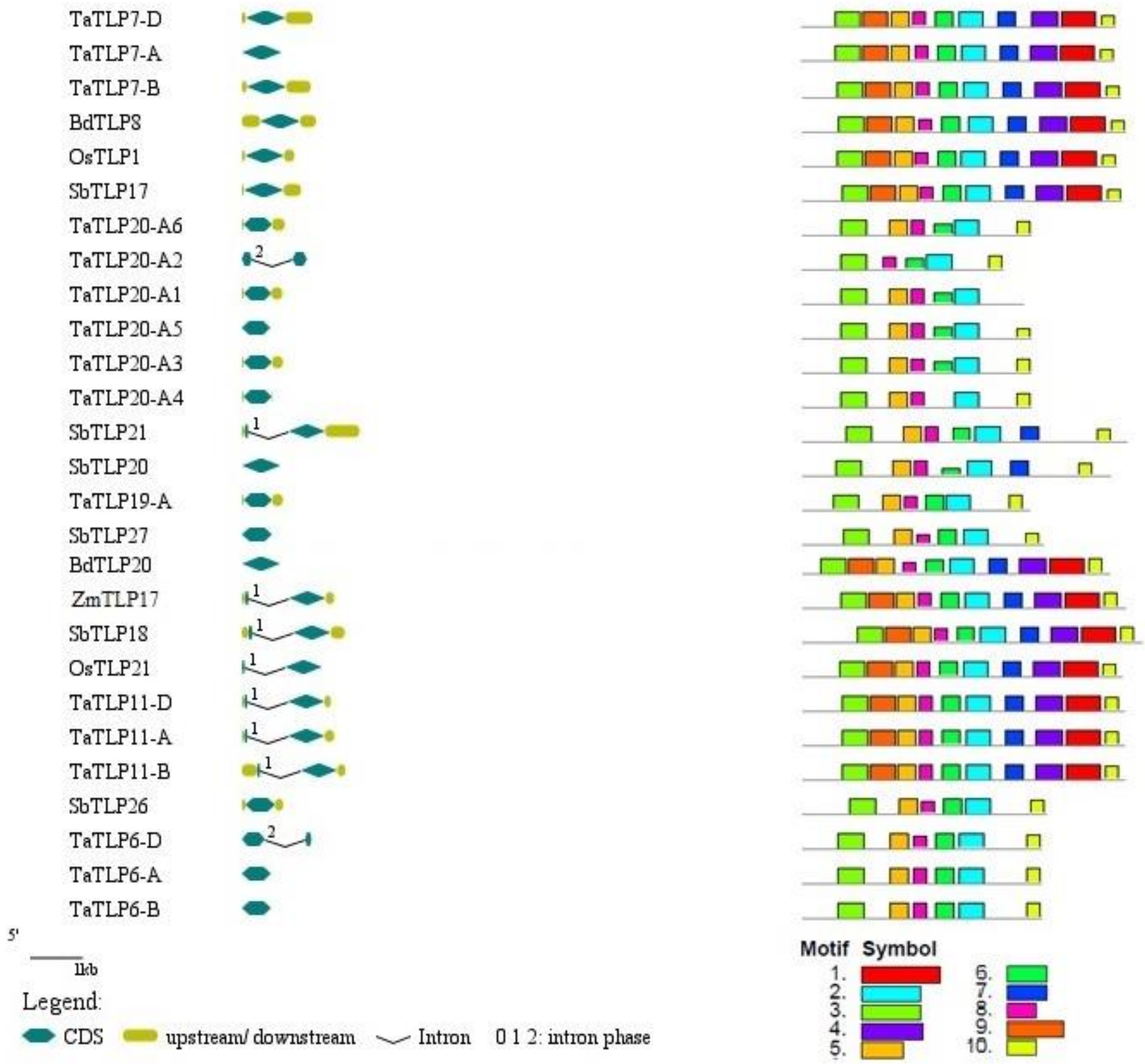
